# A transcriptional biosensor reveals mechanisms of α-ketoglutarate signaling to chromatin

**DOI:** 10.1101/2025.04.06.647450

**Authors:** Alex C. Sternisha, Haocheng Li, Jeffrey I. Traylor, Lei Guo, Ji Hyung Jun, Xin Zhao, Kumar Gajendra, Qing Ouyang, Michael Schmidt, Morgan Fleishman, Diana D. Shi, Milan R. Savani, Yi Xiao, Joyce H. Lee, Lauren G. Zacharias, Thomas P. Mathews, Ruth Gordillo, Yoon Jung Kim, Lin Xu, John G. Doench, Vidyasagar Koduri, Kalil G. Abdullah, Laura A. Banaszynski, Michalis Agathocleous, Ralph J. DeBerardinis, Eric M. Morrow, Samuel K. McBrayer

## Abstract

Alpha-ketoglutarate (αKG) is required for chromatin demethylation but mechanisms controlling αKG abundance in the nucleus are poorly defined. Therefore, we designed a biosensor system to monitor this metabolite pool in human cells using an αKG-responsive cyanobacterial transcription factor, NtcA. We then coupled this system with a genetic screen to identify genes that regulate αKG in the nucleus, defining an inter-organelle pathway in which sequential mitochondrial activities of the GPT2 transaminase and SLC25A11 transporter supply nuclear αKG. Using a mouse model of GPT2 deficiency, a human inborn error of metabolism, we found that this pathway controls chromatin methylation in the developing brain. Our work provides a tool to assess αKG signaling to chromatin and a framework for leveraging forward genetics to study nuclear metabolite pools.

## Main Text

Intracellular biochemical pathways are intimately tied to epigenetic regulation, as metabolites serve as substrates for “writer” and “eraser” enzymes that chemically modify DNA and histones (*1*, *2*). Nuclear pools of acetyl-CoA and S-Adenosylmethionine (SAM) are required for chromatin acetylation and methylation, respectively. Conversely, NAD+ and alpha- ketoglutarate (αKG) promote chromatin deacetylation and demethylation, respectively. These metabolites mediate crosstalk between the metabolome and epigenome, ultimately regulating gene expression programs (*3*, *4*). αKG in particular plays a key role in cell fate control and tumor suppression, as dysregulation of αKG availability in the nucleus alters cell state transitions (*5–9*). Moreover, recurrent cancer-associated mutations in the metabolic enzymes isocitrate dehydrogenase (IDH), succinate dehydrogenase (SDH), or fumarate hydratase (FH) produce oncometabolites that directly interfere with αKG signaling to chromatin, establishing chromatin hypermethylation states that prime cells for malignant transformation (*10–18*). Despite this important regulatory role for the nuclear αKG pool, we have limited insights into the molecular mechanisms that control it. This lack of understanding is tied to a paucity of experimental tools to measure nucleus-specific αKG in mammalian cells, as well as the vast number of enzymes and transporters that act on αKG.

We pursued a synthetic biology approach to monitor the nuclear αKG pool in living human cells. Our approach was informed by an αKG sensing mechanism that is conserved across cyanobacteria species. Cyanobacteria assimilate nitrogen by scavenging ammonium and using it to synthesize glutamine and glutamate from αKG through glutamate dehydrogenase, glutamine oxoglutarate aminotransferase, and glutamine synthetase enzymes (*19*). Under nitrogen starvation, αKG accumulates and is sensed by the transcription factor NtcA (*20*). αKG allosterically increases the affinity of NtcA homodimers for DNA (*21*), enabling them to recognize cognate binding sites throughout the genome and activate a gene expression program that promotes adaptation to nitrogen deprivation (*22*). Given prior evidence that NtcA transcription factors recognize αKG when expressed ectopically (*23*), we hypothesized that an engineered, chimeric NtcA transcription factor could be used to drive expression of a synthetic reporter gene that serves as a proxy for nuclear αKG pool size in human cells. Further, we hypothesized that such a system may allow us to apply unbiased, forward genetic approaches to identify genes that control αKG abundance in the nucleus. Although metabolite pools have been used as endpoints for genetic screens in prior studies, this work has largely focused on optimizing biosynthetic reactions in microbes. In contrast, we sought to marry the power of genetic screening with a nuclear αKG reporter to reveal mechanisms of compartmentalized metabolism that control αKG-dependent chromatin demethylation in human cells.

## Results

### Designing and optimizing the αKG-ON biosensor system to monitor nuclear αKG

To address the paucity of tools to measure nuclear αKG in human cells, we drew inspiration from two sources: the Tet-ON system for inducible gene expression (*24*, *25*) and a cyanobacterial gene expression program controlled by an αKG-regulated transcription factor, NtcA (*26*). While the Tet-ON system places transgene activation under the control of exogenously supplied tetracyclines, we hypothesized that we could leverage design principles of this system to develop a biosensor that places expression of a reporter gene under the control of the endogenous nuclear αKG pool. We sought to create two components comprising a nuclear αKG biosensor (hereafter, the αKG-ON biosensor system): a chimeric NtcA transcription factor and a promoter/reporter gene DNA element that can be transactivated by NtcA (**Fig. 1A**). We posited that fusing a cyanobacterial NtcA protein to nuclear localization signal and herpesvirus- derived VP64 transactivation domain peptides would allow us to direct NtcA chimeras to the nucleus and drive transcription in human cells. Further, we proposed that engineering a synthetic promoter with NtcA binding sites sourced from a cyanobacterial genome (hereafter, the αKG response element or αKG-RE) and placing it upstream of a GFP reporter gene would allow us to monitor changes in nuclear αKG content in living human cells. This design schema is based on biochemical (*21*) and structural (*20*, *27*) studies establishing that αKG binding to the effector binding domains (EBDs) of NtcA homodimers triggers allosteric alignment of DNA binding domains (DBDs) of these transcription factors with successive DNA major grooves (**Fig. 1B**) .

**Fig. 1.**
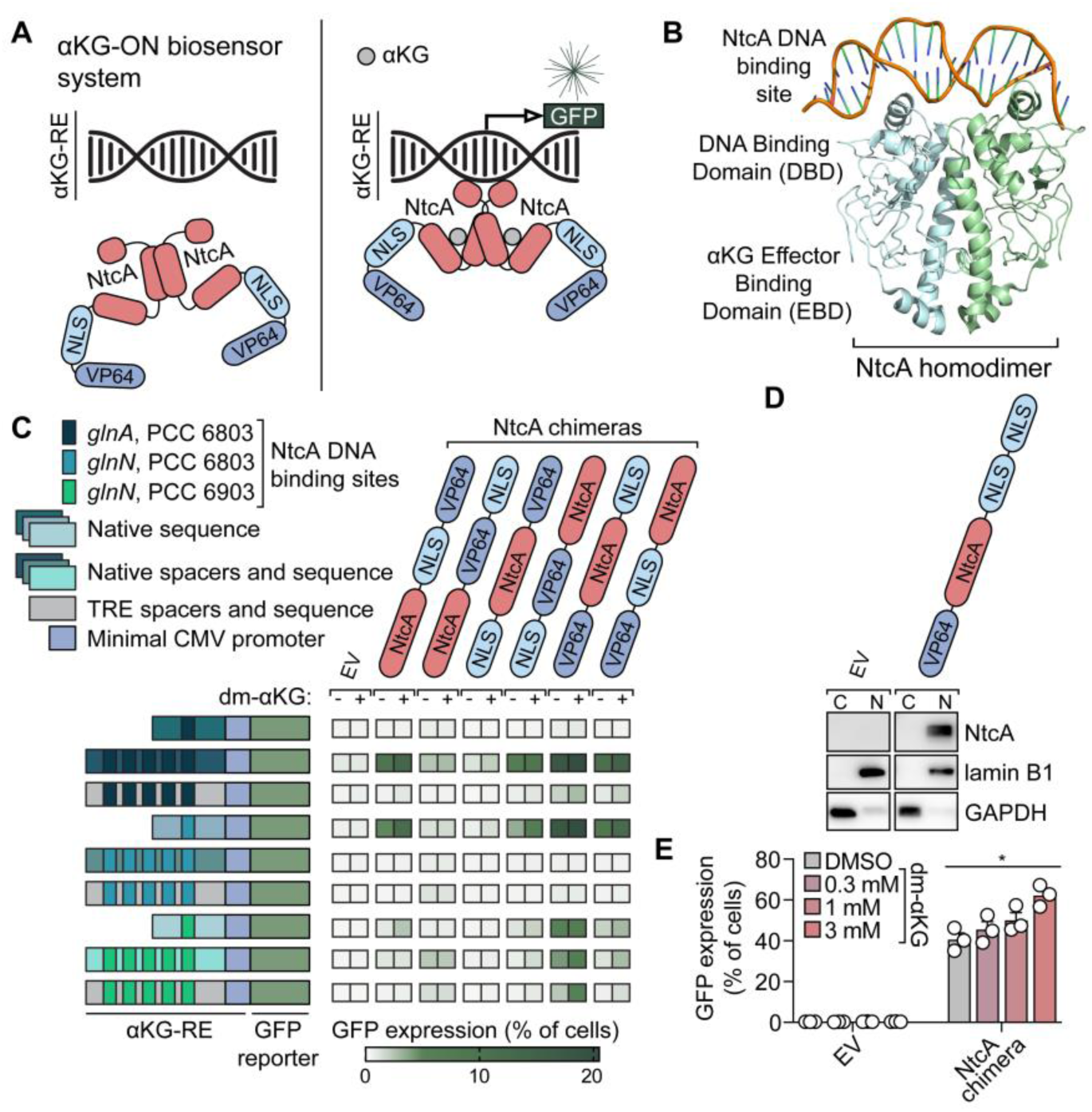
Development and optimization of the αKG-ON biosensor system to monitor the nuclear αKG pool. (**A**) Schema of αKG-ON biosensor system transcriptional activity in the presence and absence of αKG. αKG-RE = αKG response element; NLS = nuclear localization signal; VP64 = tetramer of Herpes Simplex Virus 1 (HSV-1) VP16 transactivation domains; GFP = green fluorescent protein. (**B**) Cryo-electron microscopy structure of NtcA homodimer bound to DNA. Data are derived from Protein Data Bank accession 8H40, reference (*27*). (**C**) Pairwise combinatorial screen of αKG-ON biosensor system architectures in HEK293 cells. HEK293 lines stably expressing αKG-RE elements were transiently transfected with NtcA chimeras or an empty vector (EV, negative control). αKG-ON biosensor system activation (GFP expression) was quantified by flow cytometry in cells treated with 1 mM dimethyl-αKG (dm-αKG) or DMSO for 48 hours prior to analysis. (**D**) Immunoblot of the VP64-NtcA-2xNLS_SV40_ chimera in nuclear (N, lamin B1 marker) and cytosolic (C, GAPDH marker) fractions prepared from HEK293 cells engineered to express the NtcA chimera or EV. (**E**) Flow cytometry quantification of αKG-ON biosensor system activation (GFP expression) in αKG-RE-expressing HEK293 cells transduced with the VP64-NtcA-2xNLS_SV40_ chimera or EV and treated with DMSO or the indicated doses of dm-αKG for 24 hours. Data are means ± SEM. \**p* < 0.05 (ordinary one-way ANOVA).

We first tested the αKG-ON biosensor system concept by screening varied αKG-RE and NtcA chimera architectures in a pairwise manner in HEK293 cells (**Fig. 1C and fig. S1A**). To create synthetic αKG-RE promoters, we derived NtcA binding sites from the promoters of glutamine synthetase*-*encoding *glnA* or *glnN* genes in *Synechocystis* sp. PCC 6803 or *Pseudanabaena* sp. PCC 6903 genomes (*22*, *28*, *29*) (**fig. S1B-C**). We incorporated these binding sites either individually or in 5-repeat tiles flanked by native cyanobacterial genomic sequences or sequences derived from the tetracycline response element (TRE). NtcA binding sites were placed upstream of a minimal CMV promoter. To assess outputs of biosensor system architectures under basal and αKG-stimulated conditions, we introduced biosensor components into cells, treated them with or without cell-permeable dimethyl-αKG (dm-αKG), and measured GFP expression by flow cytometry. Several combinations of NtcA chimeras and αKG-RE promoters displayed basal GFP reporter transactivation (presumably driven by endogenous nuclear αKG) that was enhanced by dm-αKG supplementation. Importantly, dm-αKG did not induce GFP in empty vector-expressing cells, indicating that αKG-ON biosensor system output requires a chimeric NtcA transcription factor. The αKG-RE promoter containing tiled NtcA binding sites from the PCC 6903 *glnN* promoter with TRE-derived spacers produced a marked difference in GFP expression between basal and αKG-stimulated conditions relative to other synthetic promoters. Therefore, this αKG-RE was selected for further study in HEK293 cells.

We noted that only a portion of cells mounted a GFP expression response upon NtcA chimera transfection and dm-αKG treatment. Therefore, we sought to optimize αKG-ON biosensor system performance by improving the function of the NtcA chimeric transcription factor. Cell fractionation experiments showed that NtcA chimeras were not exclusively nuclear and that some chimera expression constructs produced truncated protein products (**fig. S1D**). In silico analyses implicated cryptic nuclear export sequences in the VP64 peptide as a potential cause of cytosolic accumulation of NtcA chimeras (**fig. S1E**). Eliminating VP16 monomer peptides that comprise the VP64 transactivation domain in these fusion proteins dose-dependently enhanced nuclear localization but reduced biosensor output (**fig. S1F-G**). We replaced VP64 with other mammalian transactivation domains (*30–33*) and found that the Epstein-Barr virus (EBV)- derived Zta protein enhanced nuclear targeting of the NtcA chimera and eliminated truncated protein products (**fig. S1H**). A similar result was produced by removing an unnecessary methionine residue from the N-terminal portion of NtcA (to prevent alternative translation) and replacing the c-myc-derived NLS with two NLS peptides from SV40 large T-antigen in the VP64-NtcA-NLS chimera (**fig. S1I**). The VP64-NtcA-2xNLS_SV40_ transcription factor displayed superior nuclear localization and αKG-RE promoter activation relative to the Zta-NtcA-NLS_c-myc_ fusion (**Fig. 1D-E** and **fig. S1I-J**). Further, the dynamic range of biosensor system output driven by VP64-NtcA-2xNLS_SV40_ was greater than or equal to that of Zta-NtcA-NLS_c-myc_ when these chimeras were expressed under promoters of varying strength (i.e. UBC, PGK, or EF1α) (**fig. S2A-R**). Based on these findings, an EF1α promoter-driven VP64-NtcA-2xNLS_SV40_ chimera (or mutants thereof) was used in all subsequent studies.

### The αKG-ON biosensor system responds to compartment-specific changes in αKG abundance

To interrogate the specificity of the optimized αKG-ON biosensor system, we first created 6 DMEM medium preparations that produce a gradient of intracellular αKG levels in HEK293 cells (**fig. S3A**). In these medium stocks, we reduced levels of glutamine or introduced dm-αKG. Glutamine is catabolized to αKG predominantly through sequential activities of glutaminase/glutamine amidotransferase and glutamate dehydrogenase/transaminase enzymes.

Next, we created mutants of the VP64-NtcA-2xNLS_SV40_ chimera that lack αKG (R90E) (*20*, *23*) or DNA (ΔDBD) binding functions. We transduced HEK293 cells stably expressing an αKG-RE promoter/GFP reporter with either of these mutant NtcA chimeras, a wildtype (WT) NtcA chimera, or an empty vector control. Culturing the resulting stable cell lines in custom DMEM medium stocks selectively induced a gradient of GFP expression in cells engineered with the WT NtcA chimera (**Fig. 2A-H**). Importantly, GFP expression in these cells correlated linearly with cellular αKG levels over a ∼24-fold range, capturing both increases and decreases in this metabolite pool relative to standard culture conditions (i.e. 2 mM glutamine without dm-αKG). We hypothesized that the dynamic range of the GFP response could be expanded by fusing GFP to an hPEST degron to increase reporter turnover (*34*, *35*), although this proved ineffective (**fig. S3B-I**). In cell lines expressing the WT NtcA chimera or the αKG binding-deficient R90E mutant NtcA chimera (*20*, *23*) (hereafter, “functional” and “inactive” versions of the αKG-ON biosensor system, respectively), GFP expression was selectively reduced in the former line by glutamine restriction (**Fig. 2I** and **fig. S4A-B**). This effect was fully rescued by dm-αKG supplementation. Therefore, glutamine deprivation decreases functional biosensor output in an αKG-dependent manner. GFP expression was not affected by (*R*)-2-hydroxyglutarate- Bis(trifluoromethyl)benzidine [(*R*)-2HG-TFMB], a cell-permeable ester of the oncometabolite (*R*)-2HG (*36*), in either functional or inactive biosensor-expressing cells (**fig. S4C-G**). Given the structural similarity of (*R*)-2HG and αKG, our findings indicate that biosensor output is not confounded by moderate changes in chemical congeners of αKG. Collectively, these data establish that output of the αKG-ON biosensor system is specifically regulated by intracellular αKG levels and requires both the αKG and DNA binding activities of the NtcA transcription factor.

**Fig. 2.**
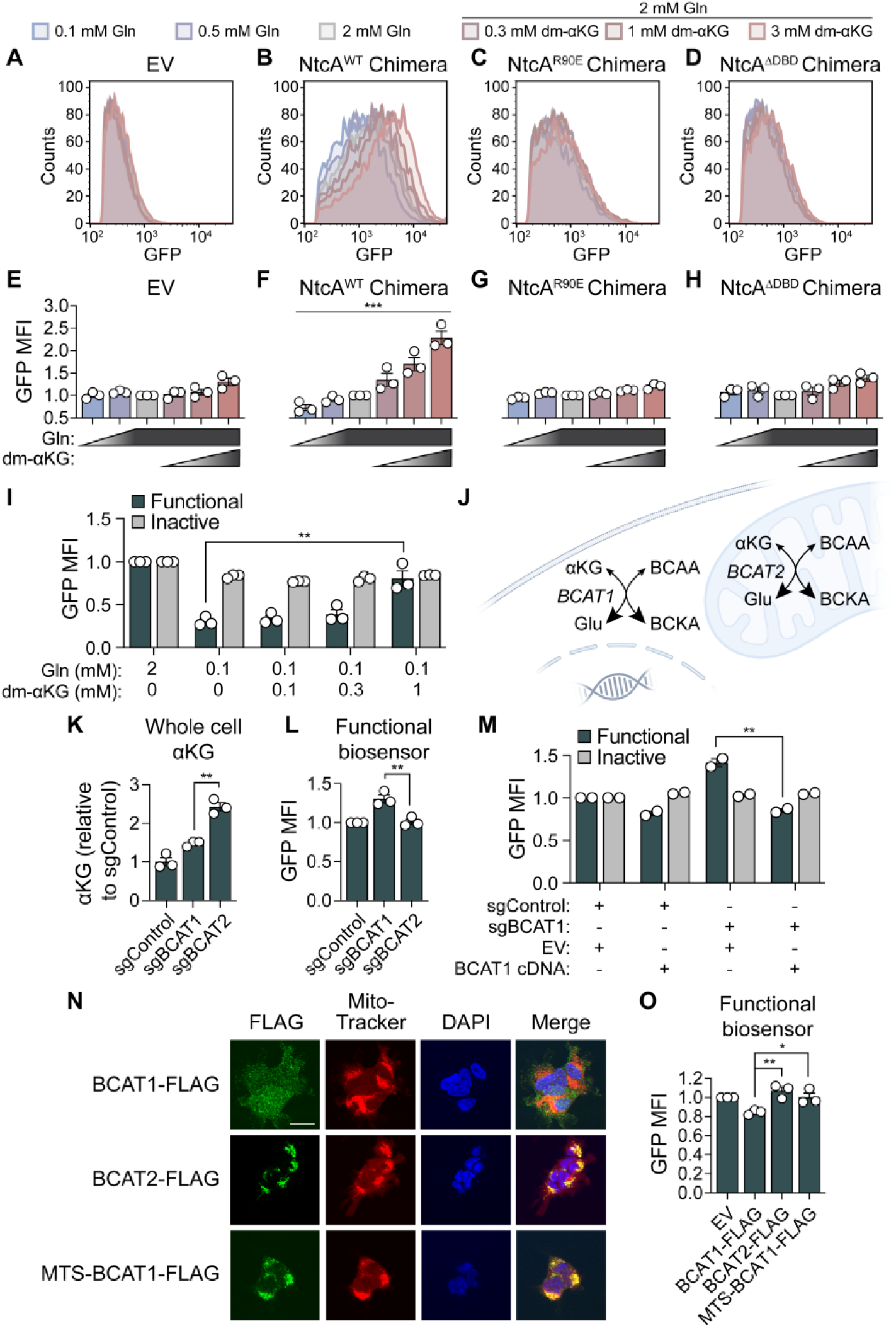
The αKG-ON biosensor system responds to metabolic and genetic perturbations of the nuclear αKG pool. (**A-H**) Representative histograms (A-D) and quantification (E-H) of GFP expression in αKG-RE-expressing HEK293 cells transduced with EV or one of three VP64- NtcA-2xNLS_SV40_ chimeras: wildtype (WT) NtcA, R90E NtcA mutant, DNA binding domain (DBD)-deleted NtcA mutant. Cells were cultured under the indicated concentrations of glutamine and/or dm-αKG for 72 hours prior to analysis. In (E-H), data are GFP mean fluorescence intensity (MFI) values normalized to 2 mM Gln condition. \*\*\**p* < 0.001 (ordinary one-way ANOVA). (**I**) Quantification of GFP expression (GFP MFI normalized to 2 mM Gln condition) in HEK293 cells engineered with functional (NtcA^WT^) or inactive (NtcA^R90E^) versions of the αKG-ON biosensor system and cultured with indicated concentrations of Gln and/or dm-αKG for 72 hours. \*\**p* < 0.01 (unpaired t-test). (**J**) Schema of branched chain amino acid (BCAA) metabolism by BCAT1 and BCAT2 transaminases in the nucleocytosolic and mitochondrial compartments, respectively. BCKA = branched chain α-ketoacid. (**K**) Relative αKG levels in whole-cell HEK293 extracts. HEK293 cells were engineered to express Cas9 and control, BCAT1, or BCAT2 sgRNAs. \*\**p* < 0.01 (unpaired t-test). (**L**) Relative nuclear αKG levels (GFP MFI normalized to sgControl line) in cells from (K) engineered with the functional αKG-ON biosensor system. \*\**p* < 0.01 (unpaired t-test). (**M**) Quantification of GFP expression (GFP MFI normalized to sgControl+EV lines) in HEK293 cells engineered with functional or inactive versions of the αKG-ON biosensor system. Cells were transduced with Cas9 and control or BCAT1 sgRNAs, as well as an sgRNA-resistant BCAT1 cDNA or EV. \*\**p* < 0.01 (unpaired t-test). (**N**) Representative immunofluorescence microscopy images of FLAG tag expression and Mito-Tracker or DAPI stains from HEK293 cells engineered to express FLAG-tagged BCAT1^WT^ enzyme, BCAT2^WT^ enzyme, or a BCAT1 mutant enzyme with an N-terminal mitochondrial targeting signal (MTS) peptide. Merge shows overlay of all signals. Scale bar = 10 μm. (**O**) Relative nuclear αKG levels (GFP MFI normalized to EV line) in cells from (N) or cells expressing an EV. All lines were also engineered with the functional αKG-ON biosensor system. \**p* < 0.05, \*\**p* < 0.01 (unpaired t-tests). In (E-I, K-M, and O), data are means ± SEM.

Although the optimized VP64-NtcA-2xNLS_SV40_ chimera localizes exclusively to the nucleus (**Fig. 1D** and **fig. S1I**), we had not yet evaluated whether the αKG-ON biosensor system responds to compartment-specific changes in αKG abundance. Further, it was not clear if the biosensor could detect genetic perturbations of the nuclear αKG pool in addition to perturbations caused by changes in nutrient availability. αKG exchange across the inner mitochondrial membrane is controlled by dedicated transport mechanisms, thereby partitioning mitochondrial αKG metabolism from that in other organelles. In contrast, the nuclear and cytosolic αKG pools are thought to equilibrate owing to the flow of metabolites and other small molecules through nuclear pores (*37*, *38*). Consistent with this idea, increased activity of the nucleocytosolic (BCAT1), but not mitochondrial (BCAT2), branched chain amino acid transaminase has been shown to cause DNA hypermethylation in leukemia cells (*39*) (**Fig. 2J**). This interaction was attributed to BCAT1-dependent depletion of nucleocytosolic αKG, which is a substrate for ten-eleven translocation (TET) methylcytosine dioxygenases that catalyze the first step in 5-methylcytosine demethylation. We exploited this difference between BCAT paralogs to assess the ability of the αKG-ON biosensor system to monitor compartmentalized αKG metabolism. BCAT2 knockout caused a larger increase in whole cell αKG levels relative to knockout of BCAT1 (**Fig. 2K** and **fig. S5A**). Conversely, BCAT1, but not BCAT2, knockout elevated nuclear αKG levels, as indicated by functional biosensor output (**Fig. 2L**). This effect was on-target because the increase in nuclear αKG pool size caused by BCAT1 knockout was rescued by sgRNA-insensitive BCAT1 cDNA expression (**Fig. 2M** and **fig. S5B-C**). Moreover, BCAT1 knockout did not affect output of the inactive αKG-ON biosensor system. BCAT1 and BCAT2 paralogs have different properties that extend beyond subcellular localization. To ask if differences in localization account for selective regulation of biosensor system output by BCAT1, we engineered cells to express FLAG-tagged WT BCAT1, WT BCAT2, or a BCAT1 mutant created by appending an N-terminal mitochondrial targeting sequence (MTS) (**fig. S5D**). We confirmed that WT BCAT1 localized to the nucleocytosolic compartment whereas WT BCAT2 and MTS-BCAT1 enzymes were mitochondrial (**Fig. 2N**). Overexpression of WT BCAT1 reduced nuclear αKG levels while WT BCAT2 and MTS-BCAT1 did not (**Fig. 2O**).

Therefore, the αKG-ON biosensor system senses compartment-specific changes in αKG abundance evoked by genetic manipulation of endogenous metabolic enzyme function.

### A forward genetic approach identifies genes that regulate the nuclear αKG pool

Development of the αKG-ON biosensor system provided us with a new tool to address a fundamental biological question: what are the key molecular regulators of the pool of αKG in the nucleus that supports chromatin demethylation? We designed an unbiased forward genetic screen to answer this question (**Fig. 3A**). We first stably expressed functional or inactive versions of the αKG-ON biosensor system in U251 differentiated glioblastoma cells. We selected this line due to its excellent performance in high-throughput genetic screens (*40*, *41*). We engineered U251 cells with an αKG-RE promoter (tiled NtcA binding sites from the *Synechocystis* sp. PCC 6803 *glnA* promoter with native spacer sequences) that drove robust NtcA-dependent GFP expression in this line and in HEK293 cells (**Fig. 1C**). Next, we sequentially transduced these lines to express Cas9 and a custom “αKG Regulators” CRISPR knockout sgRNA library. This library targets 127 genes encoding proteins that directly synthesize, catabolize, or transport αKG. Cells displaying GFP expression in the top and bottom 10 percentiles of each arm were then isolated via fluorescence-activated cell sorting (FACS). Finally, we quantified changes in sgRNA abundance between the top 10% and bottom 10% of GFP-expressing cell populations, prioritizing sgRNAs that were selectively enriched or depleted in functional biosensor-expressing cells for further study. This approach enhanced our ability to distinguish “true positives” from “false positives”, or those sgRNAs that affect biosensor output in an αKG- independent manner.

**Fig. 3.**
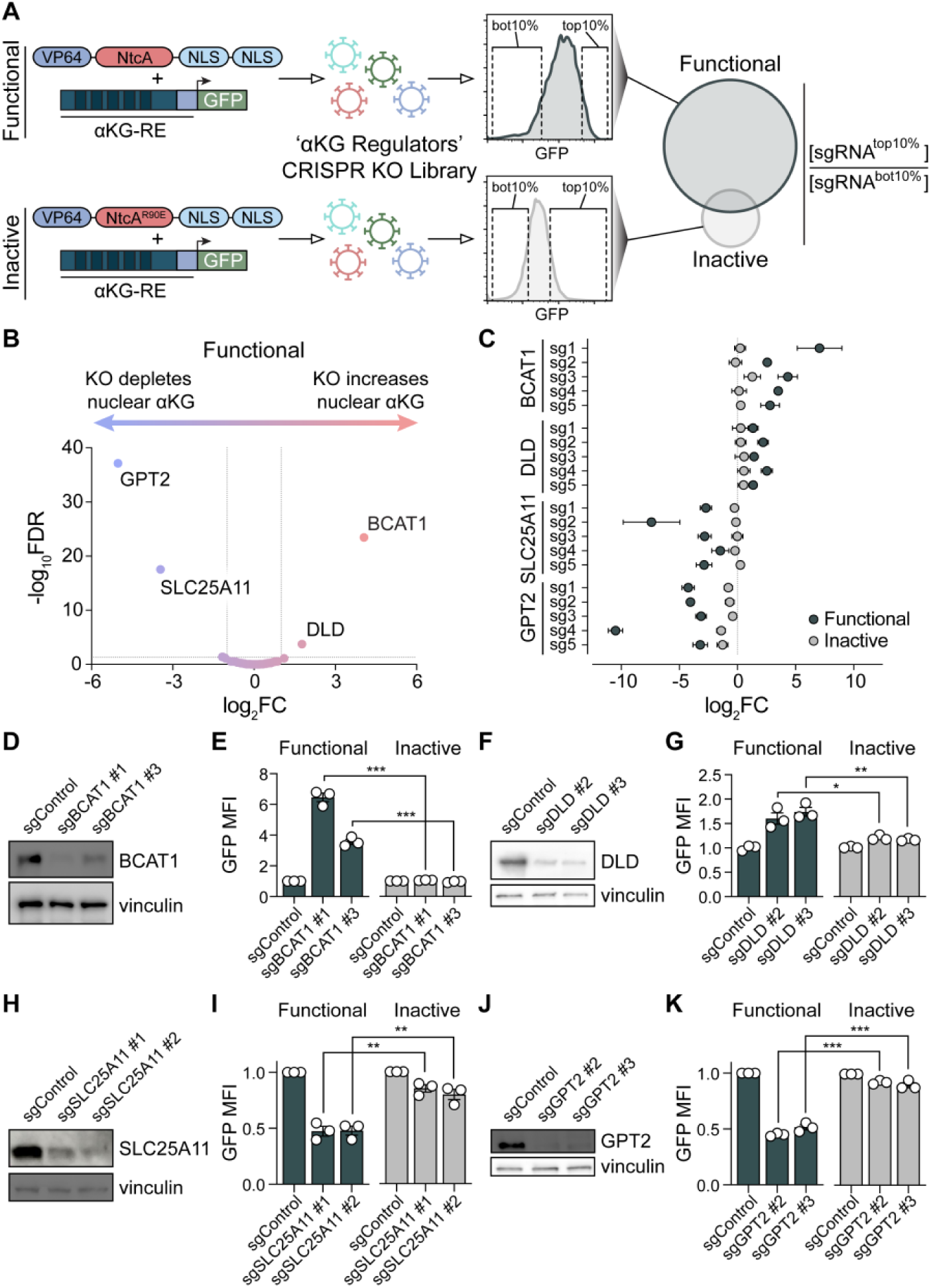
A forward genetic screen reveals molecular regulators of the nuclear αKG pool. (**A**) Schema of a fluorescence activated cell sorting (FACS)-based CRISPR-Cas9 knockout screen to identify genes that control αKG levels in the nucleus. (**B**) Volcano plot of gene-level statistics from the CRISPR-Cas9 screen in U251 cells engineered with the functional αKG-ON biosensor system shown in (A). FC = fold change; FDR = false discovery rate. Gene-level scores were derived from the ratio of sgRNA read counts in the top 10% versus bottom 10% of GFP-expressing cells. (**C**) Enrichment or depletion of individual sgRNAs targeting *BCAT1*, *DLD*, *SLC25A11* or *GPT2* genes from CRISPR-Cas9 screens in U251 cells engineered with functional or inactive versions of the αKG-ON biosensor system. (**D-K**) Validation of nuclear αKG pool regulation by sgRNAs targeting *BCAT1*, *DLD*, *SLC25A11* or *GPT2* genes. (D, F, H, and J) Immunoblots of BCAT1, DLD, SLC25A11 and GPT2 expression in U251 cells engineered to express Cas9 and control or indicated sgRNAs. (E, G, I, and K) Quantification of GFP expression (GFP MFI normalized to sgControl lines) in cells from (D, F, H, and J) engineered with functional or inactive versions of the αKG-ON biosensor system. \**p* < 0.05, \*\**p* < 0.01, \*\*\**p* < 0.001 (unpaired t-tests). In (C, E, G, I, and K), data are means ± SEM.

This screen produced four gene-level “hits”. Knockout of *BCAT1* and *DLD* increased nuclear αKG levels while knockout of *GPT2* and *SLC25A11* depleted this metabolite pool (**Fig. 3B**). Importantly, sgRNAs targeting these four genes were preferentially enriched or depleted in cells expressing the functional versus inactive version of the biosensor (**Fig. 3C** and **fig. S7A**), indicative of αKG-dependent effects. Our findings nominated two mitochondrial proteins, the alanine transaminase GPT2 and the malate/αKG antiporter SLC25A11, as key suppliers of αKG in the nucleus. Conversely, these results implicated the mitochondrial dihydrolipoamide dehydrogenase DLD and the nucleocytosolic BCAA transaminase BCAT1 as suppressors of nuclear αKG. Observing *BCAT1* as a top hit suggested that the screen was successful, given previous work (*39*) and our data linking BCAT1 with negative regulation of the nuclear αKG pool (**Fig. 2L-M**). We validated two sgRNAs targeting each of these genes in low-throughput assays, demonstrating effective depletion of protein products and selective regulation of functional αKG-ON biosensor system output (**Fig. 3D-K**). We also rescued nuclear αKG depletion caused by GPT2 or SLC25A11 knockout by expressing sgRNA-insensitive cDNAs for each target (**fig. S6A-D**), validating on-target sgRNA activity. Genes that did not score as “hits” in our screen included those that encode enzymes that can catabolize or synthesize αKG in mitochondria, including the BCAA transaminase BCAT2, the glutamate dehydrogenase GLUD1, the IDH3A subunit of the isocitrate dehydrogenase 3 (IDH3) complex, and the OGDH component of the oxoglutarate dehydrogenase complex (OGDC) (**fig. S7B**). These results imply that enzymes that can plausibly (based on functional properties) regulate mitochondrion-to- nucleus αKG flux in fact make quite variable contributions to this activity. We validated the inability of BCAT2 to regulate nuclear αKG (**fig. S7C-D**), corroborating our previous experiments (**Fig. 2L**). Likewise, low-throughput experiments confirmed that GLUD1 has a very minor role in controlling nuclear αKG in this model system despite its ability to catalyze αKG synthesis from glutamate (**fig. S7E-F**). We did, however, observe moderate negative and positive regulation of nuclear αKG upon acute IDH3A and OGDH knockout, respectively (**fig. S7G-J**).

These effects were not captured in our screen, indicating that *IDH3A* and *OGDH* genes represent “false negatives”. It may be possible to capture moderate effects (such as those mediated by IDH3A or OGDH knockout) in future screens by reducing the stringency of fluorescence intensity thresholds used for FACS.

To further characterize genes with empirically determined or putative roles in nuclear αKG regulation, we compared whole-cell and nuclear αKG levels in stable lines with knockout of the aforementioned genes (**fig. S8A**). While deletion of most genes produced congruent effects on whole-cell and nuclear αKG pools, knockout of DLD was a notable exception. DLD loss increased nuclear αKG but depleted mitochondrial αKG (**fig. S8B**). These data further highlight ability of the αKG-ON biosensor system to detect compartment-specific changes in αKG abundance that may not be accurately inferred from changes in whole-cell αKG content. We also surveyed how representative sgRNAs targeting each gene of interest impacted cell proliferation. In general, we observed modest or negligible reductions in cell fitness relative to control sgRNA-expressing cells (**fig. S8C-J**). However, one sgRNA targeting OGDH [a common essential gene (*42*)] nearly completely abolished proliferation (**fig. S8K**), an effect that correlated with potent OGDH protein depletion (**fig. S7I**). To ask how sgRNAs targeting essential genes affect biosensor output, we knocked out the ribonucleoprotein gene *SNRPG*. SNRPG-deficient lines failed to grow and displayed increased GFP levels in both functional and inactive biosensor-expressing cells (**fig. S8L-N**). This effect was partly attributable to autofluorescence of dying cells because SNRPG-deficient cells lacking the αKG-RE promoter/GFP reporter construct also displayed higher signal in the GFP channel (**fig. S8O**). These results helped explain enrichment of toxic OGDH sgRNAs in the GFP^high^ population of inactive biosensor-expressing cells in the primary screen (**fig. S7B**) and validated our use of the inactive biosensor to deprioritize sgRNAs that cause αKG-independent changes in GFP signal (**Fig. 3A**). Strategies to exclude autofluorescent cells may improve performance of future FACS-based genetic screens involving the αKG-ON biosensor system. Importantly, sgRNAs targeting the top “hits” in our screen (*BCAT1*, *DLD*, *GPT2*, and *SLC25A11*) did not cause gross defects in cell fitness or mitochondrial function (**fig. S8D-G** and **S8P**), nor did they pervasively increase output of the inactive biosensor (**Fig. 3C**).

### An inter-organelle network controls αKG-dependent chromatin demethylation

We hypothesized that the proteins nominated by our screen form a molecular network that dictates the flow of αKG from mitochondria to nuclei (**Fig. 4A**). We propose that GPT2 contributes a major fraction of the αKG pool that is exported from mitochondria in cells that display robust GPT2-dependent alanine synthesis. Another fraction is controlled by IDH3 and OGDC activities in the TCA cycle, as reflected by changes in biosensor output caused by acute IDH3A, OGDH, and DLD knockout (**Fig. 3F-G** and **fig. S7G-J**). Although OGDH sgRNAs did not score in the screen (likely owing to chronic toxicity), OGDH knockout did increase nuclear αKG in short-term experiments conducted before OGDH depletion caused loss of fitness (**fig. S7I-J**). DLD is the E3 subunit required for generating lipoamide to support catalytic cycles of three mitochondrial enzyme complexes: OGDC, pyruvate dehydrogenase (PDH), and branched chain α-ketoacid dehydrogenase (BCKDH). BCKDH and PDH activities are unlikely to account for DLD-dependent control of the nuclear αKG pool because neither BCAT2 (the upstream regulator of BCKDH) nor PDHA1 (the E1 subunit of PDH) knockout altered functional αKG- ON biosensor system output (**fig. S7C-D** and **S9A-B**). DLD likely regulates nuclear αKG via OGDC. Indeed, DLD loss (**Fig. 3F-G**) phenocopies nuclear αKG accumulation caused by acute OGDH knockout (**fig. S7I-J**). Considering that both DLD and OGDH are required for OGDC activity, it is not clear why DLD sgRNAs preserved cell viability while OGDH sgRNAs did not (**fig. S8E** and **S8K**). To complete this inter-organelle molecular network, our data suggest that mitochondrial αKG produced by GPT2 and TCA cycle activities is exported by the SLC25A11 transporter. In the nucleocytosolic compartment, αKG is catabolized to glutamate by BCAT1 or used as a substrate by Jumonji C domain-containing histone lysine demethylase (KDM) or TET αKG-dependent dioxygenases to promote histone and DNA demethylation, respectively.

**Fig. 4.**
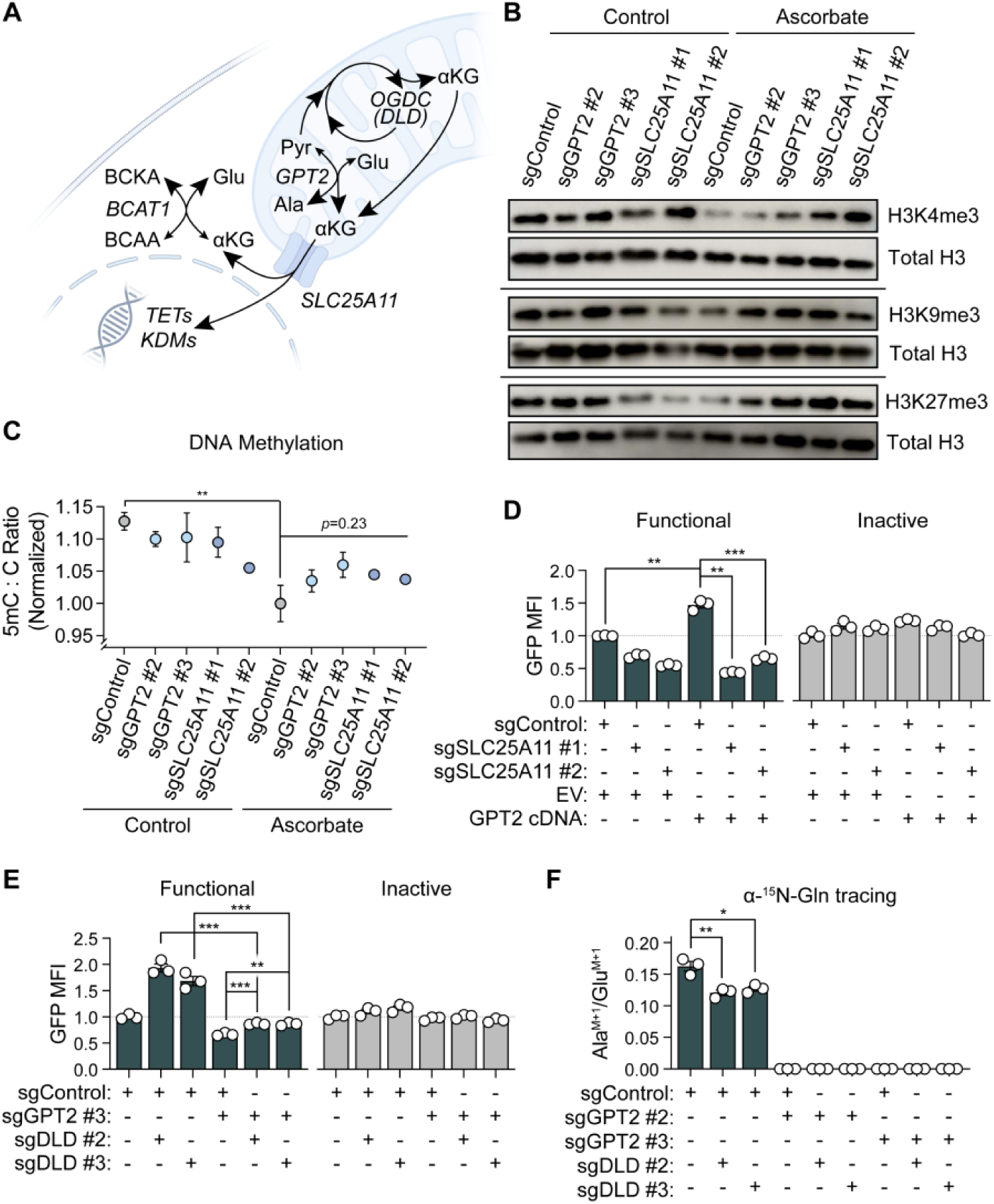
A mitochondrion-to-nucleus pathway of αKG metabolism controls chromatin methylation. (**A**) Schema of localization patterns and functions of proteins that regulate nuclear αKG levels. OGDC = oxoglutarate dehydrogenase complex; Pyr = pyruvate; TETs = TET 5- methylcytosine dioxygenases; KDMs = Jumonji C domain-containing histone lysine demethylases. (**B**) Immunoblots of histone H3 total protein levels or trimethyllysine post- translational modifications in U251 cells engineered to express Cas9 and control, GPT2, or SLC25A11 sgRNAs. Cells were treated with or without ascorbate-2-phosphate prior to harvest. (**C**) Ratios of 5-methyldeoxycytidine (5mC) to deoxycytidine (C) in hydrolyzed genomic DNA isolated from cells in (B). Ratios are normalized to ascorbate-treated sgControl line. \*\**p* < 0.01, (unpaired t-test). *p* = 0.23, (ordinary one-way ANOVA). (**D**) Quantification of GFP expression (GFP MFI normalized to sgControl+EV lines) in U251 cells engineered to express Cas9 and control or SLC25A11 sgRNAs, as well as GPT2 cDNA or EV. Cells were also engineered with functional or inactive versions of the αKG-ON biosensor system. \*\**p* < 0.01, \*\*\**p* < 0.001 (unpaired t-tests). (**E**) Quantification of GFP expression (GFP MFI normalized to sgControl lines) in U251 cells engineered to express Cas9 and control, GPT2, or DLD sgRNAs. Cells were also engineered with functional or inactive versions of the αKG-ON biosensor system. \*\**p* < 0.01, \*\*\**p* < 0.001 (unpaired t-tests). (**F**) α-^15^N-Glutamine stable isotope tracing for 2 hours in U251 cells engineered to express Cas9 and control, GPT2, or DLD sgRNAs. Fractional enrichment in the Ala (M+1) isotopologue relative to fractional enrichment in the Glu (M+1) isotopologue is shown for each line. \**p* < 0.05, \*\**p* < 0.01 (unpaired t-tests). In (C-F), data are means ± SEM.

To test this model, we first asked if nuclear αKG depletion caused by GPT2 or SLC25A11 knockout led to impaired histone or DNA demethylation. We cultured cells with or without GPT2 or SLC25A11 knockout in the presence or absence of ascorbate, a necessary cofactor for KDM and TET dioxygenases. We then measured methylation of H3K4, H3K9, H3K27, and deoxycytidine nucleosides derived from genomic DNA. GPT2 or SLC25A11 knockout caused H3K4 and H3K9 hypermethylation in the presence, but not absence, of ascorbate (**Fig. 4B** and **fig. S9C-D**). A similar trend was observed for H3K27me3 (**fig. S9E**). These data indicate that ascorbate-dependent activation of KDM enzymes is inhibited when the nuclear αKG pool is depleted by either GPT2 or SLC25A11 loss. Methylation of deoxycytidine nucleosides in genomic DNA also increased upon GPT2 or SLC25A11 knockout in an ascorbate-dependent manner (**Fig. 4C**), although genome-wide differences in 5-methyldeoxycitidine (5mC) content did not reach statistical significance. It is possible that significant differences in locus-specific 5mC content are diluted in whole-genome DNA methylation assays; this issue warrants further study. However, the ascorbate-dependent, genome-wide histone hypermethylation phenotypes caused by GPT2 or SLC25A11 loss establish that these mitochondrial proteins are vital sources of αKG for nuclear dioxygenases. This shared function of GPT2 and SLC25A11 is generalizable because knockout of either protein caused nuclear αKG pool depletion in two other neural cell lines: NHA immortalized human astrocytes and TS516 human glioma stem-like cells (**fig. S9F- I**).

### Mitochondrial GPT2 and SLC25A11 activities cooperate to supply the nuclear αKG pool

The functions of GPT2 and SLC25A11 implied that they act in a linear pathway to produce and export αKG, respectively, from mitochondria. Indeed, GPT2 overexpression increased αKG levels in the nucleus in an SLC25A11-dependent manner (**Fig. 4D** and **fig. S11A**), indicating that SLC25A11 is epistatic to GPT2 in the inter-organelle network that regulates nuclear αKG. These data suggest that canonical activity of GPT2 in mitochondria supplies the nuclear αKG pool. To test the subcellular distribution of GPT2 activity, we first quantified GPT2 splice isoforms endogenously expressed by U251 and NHA cells (**fig. S10A-B**). In both cell types, the predominant transcript expressed, ENST00000340124.9, encodes full-length GPT2 protein with a predicted N-terminal MTS (**fig. S10C-D**). However, the second most abundant transcript expressed in both cell lines, ENST00000440783.2, features a distinct 5’ exon structure and encodes a GPT2 protein isoform that lacks an MTS. To ask if the putative extramitochondrial GPT2 protein isoform produced by translation of the ENST00000440783.2 transcript contributes to the nuclear αKG pool, we overexpressed HA-tagged enzymes encoded by each of the two dominant GPT2 transcripts in U251 cells (**fig. S10E**). In contrast to the MTS-encoding full-length transcript ENST00000340124.9, the protein product of ENST00000440783.2 clearly localized to the nucleocytosolic compartment (**fig. S10F**). However, overexpressing this truncated GPT2 enzyme had a negligible effect on nuclear αKG (**fig. S10G**) relative to full-length GPT2 (**Fig. 4D** and **fig. S6D**). We conclude that αKG produced in mitochondria by GPT2 is exported by SLC25A11 to supply the pool of αKG in nuclei.

We next sought to investigate the interaction between GPT2 and DLD. Because TCA cycle and GPT2 activities rely on pyruvate as a substrate, it is possible that DLD knockout elevates αKG levels in the nucleus by diverting pyruvate away from the TCA cycle and toward GPT2.

However, both GPT2-wildtype and GPT2-null cells displayed increased nuclear αKG upon DLD knockout (**Fig. 4E** and **fig. S11B-C**). Knockout of OGDH (which, together with DLD, comprises the OGDC) also increased output of the functional αKG-ON biosensor system independent of GPT2 (**fig. S11D-E**), albeit with confounding effects on output of the inactive biosensor due to loss of cell fitness. To directly ask if DLD loss increases flux through GPT2 in the direction of pyruvate catabolism, we performed stable isotope tracing with α-^15^N-glutamine in control cells or those lacking GPT2, DLD, or both enzymes (**Fig. 4F** and **fig. S11F**). Together with pyruvate, glutamate (a portion of which is produced by glutamine catabolism) serves as a substrate for GPT2. Therefore, we evaluated the ratio of alanine labeling to glutamate labeling as a marker of GPT2 activity. We found that DLD knockout reduced, rather than enhanced, GPT2 flux in the direction of pyruvate catabolism. GPT2 knockout reduced the ratio of labeled alanine to labeled glutamate to zero, thereby validating this marker of GPT2 activity. Together, these studies suggest that GPT2 and DLD occupy critical but distinct, disconnected nodes within the nuclear αKG regulatory network.

### A mouse model of GPT2 deficiency displays chromatin hypermethylation in the developing brain

To address the potential physiological relevance of our findings, we studied a mouse model of GPT2 deficiency. GPT2 deficiency is a relatively recently described inborn error of metabolism caused by autosomal recessive inheritance of loss-of-function mutations in the *GPT2* gene (*43–46*). These mutations cause neurological and intellectual impairment. In mice, whole-body Gpt2 knockout leads to death 3-5 weeks after birth (*47*). This phenotype is fully recapitulated by neuron-specific knockout of Gpt2, underscoring a critical function for this enzyme in supporting neuronal development and homeostasis (*48*). Consistent with this key role for GPT2 activity in the central nervous system (CNS), we surveyed global histone methylation profiles of various tissues from whole-body Gpt2 knockout (KO) and Gpt2 wildtype (WT) mice and observed pervasive histone hypermethylation in the brains of Gpt2 KO animals at postnatal day 19 (P19) to P21 (**Fig. 5A** and **fig. S12A-C**). H3K4 and H3K9 trimethylation were robustly increased in brain tissues of Gpt2 KO versus WT mice, while we observed a moderate elevation of H3K27 trimethylation. In contrast, histone methylation patterns in the kidney, an organ with high Gpt2 expression, were comparable between Gpt2 WT and KO tissues (**Fig. 5B**). There was generally little effect of GPT2 loss on histone methylation in other organs (including heart, lung, spleen, liver, and colon tissues) (**fig. S12D-H**). We did, however, observe hypermethylated H3K9 as well as H3K9 and H3K27 residues in heart and spleen tissues, respectively, from Gpt2 KO mice. We also evaluated genome-wide DNA methylation in brain and kidney tissues from these mice. Consistent with effects of GPT2 knockout in cultured cells (**Fig. 4C**), we observed a trend toward global 5mC enrichment in genomic DNA extracts from Gpt2 KO versus Gpt2 WT mouse brain tissues (**Fig. 5C**). This trend was specific to brain, as Gpt2 loss did not alter genomic 5mC levels in kidney tissues (**Fig. 5D**). Therefore, Gpt2 loss causes disproportionate chromatin hypermethylation in the developing brain.

**Fig. 5.**
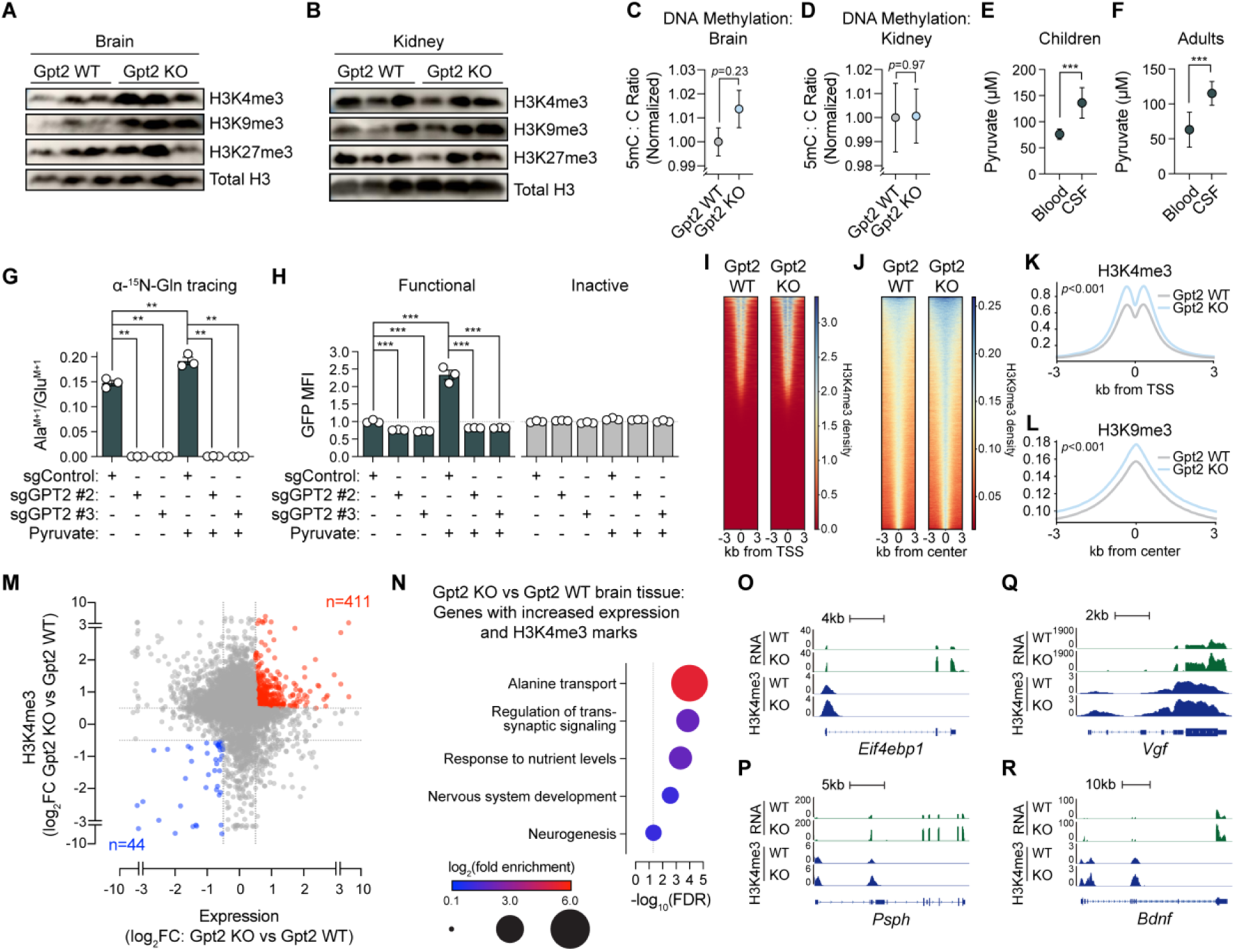
Gpt2 deficiency causes chromatin hypermethylation and transcriptional dysregulation in the brain. (**A and B**) Immunoblots of histone H3 total protein levels or trimethyllysine post-translational modifications in brain (A) or kidney (B) tissues from Gpt2 WT or Gpt2 KO mice at postnatal day 19 (P19) to P21. (**C and D**) Ratios of 5-methyldeoxycytidine (5mC) to deoxycytidine (C) in hydrolyzed genomic DNA isolated from brain (C) or kidney (D) tissues from Gpt2 WT or Gpt2 KO mice. *p* values were calculated by unpaired t-tests. (**E and F**) Concentrations of pyruvate in blood or CSF samples from children (E) or adults (F). Data are derived from reference (*49*). \*\*\**p* < 0.001 (unpaired t-tests). (**G**) α-^15^N-Glutamine stable isotope tracing for 2 hours in U251 cells engineered to express Cas9 and control or GPT2 sgRNAs and cultured with or without pyruvate. Fractional enrichment in the Ala (M+1) isotopologue relative to fractional enrichment in the Glu (M+1) isotopologue is shown for each condition. \*\**p* < 0.01 (unpaired t-tests). (**H**) Quantification of GFP expression (GFP MFI normalized to sgControl, pyruvate-free conditions) in U251 cells engineered and cultured as in (G). Cells were also engineered with functional or inactive versions of the αKG-ON biosensor system. \*\*\**p* < 0.001 (unpaired t-tests). (**I-L**) H3K4me3 or H3K9me3 ChIP-seq results from chromatin extracted from brain tissues of Gpt2 WT or Gpt2 KO mice. Tornado plots (I and J) and quantification of ChIP- seq signal (K and L) of aligned histone methylation densities are shown. *p* values were calculated by Kolmogorov-Smirnov tests. (**M**) Integrative analysis of H3K4me3 ChIP-seq and RNA-seq datasets derived from chromatin and RNA, respectively, isolated from brain tissues of Gpt2 WT and Gpt2 KO mice. Red indicates H3K4me3 hypermethylated and upregulated genes in Gpt2 KO condition; blue indicates H3K4me3 hypomethylated and downregulated genes in Gpt2 KO condition. (**N**) Gene ontology analysis of H3K4me3 hypermethylated and upregulated genes in Gpt2 KO mouse brain tissues identified in (M). (**O-R**) Density tracks for H3K4me3 ChIP-seq and RNA-seq reads associated with representative H3K4me3 hypermethylated and upregulated genes in Gpt2 KO mouse brain tissues. In (C-H), data are means ± SEM.

### Pyruvate is enriched in cerebrospinal fluid and dictates GPT2-dependent αKG synthesis

GPT2 is expressed in many tissues, indicating that differences in enzyme abundance alone cannot explain preferential effects of GPT2 loss on the CNS in mice and humans. We hypothesized that tissue-specific differences in the levels of GPT2 substrates determine the directionality and magnitude of GPT2 flux via mass action. Although we have focused on GPT2- dependent αKG and alanine synthesis (**Fig. 4F**), GPT2 is reversible and can also catabolize αKG and alanine, yielding pyruvate and glutamate. Alanine levels are reduced in brain tissues of Gpt2 KO mice (*44*), suggesting that net flux through GPT2 likely moves in the direction of αKG and alanine synthesis in brain tissue in vivo. Therefore, we asked if increased pyruvate availability may be a driver of GPT2-dependent αKG and alanine synthesis in the CNS. Intriguingly, the pyruvate content of cerebrospinal fluid (CSF) is nearly 2-fold higher than that of blood in both children and adults (**Fig. 5E-F**) (*49*), suggesting that the local supply of pyruvate may be higher in the CNS than in peripheral organs. To determine whether pyruvate availability affects GPT2- dependent supply of the nuclear αKG pool, we evaluated α-^15^N-glutamine stable isotope tracing and αKG-ON biosensor system output in control or GPT2 KO cells cultured in medium with or without pyruvate. Pyruvate supplementation stimulated GPT2 flux in the direction of alanine and αKG synthesis (**Fig. 5G** and **fig. S12I**) and markedly elevated nuclear αKG (**Fig. 5H**). Critically, GPT2 knockout abolished both effects. These results suggest that the preferential epigenetic regulatory function of GPT2 in the CNS is driven in part by high microenvironmental pyruvate concentrations.

### Epigenetic reprogramming caused by Gpt2 loss affects genes involved in neurogenesis

Although most research on the molecular pathogenesis of GPT2 deficiency has focused on defects in alanine biosynthesis, glutamine-dependent TCA cycle anaplerosis, and glutamatergic neurotransmission (*44*, *50*), our work suggests that depletion of nuclear αKG and chromatin hypermethylation may contribute to disease phenotypes as well. To better characterize epigenetic reprogramming caused by Gpt2 loss in neural cells, we performed chromatin immunoprecipitation followed by sequencing (ChIP-seq) for H3K4me3 and H3K9me3 marks in chromatin preparations from Gpt2 KO and Gpt2 WT mouse brains. In agreement with our immunoblot analyses of global H3K4me3 and H3K9me3 in these tissues (**Fig. 5A**), we found enrichment of these marks at loci throughout the genome in brain tissues from Gpt2 KO mice at P19-21(**Fig. 5I-L**). H3K4me3 marks that promote gene transactivation were enriched, as expected, around transcriptional start sites (TSSs), suggesting that this epigenetic phenotype may elicit an aberrant gene expression program. To identify genes that are both H3K4 hypermethylated and overexpressed in the brains of Gpt2 KO mice, we performed RNA-sequencing (RNA-seq) from brains of Gpt2 KO and WT mice and integrated the resulting dataset with the H3K4me3 ChIP-seq dataset (**Fig. 5M**). Over 400 genes upregulated in Gpt2 KO mouse brain tissues also displayed H3K4 hypermethylation, thereby identifying a subset of transcriptional changes caused by Gpt2 loss that are linked to nuclear αKG pool depletion and subsequent chromatin hypermethylation.

We performed a gene ontology (GO) analysis to ask if this collection of overexpressed and H3K4 hypermethylated genes may be involved in disease processes associated with GPT2 deficiency. Two of the most significantly enriched GO terms related to alanine transport and nutrient deprivation (**Fig. 5N**), suggesting that H3K4me3 hypermethylation may augment the transcriptional response to loss of alanine synthesis. In addition, genes related to electrochemical signaling and nervous system development were also significantly altered. We constructed tracks from RNA-seq and H3K4me3 ChIP-seq reads for four representative genes (two associated with the response to nutrient deprivation and two with neurogenesis) to highlight the close association between epigenetic and transcriptomic reprogramming in Gpt2 KO mice (**Fig. 5O-R** and **fig. S12J-M**). These genes include *Eif4ebp1* and *Psph*, which are stimulated by amino acid deprivation, and *Vgf* and *Bdnf*, which encode neuropeptides involved in neural cell signaling, synaptic plasticity, and neurogenesis. Each of these genes shows coincident increases in expression and H3K4 methylation upon Gpt2 loss in the brain. Taken together, these data establish a key epigenetic regulatory function for GPT2 in vivo and implicate aberrant αKG signaling to chromatin as a driver of the molecular pathogenesis of GPT2 deficiency.

## Discussion

We describe a new research tool, the αKG-ON biosensor system, that enables robust, specific measurement of changes in nuclear αKG pool size in human cells. Our work expands and diversifies the techniques available to measure αKG in live cells. While previous work in this space has focused on fluorescence resonance energy transfer (FRET)-based tools to monitor αKG predominantly in bacteria (*23*, *51–56*), the system we report may be advantageous for applications in which compartmental specificity, high sensitivity, and enhanced stability of αKG sensing in human cells is desired. We leveraged these unique attributes of the αKG-ON biosensor system to conduct a forward genetic screen using nuclear αKG abundance as the experimental endpoint. This approach enabled systematic evaluation of the contributions of individual genes to nuclear αKG control. Our results inform a molecular network spanning mitochondrial, cytoplasmic, and nuclear compartments that controls αKG levels in the nucleus. We found that GPT2 and TCA cycle activities regulate the abundance of αKG that is produced in the mitochondria and subsequently exported to the cytosol via SLC25A11. This αKG pool sustains dioxygenases involved in chromatin demethylation in the nucleus and is suppressed by BCAT1, in agreement with prior research (*39*). Importantly, we uncovered a vital role for GPT2 activity in establishing appropriate chromatin methylation patterns in the developing brain, thereby corroborating physiological relevance of the nuclear αKG regulatory network we describe.

Our findings suggest that nuclear αKG is controlled differently than the nuclear pools of several other metabolites that serve as substrates for chromatin-modifying enzymes. Indeed, nuclear synthesis of acetyl-CoA (*57*), propionyl-CoA (*58*), and S-adenosylmethionine (SAM) (*59*) has been proposed to provide local sources of these metabolites to promote chromatin modification. Accordingly, ATP-citrate lyase (ACLY) (*60*, *61*), acetyl-CoA synthetase 2 (ACSS2) (*62–64*), pyruvate dehydrogenase (PDH) (*57*, *65*), and methionine adenosyltransferase (MAT) (*66–68*) enzymes have been shown to localize to the nucleus. This contrasts with our results showing that nuclear αKG is principally derived from mitochondria. Although GPT2 has been reported to localize to the nucleus (*69*), we find that an endogenous, extramitochondrial GPT2 isoform lacking an MTS does not affect αKG levels in the nucleus. Further affirming the relevance of mitochondrial GPT2 activity, our data show that the mitochondrial malate/αKG antiporter SLC25A11 is epistatic to GPT2 in the nuclear αKG regulatory network. This key role of SLC25A11 in promoting mitochondrion-to-nucleus αKG flux aligns with a study of familial paraganglioma that showed that *SLC25A11* loss-of-function mutations phenocopy chromatin hypermethylation caused by oncometabolite-producing *SDH* or *FH* gene mutations (*70*).

Our conceptual model of nuclear αKG control provides insight into the pathogenesis of GPT2 deficiency. Patients with biallelic inactivating mutations in *GPT2* suffer from neurodevelopmental disorder with spastic paraplegia and microcephaly (NEDSPM). Symptoms of this disorder include global developmental delay, hypotonia, delays or failure to develop speech and walking, and epilepsy (*43–46*). Defects in neuronal function are central to this disorder, as we (R.J.D. and E.M.M.) previously showed that neuron-specific GPT2 loss recapitulates lethality of whole-body Gpt2 knockout in mice (*47*). Importantly, alanine supplementation partially rescues this phenotype, indicating that inhibition of GPT2 catalysis in the direction of alanine and αKG synthesis in neurons impairs CNS development (*47*). In addition to sustaining alanine pools, our data highlight a distinct yet important function of GPT2: supplying αKG to nuclear dioxygenases to facilitate chromatin demethylation. This function likely affects neuronal maturation because genes related to neurogenesis were dysregulated by GPT2 loss at both epigenetic and transcriptomic levels. Among these genes, we found *Vgf* to be strongly H3K4 hypermethylated and overexpressed in brain tissues of Gpt2 knockout mice versus controls. *Vgf* encodes a neuropeptide that, when overexpressed in transgenic mice, is sufficient to cause morphological defects in the striatum and behavioral abnormalities (*71*). Therefore, our study nominates nuclear αKG pool depletion and epigenetic reprogramming as contributors to the pathology of GPT2 deficiency.

Our work provides tools and a technical framework that may facilitate future efforts to study compartmentalized metabolism of αKG and other small molecules. The αKG-ON biosensor system may open new avenues to evaluate αKG signaling to chromatin in physiological and pathophysiological contexts beyond GPT2 deficiency, including, for example, developmental cell state transitions (*5*) and cancers with distinctive chromatin methylation states that are not linked to oncometabolites (*72–75*). Moreover, directed evolution of the NtcA transcription factor or exploitation of other evolutionarily conserved metabolite-responsive transcription factors may foster design of transcriptional biosensors for other nuclear metabolites. Irrespective of the metabolite of interest, we unveil a strategy to leverage unbiased functional genomics approaches to identify molecular regulators of metabolite pools. Our work provides proof of principle that dedicated biosensors can render endogenous metabolite pools suitable endpoints for FACS-based forward genetic screens in human cells. This experimental strategy may promote use of genetic screens to study metabolic processes that do not directly regulate cell fitness, the most common endpoint presently used in such screens.

## Acknowledgments

The authors thank members of the McBrayer, DeBerardinis, and Abdullah laboratories for insightful feedback; Dr. Ingo Mellinghoff (Memorial Sloan Kettering Cancer Center) for TS516 GBM cells; Dr. William G. Kaelin, Jr. (Dana-Farber) for DNA plasmids; Dr. Russell Pieper (UCSF) for NHA cells; Dr. Ryan Looper (Univ. of Utah) for (*R*)-2HG-TFMB; members of the Genetic Perturbation Platform (GPP) at the Broad Institute of MIT and Harvard for CRISPR screening resources; and Prof. Javier Florencio and Dr. M. Isabel Muro Pastor (Instituto de Bioquímica Vegetal y Fotosíntesis) for NtcA anti-serum. This work was supported by grants from the National Institutes of Health/National Cancer Institute (NIH/NCI) (R01CA289260, R01CA258586, P50CA165962, U19CA264504), the Cancer Prevention and Research Institute of Texas (CPRIT) (RR190034, RP230344, RP2400489, RP250278), a Distinguished Scientist Award from the Sontag Foundation, and gifts from the Jonesville Foundation and the Nick Gonzales Foundation for Brain Tumor Research to S.K.M. D.D.S. was supported by an NIH/NCI grant (K12CA0903354), a Burroughs Wellcome Career Award for Medical Scientists, and a Lubin Family Foundation Scholar Award. M.R.S. was supported by an NIH/NCI grant (F30CA271634). Y.X. was supported by an NIH/NCI grant (K99CA277576) and a Human Frontier Science Program postdoctoral fellowship award (LT0018/2022-L). L.G.Z., T.P.M. and the CRI Metabolomics Facility are supported by a CPRIT Core Facilities Support Award (RP240494). L.A.B. was supported by grants from the NIH/National Institute of General Medical Sciences (NIGMS) (R35GM124958), NIH/National Institute of Child Health and Human Development (NICHD) (R01HD109239), the Welch Foundation (I-2025), and American Cancer Society (ACS) (134230-RSG-20-043-01-DMC). M.A. was supported by a grant from NIH/National Institute of Diabetes and Digestive and Kidney Diseases (NIDDK) (R01DK125713). R.J.D. is an HHMI Investigator. E.M.M. was supported by grants from the NIH/National Institute on Aging (R01AG0874550) and NIH/National Institute of Neurological Disorders and Stroke (R01NS121618 and R01NS113141). Diagrams were produced in Adobe Illustrator using material from Biorender (https://biorender.com).

## Author contributions

Conceptualization: ACS, VK, KGA, RJD, EMM, SKM

Data curation: HL

Formal analysis: ACS, HL, JIT, LG, XZ

Funding acquisition: EMM, SKM

Investigation: ACS, HL, JIT, JHJ, DDS, YX, JHL, LGZ, TPM, RG, YJK, KG, QO, MS, MF

Methodology: ACS, JHJ, KG, QO, MS, MF, MA

Project administration: KGA, EMM, SKM

Software: XZ, MRS, LAB

Resources: LG, XZ, KG, QO, MS, MF, MRS, JGD, LAB

Supervision: DDS, TPM, LX, KGA, LAB, MA, EMM, SKM

Validation: HL

Visualization: ACS, HL, SKM

Writing – original draft: ACS, HL, SKM

Writing – review & editing: ACS, HL, VK, LAB, MA, RJD, EMM, SKM

## Data and materials availability

Further information and requests for resources and reagents should be directed to and will be fulfilled by the lead contact, Samuel K. McBrayer. DNA constructs generated in this study are available upon request or from Addgene. All relevant data are available upon request. All other data are available in the main text or the supplementary materials. Omics data will be deposited to the Gene Expression Omnibus (GEO) website. Synthetic DNA sequences will be deposited to GenBank.

## Supplementary Materials

Materials and Methods Figs. S1 to S12

Tables S1 to S5 References (*1–12*)

## References and Notes

1. S. L. Campbell, K. E. Wellen, Metabolic Signaling to the Nucleus in Cancer. Mol Cell 71, 398–408 (2018).

2. J. L. Meier, Metabolic mechanisms of epigenetic regulation. ACS Chem Biol 8, 2607–2621 (2013).

3. J. M. Schvartzman, C. B. Thompson, L. W. S. Finley, Metabolic regulation of chromatin modifications and gene expression. J Cell Biol 217, 2247–2259 (2018).

4. W. G. Kaelin, S. L. McKnight, Influence of metabolism on epigenetics and disease. Cell 153, 56–69 (2013).

5. B. W. Carey, L. W. S. Finley, J. R. Cross, C. D. Allis, C. B. Thompson, Intracellular α- ketoglutarate maintains the pluripotency of embryonic stem cells. Nature 518, 413–416 (2015).

6. P. A. Tyrakis, A. Palazon, D. Macias, K. L. Lee, A. T. Phan, P. Veliça, J. You, G. S. Chia, J. Sim, A. Doedens, A. Abelanet, C. E. Evans, J. R. Griffiths, L. Poellinger, A. W. Goldrath, R. S. Johnson, S-2-hydroxyglutarate regulates CD8+ T-lymphocyte fate. Nature 540, 236–241 (2016).

7. T. Q. Tran, E. A. Hanse, A. N. Habowski, H. Li, M. B. Ishak Gabra, Y. Yang, X. H. Lowman, A. M. Ooi, S. Y. Liao, R. A. Edwards, M. L. Waterman, M. Kong, α- Ketoglutarate attenuates Wnt signaling and drives differentiation in colorectal cancer. Nat Cancer 1, 345–358 (2020).

8. J. P. Morris, J. J. Yashinskie, R. Koche, R. Chandwani, S. Tian, C.-C. Chen, T. Baslan, Z. S. Marinkovic, F. J. Sánchez-Rivera, S. D. Leach, C. Carmona-Fontaine, C. B. Thompson, L. W. S. Finley, S. W. Lowe, α-Ketoglutarate links p53 to cell fate during tumour suppression. Nature 573, 595–599 (2019).

9. P.-S. Liu, H. Wang, X. Li, T. Chao, T. Teav, S. Christen, G. Di Conza, W.-C. Cheng, C.-H. Chou, M. Vavakova, C. Muret, K. Debackere, M. Mazzone, H.-D. Huang, S.-M. Fendt, J. Ivanisevic, P.-C. Ho, α-ketoglutarate orchestrates macrophage activation through metabolic and epigenetic reprogramming. Nat Immunol 18, 985–994 (2017).

10. J. R. Toro, M. L. Nickerson, M. H. Wei, M. B. Warren, G. M. Glenn, M. L. Turner, L. Stewart, P. Duray, O. Tourre, N. Sharma, P. Choyke, P. Stratton, M. Merino, M. M. Walther, W. M. Linehan, L. S. Schmidt, B. Zbar, Mutations in the fumarate hydratase gene cause hereditary leiomyomatosis and renal cell cancer in families in North America. Am J Hum Genet 73, 95–106 (2003).

11. M. Xiao, H. Yang, W. Xu, S. Ma, H. Lin, H. Zhu, L. Liu, Y. Liu, C. Yang, Y. Xu, S. Zhao, D. Ye, Y. Xiong, K. L. Guan, Inhibition of alpha-KG-dependent histone and DNA demethylases by fumarate and succinate that are accumulated in mutations of FH and SDH tumor suppressors. Genes Dev 26, 1326–38 (2012).

12. W. Xu, H. Yang, Y. Liu, Y. Yang, P. Wang, S. H. Kim, S. Ito, C. Yang, M. T. Xiao, L. X. Liu, W. Q. Jiang, J. Liu, J. Y. Zhang, B. Wang, S. Frye, Y. Zhang, Y. H. Xu, Q. Y. Lei, K. L. Guan, S. M. Zhao, Y. Xiong, Oncometabolite 2-hydroxyglutarate is a competitive inhibitor of alpha-ketoglutarate-dependent dioxygenases. Cancer Cell 19, 17–30 (2011).

13. S. Turcan, D. Rohle, A. Goenka, L. A. Walsh, F. Fang, E. Yilmaz, C. Campos, A. W. Fabius, C. Lu, P. S. Ward, C. B. Thompson, A. Kaufman, O. Guryanova, R. Levine, A. Heguy, A. Viale, L. G. Morris, J. T. Huse, I. K. Mellinghoff, T. A. Chan, IDH1 mutation is sufficient to establish the glioma hypermethylator phenotype. Nature 483, 479–83 (2012).

14. G. J. Rahme, N. M. Javed, K. L. Puorro, S. Xin, V. Hovestadt, S. E. Johnstone, B. E. Bernstein, Modeling epigenetic lesions that cause gliomas. Cell 186, 3674–3685.e14 (2023).

15. W. A. Flavahan, Y. Drier, B. B. Liau, S. M. Gillespie, A. S. Venteicher, A. O. Stemmer-Rachamimov, M. L. Suva, B. E. Bernstein, Insulator dysfunction and oncogene activation in IDH mutant gliomas. Nature 529, 110–4 (2016).

16. W. A. Flavahan, Y. Drier, S. E. Johnstone, M. L. Hemming, D. R. Tarjan, E. Hegazi, S. J. Shareef, N. M. Javed, C. P. Raut, B. K. Eschle, P. C. Gokhale, J. L. Hornick, E. T. Sicinska, G. D. Demetri, B. E. Bernstein, Altered chromosomal topology drives oncogenic programs in SDH-deficient GISTs. Nature 575, 229–233 (2019).

17. 17. Cancer Genome Atlas Research Network, W. M. Linehan, P. T. Spellman, C. J. Ricketts, C. J. Creighton, S. S. Fei, C. Davis, D. A. Wheeler, B. A. Murray, L. Schmidt, C. D. Vocke, M. Peto, A. A. M. Al Mamun, E. Shinbrot, A. Sethi, S. Brooks, W. K. Rathmell, A. N. Brooks, K. A. Hoadley, A. G. Robertson, D. Brooks, R. Bowlby, S. Sadeghi, H. Shen, D. J. Weisenberger, M. Bootwalla, S. B. Baylin, P. W. Laird, A. D. Cherniack, G. Saksena, S. Haake, J. Li, H. Liang, Y. Lu, G. B. Mills, R. Akbani, M. D. M. Leiserson, B. J. Raphael, P. Anur, D. Bottaro, L. Albiges, N. Barnabas, T. K. Choueiri, B. Czerniak, A. K. Godwin, A. A. Hakimi, T. H. Ho, J. Hsieh, M. Ittmann, W. Y. Kim, B. Krishnan, M. J. Merino, K. R. Mills Shaw, V. E. Reuter, E. Reznik, C. S. Shelley, B. Shuch, S. Signoretti, R. Srinivasan, P. Tamboli, G. Thomas, S. Tickoo, K. Burnett, D. Crain, J. Gardner, K. Lau, D. Mallery, S. Morris, J. D. Paulauskis, R. J. Penny, C. Shelton, W. T. Shelton, M. Sherman, E. Thompson, P. Yena, M. T. Avedon, J. Bowen, J. M. Gastier-Foster, M. Gerken, K. M. Leraas, T. M. Lichtenberg, N. C. Ramirez, T. Santos, L. Wise, E. Zmuda, J. A. Demchok, I. Felau, C. M. Hutter, M. Sheth, H. J. Sofia, R. Tarnuzzer, Z. Wang, L. Yang, J. C. Zenklusen, J. Zhang, B. Ayala, J. Baboud, S. Chudamani, J. Liu, L. Lolla, R. Naresh, T. Pihl, Q. Sun, Y. Wan, Y. Wu, A. Ally, M. Balasundaram, S. Balu, R. Beroukhim, T. Bodenheimer, C. Buhay, Y. S. N. Butterfield, R. Carlsen, S. L. Carter, H. Chao, E. Chuah, A. Clarke, K. R. Covington, M. Dahdouli, N. Dewal, N. Dhalla, H. V. Doddapaneni, J. A. Drummond, S. B. Gabriel, R. A. Gibbs, R. Guin, W. Hale, A. Hawes, D. N. Hayes, R. A. Holt, A. P. Hoyle, S. R. Jefferys, S. J. M. Jones, C. D. Jones, D. Kalra, C. Kovar, L. Lewis, J. Li, Y. Ma, M. A. Marra, M. Mayo, S. Meng, M. Meyerson, P. A. Mieczkowski, R. A. Moore, D. Morton, L. E. Mose, A. J. Mungall, D. Muzny, J. S. Parker, C. M. Perou, J. Roach, J. E. Schein, S. E. Schumacher, Y. Shi, J. V. Simons, P. Sipahimalani, T. Skelly, M. G. Soloway, C. Sougnez, A. Tam, D. Tan, N. Thiessen, U. Veluvolu, M. Wang, M. D. Wilkerson, T. Wong, J. Wu, L. Xi, J. Zhou, J. Bedford, F. Chen, Y. Fu, M. Gerstein, D. Haussler, K. Kasaian, P. Lai, S. Ling, A. Radenbaugh, D. Van Den Berg, J. N. Weinstein, J. Zhu, M. Albert, I. Alexopoulou, J. J. Andersen, J. T. Auman, J. Bartlett, S. Bastacky, J. Bergsten, M. L. Blute, L. Boice, R. J. Bollag, J. Boyd, E. Castle, Y.-B. Chen, J. C. Cheville, E. Curley, B. Davies, A. DeVolk, R. Dhir, L. Dike, J. Eckman, J. Engel, J. Harr, R. Hrebinko, M. Huang, L. Huelsenbeck-Dill, M. Iacocca, B. Jacobs, M. Lobis, J. K. Maranchie, S. McMeekin, J. Myers, J. Nelson, J. Parfitt, A. Parwani, N. Petrelli, B. Rabeno, S. Roy, A. L. Salner, J. Slaton, M. Stanton, R. H. Thompson, L. Thorne, K. Tucker, P. M. Weinberger, C. Winemiller, L. A. Zach, R. Zuna, Comprehensive Molecular Characterization of Papillary Renal-Cell Carcinoma. N Engl J Med 374, 135–145 (2016).

18. E. Letouzé, C. Martinelli, C. Loriot, N. Burnichon, N. Abermil, C. Ottolenghi, M. Janin, M. Menara, A. T. Nguyen, P. Benit, A. Buffet, C. Marcaillou, J. Bertherat, L. Amar, P. Rustin, A. De Reyniès, A.-P. Gimenez-Roqueplo, J. Favier, SDH mutations establish a hypermethylator phenotype in paraganglioma. Cancer Cell 23, 739–752 (2013).

19. M. I. Muro-Pastor, J. C. Reyes, F. J. Florencio, Ammonium assimilation in cyanobacteria. Photosynth Res 83, 135–150 (2005).

20. M.-X. Zhao, Y.-L. Jiang, Y.-X. He, Y.-F. Chen, Y.-B. Teng, Y. Chen, C.-C. Zhang, C.-Z. Zhou, Structural basis for the allosteric control of the global transcription factor NtcA by the nitrogen starvation signal 2-oxoglutarate. Proc Natl Acad Sci U S A 107, 12487–12492 (2010).

21. M. F. Vazquez-Bermudez, A. Herrero, E. Flores, 2-Oxoglutarate increases the binding affinity of the NtcA (nitrogen control) transcription factor for the Synechococcus glnA promoter. FEBS Lett 512, 71–4 (2002).

22. J. C. Reyes, M. I. Muro-Pastor, F. J. Florencio, Transcription of glutamine synthetase genes (glnA and glnN) from the cyanobacterium Synechocystis sp. strain PCC 6803 is differently regulated in response to nitrogen availability. J Bacteriol 179, 2678–2689 (1997).

23. H.-L. Chen, A. Latifi, C.-C. Zhang, C. S. Bernard, Biosensors-Based In Vivo Quantification of 2-Oxoglutarate in Cyanobacteria and Proteobacteria. Life (Basel) 8, 51 (2018).

24. M. Gossen, S. Freundlieb, G. Bender, G. Muller, W. Hillen, H. Bujard, Transcriptional activation by tetracyclines in mammalian cells. Science 268, 1766–9 (1995).

25. M. Gossen, H. Bujard, Tight control of gene expression in mammalian cells by tetracycline-responsive promoters. Proc. Natl. Acad. Sci. (USA) 89, 5547–5551 (1992).

26. K. Forchhammer, K. A. Selim, Carbon/nitrogen homeostasis control in cyanobacteria. FEMS Microbiol Rev 44, 33–53 (2020).

27. S.-J. Han, Y.-L. Jiang, L.-L. You, L.-Q. Shen, X. Wu, F. Yang, N. Cui, W.-W. Kong, H. Sun, K. Zhou, H.-C. Meng, Z.-P. Chen, Y. Chen, Y. Zhang, C.-Z. Zhou, DNA looping mediates cooperative transcription activation. Nat Struct Mol Biol 31, 293–299 (2024).

28. J. L. Crespo, M. García-Domínguez, F. J. Florencio, Nitrogen control of the glnN gene that codes for GS type III, the only glutamine synthetase in the cyanobacterium Pseudanabaena sp. PCC 6903. Mol Microbiol 30, 1101–1112 (1998).

29. C.-A. Dudek, D. Jahn, PRODORIC: state-of-the-art database of prokaryotic gene regulation. Nucleic Acids Res 50, D295–D302 (2022).

30. P. M. Lieberman, A. J. Berk, The Zta trans-activator protein stabilizes TFIID association with promoter DNA by direct protein-protein interaction. Genes Dev 5, 2441–2454 (1991).

31. A. Chavez, J. Scheiman, S. Vora, B. W. Pruitt, M. Tuttle, E. P R Iyer, S. Lin, S. Kiani, C. D. Guzman, D. J. Wiegand, D. Ter-Ovanesyan, J. L. Braff, N. Davidsohn, B. E. Housden, N. Perrimon, R. Weiss, J. Aach, J. J. Collins, G. M. Church, Highly efficient Cas9-mediated transcriptional programming. Nat Methods 12, 326–328 (2015).

32. E. K. Flemington, A. M. Borras, J. P. Lytle, S. H. Speck, Characterization of the Epstein-Barr Virus BZLF1 Protein Transactivation Domain. J. Virol. 66, 922–929 (1992).

33. K. Akagi, M. Kanai, H. Saya, T. Kozu, A. Berns, A novel tetracycline-dependent transactivator with E2F4 transcriptional activation domain. Nucleic Acids Res 29, E23 (2001).

34. S. Rogers, R. Wells, M. Rechsteiner, Amino acid sequences common to rapidly degraded proteins: the PEST hypothesis. Science 234, 364–368 (1986).

35. X. Li, X. Zhao, Y. Fang, X. Jiang, T. Duong, C. Fan, C. C. Huang, S. R. Kain, Generation of destabilized green fluorescent protein as a transcription reporter. J Biol Chem 273, 34970–34975 (1998).

36. J. A. Losman, R. E. Looper, P. Koivunen, S. Lee, R. K. Schneider, C. McMahon, G. S. Cowley, D. E. Root, B. L. Ebert, W. G. Kaelin, (R)-2-hydroxyglutarate is sufficient to promote leukemogenesis and its effects are reversible. Science 339, 1621–5 (2013).

37. B. Naim, V. Brumfeld, R. Kapon, V. Kiss, R. Nevo, Z. Reich, Passive and facilitated transport in nuclear pore complexes is largely uncoupled. J Biol Chem 282, 3881–3888 (2007).

38. P. L. Paine, L. C. Moore, S. B. Horowitz, Nuclear envelope permeability. Nature 254, 109– 114 (1975).

39. S. Raffel, M. Falcone, N. Kneisel, J. Hansson, W. Wang, C. Lutz, L. Bullinger, G. Poschet, Y. Nonnenmacher, A. Barnert, C. Bahr, P. Zeisberger, A. Przybylla, M. Sohn, M. Tonjes, A. Erez, L. Adler, P. Jensen, C. Scholl, S. Frohling, S. Cocciardi, P. Wuchter, C. Thiede, A. Florcken, J. Westermann, G. Ehninger, P. Lichter, K. Hiller, R. Hell, C. Herrmann, A. D. Ho, J. Krijgsveld, B. Radlwimmer, A. Trumpp, BCAT1 restricts alphaKG levels in AML stem cells leading to IDHmut-like DNA hypermethylation. Nature 551, 384–388 (2017).

40. Y. Ma, Q. Zhu, J. Liang, Y. Li, M. Li, Y. Zhang, X. Wang, Y. Zeng, Y. Jiao, A CRISPR knockout negative screen reveals synergy between CDKs inhibitor and metformin in the treatment of human cancer in vitro and in vivo. Signal Transduct Target Ther 5, 152 (2020).

41. G.-D. Zhu, J. Yu, Z.-Y. Sun, Y. Chen, H.-M. Zheng, M.-L. Lin, S. Ou-Yang, G.-L. Liu, J.-W. Zhang, F.-M. Shao, Genome-wide CRISPR/Cas9 screening identifies CARHSP1 responsible for radiation resistance in glioblastoma. Cell Death Dis 12, 724 (2021).

42. A. Tsherniak, F. Vazquez, P. G. Montgomery, B. A. Weir, G. Kryukov, G. S. Cowley, S. Gill, W. F. Harrington, S. Pantel, J. M. Krill-Burger, R. M. Meyers, L. Ali, A. Goodale, Y. Lee, G. Jiang, J. Hsiao, W. F. J. Gerath, S. Howell, E. Merkel, M. Ghandi, L. A. Garraway, D. E. Root, T. R. Golub, J. S. Boehm, W. C. Hahn, Defining a Cancer Dependency Map. Cell 170, 564–576.e16 (2017).

43. K. Celis, S. Shuldiner, E. V. Haverfield, J. Cappell, R. Yang, D.-W. Gong, W. K. Chung, Loss of function mutation in glutamic pyruvate transaminase 2 (GPT2) causes developmental encephalopathy. J Inherit Metab Dis 38, 941–948 (2015).

44. Q. Ouyang, T. Nakayama, O. Baytas, S. M. Davidson, C. Yang, M. Schmidt, S. B. Lizarraga, S. Mishra, M. Ei-Quessny, S. Niaz, M. Gul Butt, S. Imran Murtaza, A. Javed, H. R. Chaudhry, D. J. Vaughan, R. S. Hill, J. N. Partlow, S.-Y. Yoo, A.-T. N. Lam, R. Nasir, M. Al-Saffar, A. J. Barkovich, M. Schwede, S. Nagpal, A. Rajab, R. J. DeBerardinis, D. E. Housman, G. H. Mochida, E. M. Morrow, Mutations in mitochondrial enzyme GPT2 cause metabolic dysfunction and neurological disease with developmental and progressive features. Proc Natl Acad Sci U S A 113, E5598–5607 (2016).

45. H. Kaymakcalan, Y. Yarman, N. Goc, F. Toy, C. Meral, A. G. Ercan-Sencicek, M. Gunel, Novel compound heterozygous mutations in GPT2 linked to microcephaly, and intellectual developmental disability with or without spastic paraplegia. Am J Med Genet A 176, 421– 425 (2018).

46. H. Hengel, R. Keimer, W. Deigendesch, A. Rieß, H. Marzouqa, J. Zaidan, P. Bauer, L. Schöls, GPT2 mutations cause developmental encephalopathy with microcephaly and features of complicated hereditary spastic paraplegia. Clin Genet 94, 356–361 (2018).

47. O. Baytas, S. M. Davidson, R. J. DeBerardinis, E. M. Morrow, Mitochondrial enzyme GPT2 regulates metabolic mechanisms required for neuron growth and motor function in vivo. Hum Mol Genet 31, 587–603 (2022).

48. O. Baytas, J. A. Kauer, E. M. Morrow, Loss of mitochondrial enzyme GPT2 causes early neurodegeneration in locus coeruleus. Neurobiol Dis 173, 105831 (2022).

49. 49. “Geigy scientific tables. 1: Units of measurement, body fluids, composition of the body, nutrition” (Ciba-Geigy, Basel, 8., rev.enl. ed., 1981).

50. O. Baytas, S. M. Davidson, J. A. Kauer, E. M. Morrow, Loss of mitochondrial enzyme GPT2 leads to reprogramming of synaptic glutamate metabolism. Mol Brain 17, 87 (2024).

51. J. Lüddecke, L. Francois, P. Spät, B. Watzer, T. Chilczuk, G. Poschet, R. Hell, B. Radlwimmer, K. Forchhammer, PII Protein-Derived FRET Sensors for Quantification and Live-Cell Imaging of 2-Oxoglutarate. Sci Rep 7, 1437 (2017).

52. T. Suzuki, M. Hayashi, T. Komatsu, A. Tanioka, M. Nagasawa, K. Tanimura-Inagaki, M. S. Rahman, S. Masuda, K. Yusa, J. Sakai, H. Shibata, T. Inagaki, Measurement of the nuclear concentration of α-ketoglutarate during adipocyte differentiation by using a fluorescence resonance energy transfer-based biosensor with nuclear localization signals. Endocr J 68, 1429–1438 (2021).

53. C. Zhang, Z.-H. Wei, B.-C. Ye, Quantitative monitoring of 2-oxoglutarate in Escherichia coli cells by a fluorescence resonance energy transfer-based biosensor. Appl Microbiol Biotechnol 97, 8307–8316 (2013).

54. C. Zhang, B.-C. Ye, A single fluorescent protein-based sensor for in vivo 2-oxogluatarate detection in cell. Biosens Bioelectron 54, 15–19 (2014).

55. J. Lüddecke, K. Forchhammer, From PII signaling to metabolite sensing: a novel 2- oxoglutarate sensor that details PII-NAGK complex formation. PLoS One 8, e83181 (2013).

56. H.-L. Chen, C. S. Bernard, P. Hubert, L. My, C.-C. Zhang, Fluorescence resonance energy transfer based on interaction of PII and PipX proteins provides a robust and specific biosensor for 2-oxoglutarate, a central metabolite and a signalling molecule. FEBS J 281, 1241–1255 (2014).

57. G. Sutendra, A. Kinnaird, P. Dromparis, R. Paulin, T. H. Stenson, A. Haromy, K. Hashimoto, N. Zhang, E. Flaim, E. D. Michelakis, A nuclear pyruvate dehydrogenase complex is important for the generation of acetyl-CoA and histone acetylation. Cell 158, 84–97 (2014).

58. S. Trefely, K. Huber, J. Liu, M. Noji, S. Stransky, J. Singh, M. T. Doan, C. D. Lovell, E. von Krusenstiern, H. Jiang, A. Bostwick, H. L. Pepper, L. Izzo, S. Zhao, J. P. Xu, K. C. Bedi, J. E. Rame, J. G. Bogner-Strauss, C. Mesaros, S. Sidoli, K. E. Wellen, N. W. Snyder, Quantitative subcellular acyl-CoA analysis reveals distinct nuclear metabolism and isoleucine-dependent histone propionylation. Mol Cell 82, 447–462.e6 (2022).

59. S. Li, S. K. Swanson, M. Gogol, L. Florens, M. P. Washburn, J. L. Workman, T. Suganuma, Serine and SAM Responsive Complex SESAME Regulates Histone Modification Crosstalk by Sensing Cellular Metabolism. Mol Cell 60, 408–421 (2015).

60. K. E. Wellen, G. Hatzivassiliou, U. M. Sachdeva, T. V. Bui, J. R. Cross, C. B. Thompson, ATP-citrate lyase links cellular metabolism to histone acetylation. Science 324, 1076–80 (2009).

61. S. Sivanand, S. Rhoades, Q. Jiang, J. V. Lee, J. Benci, J. Zhang, S. Yuan, I. Viney, S. Zhao, A. Carrer, M. J. Bennett, A. J. Minn, A. M. Weljie, R. A. Greenberg, K. E. Wellen, Nuclear Acetyl-CoA Production by ACLY Promotes Homologous Recombination. Mol Cell 67, 252–265.e6 (2017).

62. V. Bulusu, S. Tumanov, E. Michalopoulou, N. J. van den Broek, G. MacKay, C. Nixon, S. Dhayade, Z. T. Schug, J. Vande Voorde, K. Blyth, E. Gottlieb, A. Vazquez, J. J. Kamphorst, Acetate Recapturing by Nuclear Acetyl-CoA Synthetase 2 Prevents Loss of Histone Acetylation during Oxygen and Serum Limitation. Cell Rep 18, 647–658 (2017).

63. X. Li, W. Yu, X. Qian, Y. Xia, Y. Zheng, J.-H. Lee, W. Li, J. Lyu, G. Rao, X. Zhang, C.-N. Qian, S. G. Rozen, T. Jiang, Z. Lu, Nucleus-Translocated ACSS2 Promotes Gene Transcription for Lysosomal Biogenesis and Autophagy. Mol Cell 66, 684–697.e9 (2017).

64. S. Zhao, A. Torres, R. A. Henry, S. Trefely, M. Wallace, J. V. Lee, A. Carrer, A. Sengupta, S. L. Campbell, Y.-M. Kuo, A. J. Frey, N. Meurs, J. M. Viola, I. A. Blair, A. M. Weljie, C. M. Metallo, N. W. Snyder, A. J. Andrews, K. E. Wellen, ATP-Citrate Lyase Controls a Glucose-to-Acetate Metabolic Switch. Cell Rep 17, 1037–1052 (2016).

65. R. Nagaraj, M. S. Sharpley, F. Chi, D. Braas, Y. Zhou, R. Kim, A. T. Clark, U. Banerjee, Nuclear Localization of Mitochondrial TCA Cycle Enzymes as a Critical Step in Mammalian Zygotic Genome Activation. Cell 168, 210–223.e11 (2017).

66. E. Reytor, J. Pérez-Miguelsanz, L. Alvarez, D. Pérez-Sala, M. A. Pajares, Conformational signals in the C-terminal domain of methionine adenosyltransferase I/III determine its nucleocytoplasmic distribution. FASEB J 23, 3347–3360 (2009).

67. Y. Katoh, T. Ikura, Y. Hoshikawa, S. Tashiro, T. Ito, M. Ohta, Y. Kera, T. Noda, K. Igarashi, Methionine adenosyltransferase II serves as a transcriptional corepressor of Maf oncoprotein. Mol Cell 41, 554–66 (2011).

68. Y. Kera, Y. Katoh, M. Ohta, M. Matsumoto, T. Takano-Yamamoto, K. Igarashi, Methionine adenosyltransferase II-dependent histone H3K9 methylation at the COX-2 gene locus. J Biol Chem 288, 13592–13601 (2013).

69. B. Zhang, Y. Chen, L. Bao, W. Luo, GPT2 Is Induced by Hypoxia-Inducible Factor (HIF)-2 and Promotes Glioblastoma Growth. Cells 11, 2597 (2022).

70. A. Buffet, A. Morin, L.-J. Castro-Vega, F. Habarou, C. Lussey-Lepoutre, E. Letouzé, H. Lefebvre, I. Guilhem, M. Haissaguerre, I. Raingeard, M. Padilla-Girola, T. Tran, L. Tchara, J. Bertherat, L. Amar, C. Ottolenghi, N. Burnichon, A.-P. Gimenez-Roqueplo, J. Favier, Germline Mutations in the Mitochondrial 2-Oxoglutarate/Malate Carrier SLC25A11 Gene Confer a Predisposition to Metastatic Paragangliomas. Cancer Res 78, 1914–1922 (2018).

71. T. Mizoguchi, H. Minakuchi, M. Ishisaka, K. Tsuruma, M. Shimazawa, H. Hara, Behavioral abnormalities with disruption of brain structure in mice overexpressing VGF. Sci Rep 7, 4691 (2017).

72. D. J. Weisenberger, K. D. Siegmund, M. Campan, J. Young, T. I. Long, M. A. Faasse, G. H. Kang, M. Widschwendter, D. Weener, D. Buchanan, H. Koh, L. Simms, M. Barker, B. Leggett, J. Levine, M. Kim, A. J. French, S. N. Thibodeau, J. Jass, R. Haile, P. W. Laird, CpG island methylator phenotype underlies sporadic microsatellite instability and is tightly associated with BRAF mutation in colorectal cancer. Nat Genet 38, 787–793 (2006).

73. J. Klughammer, B. Kiesel, T. Roetzer, N. Fortelny, A. Nemc, K.-H. Nenning, J. Furtner, N. C. Sheffield, P. Datlinger, N. Peter, M. Nowosielski, M. Augustin, M. Mischkulnig, T. Ströbel, D. Alpar, B. Ergüner, M. Senekowitsch, P. Moser, C. F. Freyschlag, J. Kerschbaumer, C. Thomé, A. E. Grams, G. Stockhammer, M. Kitzwoegerer, S. Oberndorfer, F. Marhold, S. Weis, J. Trenkler, J. Buchroithner, J. Pichler, J. Haybaeck, S. Krassnig, K. Mahdy Ali, G. von Campe, F. Payer, C. Sherif, J. Preiser, T. Hauser, P. A. Winkler, W. Kleindienst, F. Würtz, T. Brandner-Kokalj, M. Stultschnig, S. Schweiger, K. Dieckmann, M. Preusser, G. Langs, B. Baumann, E. Knosp, G. Widhalm, C. Marosi, J. A. Hainfellner, A. Woehrer, C. Bock, The DNA methylation landscape of glioblastoma disease progression shows extensive heterogeneity in time and space. Nat Med 24, 1611–1624 (2018).

74. M. Fang, J. Ou, L. Hutchinson, M. R. Green, The BRAF oncoprotein functions through the transcriptional repressor MAFG to mediate the CpG Island Methylator phenotype. Mol Cell 55, 904–915 (2014).

75. Cancer Genome Atlas Research Network, Comprehensive molecular characterization of gastric adenocarcinoma. Nature 513, 202–209 (2014).

