## Supplemental Files for "A transcriptional biosensor reveals mechanisms of α-ketoglutarate signaling to chromatin"

**The PDF file includes:**

Materials and Methods
Figs. S1 to S12
Tables S1 to S5
References (*1-12*)

**Materials and Methods**
Experimental Model Details

Mice
All experiments involving live mice were carried out in accordance with the National Institutes of Health *Guide for the Care and Use of Laboratory Animals* under the protocols approved by the Brown University Institutional Animal Care and Use Committee. Gpt2 cryopreserved mutant embryos were established through resources at the Knockout Mouse Project at University of California, Davis (Project ID CSD24977). The background of the embryos is C57BL/6N, and these mice were fully backcrossed into a C57BL/6J line. The targeting construct involved a gene trap (splice acceptor) and *LacZ*-Neo cassette (*1*). The mice under the C57BL/6J background were then fully backcrossed onto the CD-1 line. Heterozygous males and heterozygous females were mated to produce offspring of both genders and all genotypes. Animals were genotyped by PCR with forward primer GPT2 neF (5'– TGATGGCACTCTGCACCTAC-3') and reverse primer GPT2 neR (5'– TCACTCTTGGCTCTGGACCT-3') for a WT band of 830 bp and a mutant band of 900 bp, as described previously (*1*). Mice were maintained on standard chow. On postnatal days 19 to 21 (P19-21), tissues including brain, heart, liver, kidney, lungs, spleen, stomach, colon, and sciatic nerve were harvested. All tissues were immediately snap-frozen and subsequently stored in liquid nitrogen until further analysis.

Cell lines
NHA cells (human astrocytes immortalized with HPV E6 and E7 and hTERT, sex unknown) were a kind gift of Dr. Russell Pieper (UCSF) (*2*). HEK293 cells (female) were purchased from ATCC (CRL-1573). HEK293T cells (female, transduced with SV40 T-antigen) were a kind gift of Dr. William G. Kaelin Jr. (Dana-Farber). U251 cells (male) were purchased from Sigma (09063001-1VL). Human TS516 glioblastoma stem-like cells (GSCs) (sex unknown) were obtained from Dr. Ingo Mellinghoff at MSKCC and reported previously (*3*).

All cells were cultured at 37°C, in the presence of ambient oxygen and 5% $CO_2$. HEK293, HEK293T, U251, and NHA cells were cultured in DMEM (Gibco 11995065) with 10% FBS and 1% penicillin/streptomycin unless otherwise stated. TS516 cells were cultured in NeuroCult NS-A Basal Medium (Human) with Proliferation Supplement (StemCell Technologies 05750) supplemented with EGF (20 ng/mL), bFGF (20 ng/mL), Heparin (2 mg/mL), 1% penicillin/streptomycin, amphotericin B (250 ng/mL), and Plasmocin (2.5 μg/mL).

All cell lines were routinely tested for mycoplasma contamination and were confirmed to be negative.

Methods

NtcA binding site prediction
Previously defined gene-specific NtcA binding sites were compared with the consensus NtcA binding site. The PRODORIC database of prokaryotic gene regulation was used to construct a position weight matrix defining the prevalence of nucleotides at each position within conserved NtcA binding sites derived from various cyanobacterial genomes. The PRODORIC database was accessed at: https://www.prodoric.de/.

Nuclear export signal prediction
The NetNES1.1 tool (*4*) was used to evaluate putative nuclear export signals (NES) throughout the primary amino acid sequence of the VP64-NtcA-$NLS_{c\text{-}myc}$ NtcA chimera. The NetNES1.1 tool was accessed at: https://services.healthtech.dtu.dk/services/NetNES-1.1/.

Mitochondrial targeting sequence prediction
Prediction of mitochondrial targeting sequences in protein products of GPT2 splice isoforms was conducted using TargetP-2.0. The TargetP-2.0 tool was accessed at: https://services.healthtech.dtu.dk/services/TargetP-2.0/.

NtcA structural analysis
Structural data describing the interaction between NtcA and DNA were derived from a cryo-electron microscopy (cryo-EM) study of the cyanobacterial transcription activation complex involving NtcA and NtcB (Protein Data Bank accession 8H3V). The NtcA-DNA complex was isolated from the larger transcription activation complex cryo-EM structure using Pymol software (Schrödinger).

Cloning
All synthetic DNA sequences, DNA plasmids, and CRISPR-Cas9 sgRNA sequences are listed in **table S2**, **table S3**, and **table S4**, respectively. In-Fusion cloning reactions were transformed into XL10-Gold Ultracompetent Cells (Agilent Technologies) following manufacturer's protocol. Gateway cloning reactions were performed with BP Clonase II (Invitrogen) or LR Clonase II (Invitrogen) enzymes. Gateway cloning reactions were transformed into HB101 Competent Cells (Promega). Ligation reactions were transformed into XL10-Gold Ultracompetent Cells (Agilent Technologies). PCR reactions were performed using KOD Xtreme Hot Start DNA Polymerase (EMD Millipore).

αKG-RE promoter/GFP reporter gene lentiviral vectors
First, sense and antisense ssDNAs comprising αKG-RE synthetic promoters were annealed to produce dsDNAs. In parallel, a $blasticidin^{resistance}$-P2A-GFP cDNA flanked 5' by a minimal CMV promoter was cloned into pEN_TTmcs (Addgene 25755) via In-Fusion cloning (In-Fusion HD Cloning Kit, Takara 639650) to produce the vector pEN_TTmcs-$P_{CMVmin}$- $blast^R$-P2A-GFP. Next, In-Fusion cloning was used to insert αKG-RE sequences into pEN_TTmcs-$P_{CMVmin}$-$blast^R$-P2A-GFP while removing the Tetracycline Response Element (TRE) from this vector. The resulting plasmids were referred to as pEN_TTmcs-[αKG-RE]-$P_{CMVmin}$- $blast^R$-P2A-GFP.

To accommodate αKG-RE promoter/GFP reporter gene expression in settings in which preservation of cellular blasticidin sensitivity was desired, the $blast^R$-P2A cassette 5' to the GFP cDNA was deleted by In-Fusion cloning, yielding pEN_TTmcs-[αKG-RE]-$P_{CMVmin}$-GFP vectors. For experiments testing the impact of GFP turnover on the dynamic range of the αKG-ON biosensor system, an hPEST domain was appended 3' to the GFP reporter, yielding pEN_TTmcs-[αKG-RE]-$P_{CMVmin}$- $blast^R$-P2A-GFP-hPEST vectors.

To create a promoterless Gateway lentiviral Destination vector, the PGK promoter was removed from pLenti PGK Neo DEST (w531-1) (Addgene 19067) by In-Fusion cloning to produce the vector pLenti Neo DEST. Finally, Gateway LR cloning reactions (Invitrogen Gateway LR clonase, Fisher Scientific 11-791-100) were performed between αKG-RE/GFP reporter gene-containing pEN_TTmcs vectors and the pLenti Neo DEST vector to generate lentiviral vectors for human cell transduction.

cDNA expression lentiviral vectors

cDNAs encoding NtcA chimeras or other proteins were produced via custom gene synthesis. They were cloned into lentiviral expression vectors via Gateway cloning reactions. Gateway destination vectors used for expression of NtcA chimeras and other cDNAs include: pLenti-Ubc-HA-Gate-PGK-HYG (Addgene 107396), pLenti PGK Hygro DEST (w530-1) (Addgene 19066), pLenti-EF1α-Gateway-PGK-Hygro (kind gift of Dr. William G. Kaelin, Jr.), and pLenti CMV Hygro DEST (w117-1) (Addgene 17454).

CRISPR-Cas9 lentiviral vectors

Cloning of sgRNAs (listed in **table S4**) into vectors was performed using FastDigest BsmBI (Esp3I) (Thermo Scientific) to digest plasmids. Oligonucleotides were annealed and phosphorylated using T4 PNK (NEB) in T4 Ligation Buffer (NEB). A ligation reaction with digested lentiGuide-puro (Addgene 52963), lentiCRISPR_v2-puro (Addgene 52961), lentiCRISPR_v2-blast (Addgene 83480), or lentiCRISPR_v2-mCherry (Addgene 99154) plasmid and the phosphorylated oligonucleotides was performed using Quick Ligase (NEB).

Stable cell line generation

Lentivirus was made using TransIT-LT1 Transfection Reagent (Mirus Bio MIR 2304). HEK293T cells were transfected by mixing TransIT-LT1 Transfection Reagent with expression vectors (listed in **table S3**) and packaging plasmids psPAX2 (Addgene 12260) and pMD2.G (Addgene 12259) in a ratio of 4:3:1. Virus-containing media were collected 48 and 72 hours after transfection and passed through a 0.45 μm filter to remove cell debris. Cells to be transduced were plated at a density of 150,000 – 300,000 cells per well in a 6-well plate. The next day, lentivirus was added with polybrene at a final concentration of 8 μg/mL. Cells were then incubated with lentivirus overnight before discarding lentivirus-containing medium and expanding the cells.

Antibiotic concentrations for selection of lentivirus-transduced cell cultures were as follows: puromycin (1-2 μg/mL), hygromycin (500 μg/mL), G418 (1 mg/mL), blasticidin (50 μg/mL), zeocin (50 μg/mL).

αKG-ON biosensor system expression

DNA plasmids used to express NtcA chimera and αKG-RE promoter/GFP reporter components of the αKG-ON biosensor system in each figure panel are listed in **table S5**.

CRISPR screening and analysis

U251 cells expressing the αKG-ON biosensor system were transduced with a Cas9 plasmid with a zeocin resistance marker. Lentivirus containing the custom "αKG Regulators" CRISPR-Cas9 library was infected at a multiplicity of infection (MOI) of ~0.3. 3E6 cells were infected per arm of the screen, with cells plated at a density of 1.5E6 cells per well of a 6-well plate. After transduction, cells were split into three replicates per arm and selected with puromycin. One week post-transduction, cells were plated at 250,000 cells per well (1.5E6 total per replicate per arm), and media was changed daily until day ten post-transduction, when cells were harvested for FACS. For each replicate of each arm, the top 10% highest GFP expressing cells, as well as the bottom 10% of GFP expressing cells, were sorted into separate tubes.

Genomic DNA was extracted from cells using the Blood & Tissue DNeasy kit (Qiagen). sgRNA cassettes were amplified and appended to Illumina primers by PCR using KOD Xtreme Hot Start DNA Polymerase (EMD Millipore). Amplicons were purified using SPRI Right Side Size Selection using a 0.7x ratio. Library QC was performed using a TapeStation instrument (Agilent). Library DNA content was quantified using a Qubit fluorometer (Thermo Fisher Scientific) before library sequencing. The sequencing results were then analyzed by MAGeCK (*5*) and Apron (Genetic Perturbation Platform, Broad Institute of MIT and Harvard).

Immunoblotting

Cells were lysed with EBC lysis buffer with protease inhibitor added (Roche). Nuclear and cytoplasmic fractionation was performed using the NE-PER Nuclear and Cytoplasmic Extraction Reagents (Thermo Scientific 78833). For histone western blots, cells were lysed using 2x loading buffer. Lysate protein concentrations were measured using the Bio-Rad Protein Assay Dye Reagent (#5000006). Extracted proteins were boiled at 100 °C for 10 min, subjected to polyacrylamide gel electrophoresis using the Mini-PROTEAN system (Bio-Rad), and transferred onto nitrocellulose membranes using the Mini Trans-Blot Cell (Bio-Rad). Primary antibodies (listed in **table S1**) were suspended in 5% BSA in TBST and secondary antibodies were diluted in 5% milk in TBST. Immobilon Western Chemiluminescent HRP substrate (EMD Millipore) was used for visualization and imaging was performed using an ImageQuant 800 biomolecular imager (Amersham). For densitometric analysis, Western blot images were scanned using Adobe Photoshop (Adobe Systems Inc.) and quantified with the gel analysis macros available in FIJI (NIH, https://imagej.net/software/fiji).

Flow cytometry and FACS

Flow cytometry analysis was performed using a FACS LSRFortessa from BD Biosciences. Cell sorting was performed using a FACS Aria II SORP (4 or 5 lasers) from BD Biosciences. Analysis of flow cytometry and FACS data was performed using FCS Express software from De Novo Software. Cells for analysis were suspended in a buffered solution with 50 ng/mL DAPI.

For initial experiments to determine the optimal transcription factor configuration, cells were plated at 150,000 cells per 6-well plate and transfected the next day (day one). Medium changes were performed each day after transfection, with cell harvest and analysis on day 4.

For cells transduced with an αKG-RE promoter/GFP reporter and NtcA chimera in a single vector, cells were plated at 125,000 cells per well in a 6-well plate followed by three days of media changes (glutamine deprivation as well as dm-αKG supplementation) before analysis.

For cells transduced with an αKG-RE promoter/GFP reporter and NtcA chimera in separate vectors, cells were plated at 250,000 cells per well in a 6-well plate and subjected to two days of media changes before analysis.

Immunofluorescence
500,000 HEK293 or U251 cells were plated on micro-well glass bottom plates with 20 mm micro-well #1.5 cover glass (Cellvis P06-20-1.5-N). Plates were pre-treated with 0.01% poly-L-ornithine solution for HEK293 cells. The next day, 100 nM Mito-Tracker Red CMXRos dye (Cell Signaling Technology 9082S) was added in 2 mL cell culture medium and incubated at 37°C for 20 minutes. 4% paraformaldehyde was added to the cells after removing media with MitoTracker Red, followed by 0.2% Triton-X 1000 treatment, and finally blocking with 0.2% fish skin gelatin (Sigma Aldrich G7041). Primary antibody was added in 0.2% fish skin gelatin and incubated overnight at 4 °C, including anti-FLAG Tag (1:800) or anti-HA Tag (1:1000) antibodies (listed in **table S1**). A 2 mg/mL stock of Goat anti-Mouse IgG (H+L) Cross-Adsorbed Secondary Antibody conjugated to Alexa Fluor 488 was diluted 1:2000 in 1 mL of 0.2% fish skin gelatin and added after primary antibody and incubated at 4 °C on a rocker for 2 hours. 300 nM DAPI in 0.2% fish skin gelatin was added for the last 5 minutes of incubation. Images were obtained in dual camera mode on a CSU-W1 spinning disk confocal microscope (Nikon).

Pyruvate quantification in blood and CSF
Absolute pyruvate concentrations in blood or CSF samples from children or adults were obtained from Geigy Scientific Tables, 1981 (*6*).

Liquid chromatography-mass spectrometry and stable isotope tracing
Relative quantification of 2HG (Figure S4G) was performed by plating cells at a density of 250,000 cells per well in a 6-well plate. The cells were subjected to two days of media changes with supplementation of (*R*)-2HG-TFMB or DMSO before collection and analysis.

Relative quantification of whole-cell αKG (Figures 2K and S8A-B) was performed by plating cells at a density of 250,000 cells per well in a 6-well plate. Cell culture media was changed the next day, two hours before harvesting the cells.

For stable isotope tracing experiments, cells were plated at 25,000 cells per well in 6-well plates. Cells were cultured in the presence or absence of 1 mM pyruvate for three days, with media changes each day. Two hours before harvest, media was changed again to glutamine-free DMEM supplemented 1) with 4 mM unlabeled glutamine or 4 mM $\alpha$-$^{15}$N-glutamine, and 2) with or without 1 mM pyruvate. Stable isotope tracing was conducted for 2 hours before harvesting cells.

All samples were washed twice with ice-cold saline and snap frozen in liquid nitrogen. Cells were harvested by scraping and metabolites were extracted directly in 80% acetonitrile (1 µL per 1,000 cells) and vortexed for 20 min at 4°C. Cells were then centrifuged at maximum speed for 10 minutes in a microcentrifuge at 4°C. The supernatant was harvested and centrifuged an additional time to remove debris. The final supernatant was injected and analyzed with a Q Exactive HF-X or Orbitrap Exploris hybrid quadrupole-orbitrap mass spectrometer (Thermo Fisher) coupled to a Vanquish Flex UHPLC system (Thermo Fisher). Chromatographic resolution of metabolites was achieved using a Millipore ZIC-pHILIC column using a linear gradient of 10 mM ammonium formate pH 9.8 and acetonitrile. Spectra were acquired with a resolving power of either 120,000 or 240,000 full width at half maximum (FWHM), a scan range set to 80–1,200 m/z, and polarity switching. Peaks were integrated using El-Maven 0.12.0

software (Elucidata). Total ion counts were quantified using TraceFinder 5.2 SP1 software (Thermo Fisher). Peaks were normalized to total ion counts using the R statistical programming language. For stable isotope tracing studies, correction for natural abundance of metabolite labeling was performed using the AccuCor package (*7*) in the R statistical programming language.

Gas chromatography-mass spectrometry

Quantification and analysis of steady state αKG levels in Figure S3A were performed using methods previously described (*8*). Briefly, cells were plated in 6-well plates at a density of 125,000 per well. Medium was changed daily for 3 days, and on day 4, cells were washed with ice-cold normal saline, and plates were snap frozen with liquid nitrogen. Metabolites were extracted by adding 350 µL ice-cold 70% methanol to each well. Cells were scraped off each well, and the cell suspension was transferred into an Eppendorf tube on dry ice. To each sample, 150 µL chloroform was added, and the samples were vortexed at 4 °C for 20 minutes, followed by centrifugation at 17,000 g for 10 minutes. The upper layer of methanol containing polar metabolites was transferred to a separate tube and dried overnight on a vacuum rotary evaporator (CentriVap, Labconco) at 4 °C. Samples not used immediately for analysis were stored at -80 °C.

Dried samples were derivatized by adding 20 µL methoxamine (MOX, Thermo Fisher 45950) per sample and vortexed for 20 minutes at 4°C, followed by incubation at 37°C for 1 hour. 30 µL N-tert-Butyldimethylsilyl-N-methyltrifluoroacetamide with 1% tert-Butyldimethylchlorosilane (TBDMS, Sigma 375934) was then added to each sample. The samples were vortexed briefly at room temperature and incubated for 1.5 hours at 65°C.

GC-MS analysis was performed on derivatized samples using an Agilent 7890B GC/5977A MSD system. Peak integration was performed using the Metran software tool (*9*). Relative metabolite quantification analyses were performed by normalizing metabolite ion counts to total ion counts measured within each sample.

DNA methylation

For analysis of U251 stable lines, cells were cultured in DMEM supplemented with 10% FBS and 1% penicillin-streptomycin as described in the cell culture methods. After transduction, cells were divided into two groups: one supplemented with 200 µM ascorbate-2-phosphate and the other without. The cells were maintained under these conditions for 2.5 weeks before harvesting.

Genomic DNA (gDNA) was extracted from cultured cells or brain/kidney tissues of Gpt2 WT or GPT2 KO mice using the Qiagen Blood & Tissue DNeasy kit, with RNase (NEB) added to remove RNA. The DNA was eluted in nuclease-free water and incubated with DNA Degradase Plus and Benzonase for four hours. Enzymatic reactions were quenched with methanol, and the samples were speed-dried. Finally, the dried samples were reconstituted in ammonium formate for UHPLC-MS/MS analysis. Samples were analyzed at the UT Southwestern Metabolic Phenotyping Core. 5 µL of sample was injected on a Nexera X2 UHPLC instrument coupled to an LCMS-8060 (Shimadzu Scientific Instruments, Columbia, MD, USA) triple quadrupole mass spectrometer using the electrospray ion source in positive mode. 5-methyldeoxycitidine (5mC), 5-hydroxymethyldeoxycitidine (5hmC), and deoxycytidine (C) were analyzed by selective reaction monitoring using the following transitions: 5mC 242 → 126, 5hmC 258 → 124 and 258

→ 142, C 228 → 112. Targeted nucleosides were resolved on a Shimadzu Cell Culture Profiling Column using a gradient of solvent B MeOH/MeCN (1:1, v/v) 0.1% formic acid over solvent A $H_2O$ 0.1 % formic acid at a 0.350 mL/min flow rate.

<u>Seahorse assays of mitochondrial function</u>
Oxygen consumption rate (OCR) was assessed using the Seahorse XF Cell Mito Stress Test Kit. 10,000 cells in each well were plated one day before the assay. Next day, the culture medium was replaced with OXPHOS assay medium, which consisted of DMEM without phenol red, supplemented with 2 mM glutamine, 1 mM sodium pyruvate, and 10 mM glucose, adjusted to pH 7.4. The plate was pre-incubated at 37°C for 1 hour in a non-$CO_2$ incubator. OCR measurements were initially taken under basal conditions, followed by sequential injections of specific reagents: 2 µM oligomycin, an inhibitor of Complex V that enables the calculation of mitochondrial ATP production; 1 µM carbonyl cyanide-p-trifluoromethoxyphenylhydrazone (FCCP), an uncoupling agent used to determine maximal respiration and spare capacity; and finally, 1 µM antimycin A, an inhibitor of Complex III, to halt mitochondrial respiration and allow the determination of non-mitochondrial respiration.

<u>Cell proliferation</u>
Cell proliferation assays were performed using a Celigo Image Cytometer (Revvity). Cells were plated in 96-well plates at a seeding density of 5,000 cells per well. To construct growth curves, cells were incubated with Hoechst 33342 (20 µM) and propidium iodide (PI) (1.0 µg/mL) dyes and viable cell numbers were measured each day from day 0 to day 4. To ensure robustness, we performed three independent replicates, each in technical triplicate, for each condition.

<u>Splice isoform analysis</u>
Paired FASTQ RNA-seq files were uploaded into UT Southwestern Medical Center's high performance computing system (BioHPC) on an established Nextflow (v20.01.0) platform for RNA-seq analysis called Astrocyte (v2.1.0). Reads were trimmed using TrimGalore (v0.4.1) and then aligned using HiSAT (v2.0.1). Picard (v1.127) was then used to mark and remove duplicate reads. Transcript-level splice isoform expression analysis was performed using the HiSAT, StringTie (v1.1.2), and Ballgown protocol which has been described (*10*). Ballgown is available through the Bioconductor suite through the R statistical environment (R v4.1.1). U251 RNA-seq data were obtained from the European Nucleotide Archive project (accession PRJEB3371) as well as from NCBI BioProject database (accession PRJNA631805) (*11*). Normal human astrocyte (NHA) RNA-seq data were obtained from NCBI (accession PRJNA631805) (*12*).

<u>RNA-seq</u>
Total RNA from 10 mg brain tissue samples was extracted using the RNeasy Mini Kit according to manufacturer's protocol (Qiagen 74004) and resuspended in 30 µl nuclease-free water. RNA-seq libraries were prepared by Novogene. The libraries were sequenced on an Illumina Novaseq X plus platform (Novogene).

<u>RNA-seq analysis</u>
FASTQ files were trimmed with TrimGalore (v0.6.10). Pre- and post-trimming quality control was done using FastQC (v0.12.1). RNA-seq data were aligned to the mouse reference genome (mm10) using HiSAT2 (v2.2.1). The SAM files were converted to BAM files by samtools

(v1.6). Then the transcript coverage of UCSC gene annotations were performed using featureCounts (v2.0.6). Bigwig files were created from BAM files using the bamCoverage function in deepTools. Gene ontology (GO) analysis was performed via the gene ontology platform (https://geneontology.org). Volcano plots were generated using ggplot2, displaying log-transformed *p* values on the y-axis and log2-transformed fold change values on the x-axis. Gene set enrichment analysis (GSEA) was conducted using GSEA software (v4.3.3) (https://www.gsea-msigdb.org/gsea).The normalized enrichment score (NES) and the false discovery rate (FDR) Q value were calculated by permuting gene set types, with a significance threshold of FDR $\leqslant$ 0.25 used to identify significantly enriched gene sets.

ChIP-seq

Brain tissues were disrupted with a TissueLyser instrument (Qiagen) then cross-linked with 1% formaldehyde in PBS for 10 minutes at room temperature. Cross-linking reactions were quenched with glycine for 5 minutes, followed by two washes in cold PBS. The crosslinked tissue was homogenized using a cooled blade homogenizer in homogenization buffer (50 mM Tris-HCl pH 7.5, 1% Nonidet P-40, 0.25% deoxycholic acid, 1 mM EDTA). Then, cells were lysed with Farnham Buffer (5 mM PIPES, pH 8.0, 85 mM KCl, 0.5% Nonidet P-40, 1 mM DTT, 0.1 mM PMSF) and followed by SDS Lysis Buffer (50 mM Tris-HCl pH 7.9, 10 mM EDTA, 1% SDS, 1 mM DTT) to obtain soluble chromatin.

Chromatin was sheared using a Covaris M220 Focused-ultrasonicator to generate DNA fragments approximately 200 to 400 bp in size, as verified by gel electrophoresis. The sonicated chromatin was centrifuged at maximum speed in a microcentrifuge for 1 minute at 4°C to remove insoluble debris. The soluble chromatin supernatant was diluted 1:9 with Dilution Buffer (20 mM Tris-HCl pH 7.9, 300 mM NaCl, 2 mM EDTA, 0.5% Triton X-100, 1 mM DTT, 0.2 mM PMSF) supplemented with a protease inhibitor cocktail (Roche). The diluted chromatin was precleared with 20 µL/mL protein A-agarose beads (ThermoFisher Scientific) on a nutator for 1 hour at 4°C. After centrifugation at 1000 x g for 3 minutes at 4°C, the supernatant was reserved for immunoprecipitation and 24 ng of spike-in chromatin (Active Motif 53083) was added per sample, with 2% set aside as input DNA. For immunoprecipitation, antibodies used included 5 µl H3K9me3 (Abcam AB8898 1063771-1) or 5 µl H3K4me3 (Active Motif 39159), along with 2 µg of spike-in antibody (Active Motif 61686), which were bound to 50 µl protein G Dynabeads (Invitrogen). The mixture was incubated with precleared supernatant overnight at 4°C with rotation. The Dynabeads were washed sequentially with the following buffers: low-salt wash buffer (10 mM Tris-HCl, pH 8, 2 mM EDTA, 0.1% SDS, 1% Triton X-100, 150 mM NaCl), high-salt wash buffer (10 mM Tris-HCl, pH 8, 2 mM EDTA, 0.1% SDS, 1% Triton X-100, 500 mM NaCl), LiCl wash buffer (10 mM Tris-HCl, pH 8, 1 mM EDTA, 1% NP-40, 1% Na-Deoxycholate, 250 mM LiCl), and a final wash with TE buffer containing 50 mM NaCl. Chromatin was eluted, incubated overnight at 65°C, and treated with RNase A and proteinase K. DNA was purified using the QIAquick PCR Purification Kit (Qiagen).

For library preparation, 10 ng of H3K9me3 ChIP DNA and 3 ng of H3K4me3 ChIP DNA were used with the NEBNext Ultra II DNA Library Preparation Kit for Illumina (New England Biolabs; NEB). Library quality was assessed using a High Sensitivity D5000 ScreenTape on an Agilent 2200 TapeStation and quantified with the Qubit dsDNA HS Assay Kit (Thermo Fisher). Libraries with unique adaptor barcodes were multiplexed and sequenced on an Illumina NextSeq

2000 platform (paired-end, 100 base pair reads), with a sequencing depth of 50 million reads for H3K9me3 ChIP and 30 million reads per sample for H3K4me3 ChIP.

ChIP-seq analysis

FASTQ files were trimmed with TrimGalore (v.0.6.10), and quality control was performed both before and after trimming using FastQC (v.0.12.1). ChIP-seq reads were aligned to the mm10 mouse reference genome using Bowtie2 (v.2.5.1). The SAM files were converted to BAM files by samtools (v.1.6). PCR duplicates were removed with PicardTools (v.3.0). We chose MACS2 (v.2.2.9.1) to call the peaks using the broad peaks setting for H3K9me3 (FDR $< 0.1$); we used the narrow peaks for H3K4me3 (FDR $< 0.1$) in the analysis. ChIP–seq peaks that were significantly increased or decreased in the Gpt2 KO mice were then identified using the DiffBind (v.3.16.0) package in R (v.4.2). Bigwig files for heatmap generation were created from BAM files using the bamCoverage function in deepTools. For visualization, representative track diagrams were generated using the Integrated Genomics Viewer (IGV) (v.2.9.4).

Ontology analysis of upregulated and H3K4 hypermethylated genes in Gpt2 KO versus Gpt2 WT mouse brain tissues

Genes (n = 169) displaying H3K4me3 hypermethylation and overexpression ($\log_2$FC $> 0.5$ and $p < 0.05$ for each analysis) in Gpt2 KO vs GPT2 WT mouse brain tissues were used to conduct a gene ontology (GO) analysis. The PANTHER database was queried to identify enriched biological processes in this gene set using a web tool made available through the Gene Ontology Consortium. The tool was accessed at: https://geneontology.org/.

Statistical analysis

Statistical analysis was performed using Graphpad Prism software. Figure legends include information on all statistical tests performed. $p$ values were calculated by unpaired $t$-tests for tests of statistical significance involving comparison of two groups. For tests of statistical significance involving comparison of three or more groups, $p$ values were calculated by one-way ANOVA tests. For analyses of H3K4me3 and H3K9me3 distributions in Fig. 5K and 5L, $p$ values were calculated by Kolmogorov-Smirnov tests. For all tests, $p$ values less than 0.05 were considered statistically significant. Where appropriate, statistically significant outlier data points were identified by Grubbs' test ($p < 0.05$) and excluded from data presentation.

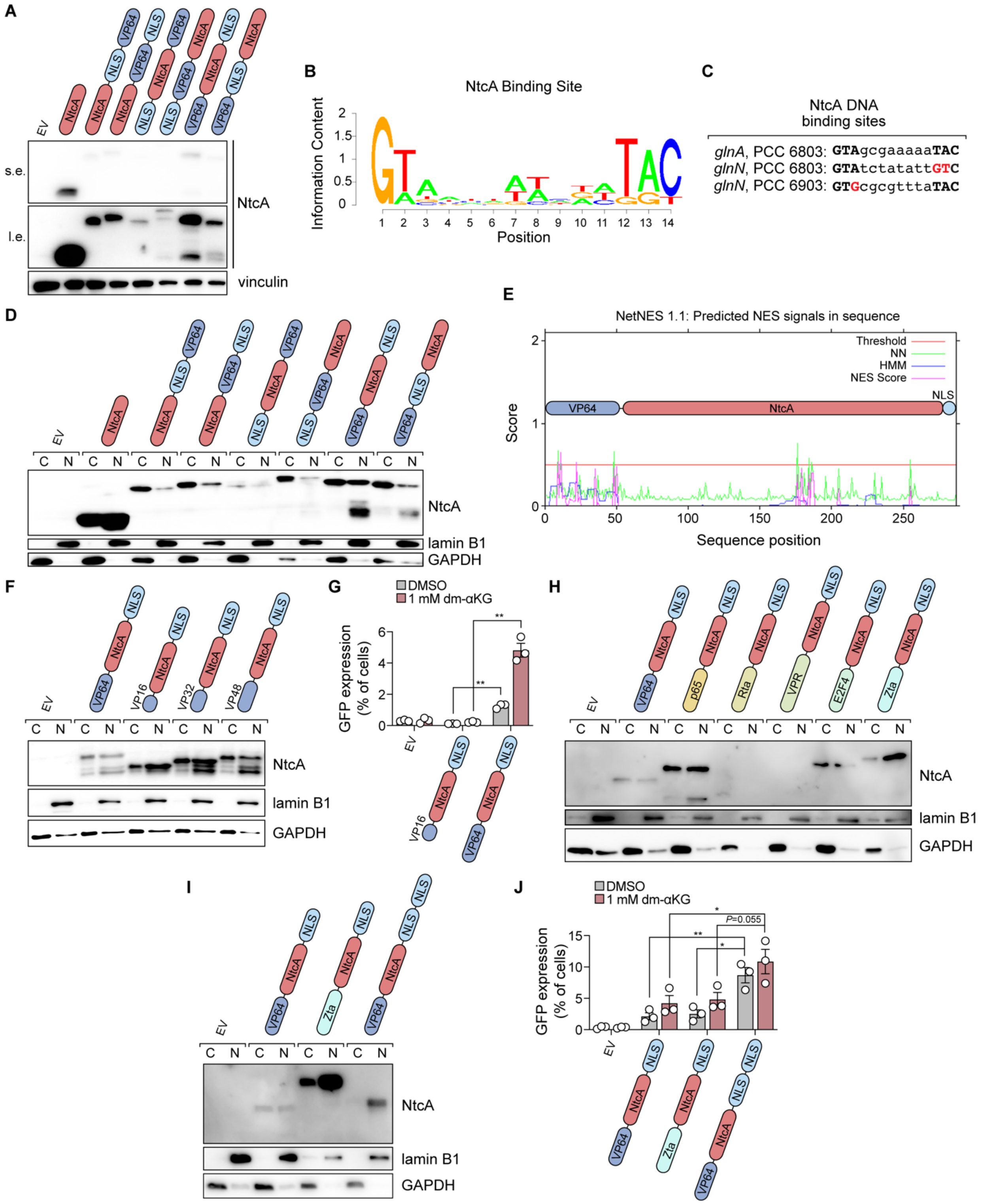
A
EV
NtcA
NLS
VP64
s.e.
l.e.
NtcA
vinculin
B
NtcA Binding Site
Information Content
Position
C
NtcA DNA binding sites
glnA, PCC 6803: GTAgcgaaaaaTAC
glnN, PCC 6803: GTAtctatattGTC
glnN, PCC 6903: GTGcgcgtttaTAC
D
C N
NtcA
lamin B1
GAPDH
E
NetNES 1.1: Predicted NES signals in sequence
Threshold
NN
HMM
NES Score
VP64
NtcA
NLS
Score
Sequence position
F
VP16
VP32
VP48
G
DMSO
1 mM dm-αKG
GFP expression (% of cells)
H
p65
Rta
VPR
E2F4
Zta
I
J
P=0.055

**Fig. S1. Optimization of the αKG-ON biosensor system.** (**A**) Immunoblot of NtcA proteins in HEK293 cells. NtcA chimeras constitute different configurations of VP64, NtcA, and $NLS_{c\text{-}myc}$ components. s.e. = short exposure; l.e. = long exposure. (**B**) Consensus NtcA binding site analysis. Position-specific nucleotide frequencies of NtcA binding sites from NtcA target genes in *Synechocystis* sp. PCC 6803 cyanobacteria. Data are from PRODORIC2 database. (**C**) Sequences of NtcA binding sites in promoters of indicated cyanobacterial genes. (**D**) Immunoblot of NtcA proteins in nuclear (N, lamin B1 marker) and cytosolic (C, GAPDH marker) fractions prepared from HEK293 cells. NtcA chimeras are identical to (A). (**E**) Predicted NES signals in VP64-NtcA-$NLS_{c\text{-}myc}$ chimera. (**F**) Immunoblot of NtcA chimeras in nuclear (N, lamin B1 marker) and cytosolic (C, GAPDH marker) fractions prepared from HEK293 cells. NtcA chimeras constitute the VP64-NtcA-$NLS_{c\text{-}myc}$ protein and various mutants featuring truncation of the VP64 domain. (**G**) Flow cytometry quantification of αKG-ON biosensor system activation (GFP expression) in αKG-RE-expressing HEK293 cells transfected with the VP16-NtcA-$NLS_{c\text{-}myc}$ or VP64-NtcA-$NLS_{c\text{-}myc}$ NtcA chimeras or EV and treated with DMSO or the indicated doses of dm-αKG for 48 hours. $^{**}p < 0.01$ (unpaired t-tests). (**H**) Immunoblot of NtcA proteins in nuclear (N, lamin B1 marker) and cytosolic (C, GAPDH marker) fractions prepared from HEK293 cells. NtcA chimeras constitute the VP64-NtcA-$NLS_{c\text{-}myc}$ protein and various mutants created by swapping the VP64 domain for alternative transactivation domains. (**I**) Immunoblot of NtcA proteins in nuclear (N, lamin B1 marker) and cytosolic (C, GAPDH marker) fractions prepared from HEK293 cells. NtcA chimeras constitute VP64-NtcA-$NLS_{c\text{-}myc}$, Zta-NtcA-$NLS_{c\text{-}myc}$, or VP64-NtcA-2x$NLS_{SV40}$ proteins. (**J**) Flow cytometry quantification of αKG-ON biosensor system activation (GFP expression) in αKG-RE-expressing HEK293 cells. Cells were transfected to express an NtcA chimera (VP64-NtcA-$NLS_{c\text{-}myc}$, Zta-NtcA-$NLS_{c\text{-}myc}$, VP64-NtcA-2x$NLS_{SV40}$) or EV and treated with DMSO or the indicated doses of dm-αKG for 24 hours. $^{*}p < 0.05$, $^{**}p < 0.01$ (unpaired t-tests). In (G and J), data are means ± SEM.

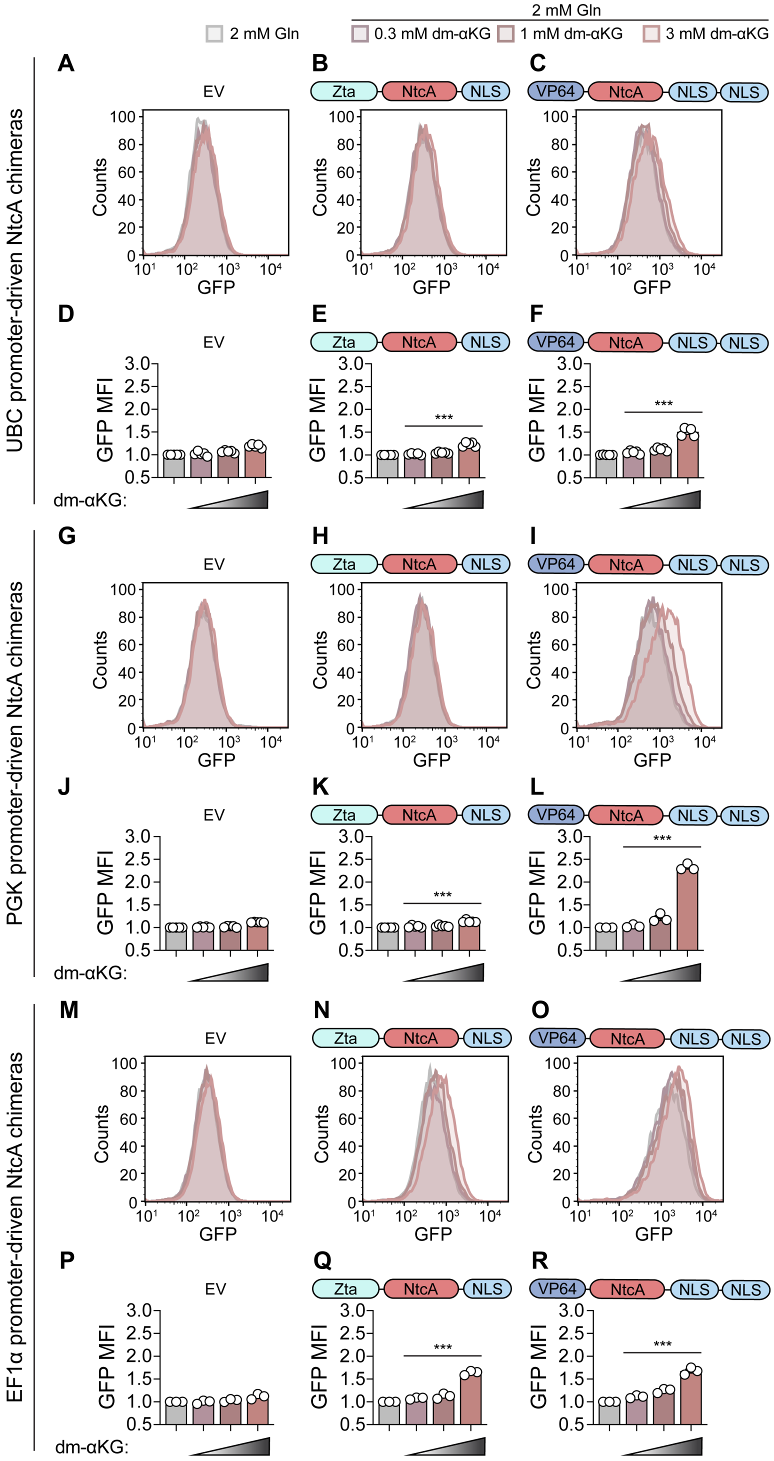
2 mM Gln
2 mM Gln
0.3 mM dm-αKG
1 mM dm-αKG
3 mM dm-αKG
UBC promoter-driven NtcA chimeras
PGK promoter-driven NtcA chimeras
EF1α promoter-driven NtcA chimeras
A
B
C
D
E
F
G
H
I
J
K
L
M
N
O
P
Q
R
EV
Zta
NtcA
NLS
VP64
Counts
GFP
GFP MFI
dm-αKG:
***

**Fig. S2. Testing the effect of NtcA chimera expression level on dynamic range of the αKG-ON biosensor system.** (**A-F**) Representative histograms (A-C) and quantification (D-F) of GFP expression in αKG-RE-expressing HEK293 cells transduced with EV or two different NtcA chimeras driven by the UBC promoter: Zta-NtcA-$NLS_{c\text{-}myc}$ or VP64-NtcA-2x$NLS_{SV40}$. Cells were cultured under the indicated concentrations of dm-αKG for 48 hours prior to analysis. In (D-F), data are GFP mean fluorescence intensity (MFI) values normalized to baseline conditions without dm-αKG. ***$p < 0.001$ (ordinary one-way ANOVA). (**G-L**) Representative histograms (G-I) and quantification (J-L) of GFP expression in αKG-RE-expressing HEK293 cells transduced with EV or two different NtcA chimeras driven by the PGK promoter: Zta-NtcA-$NLS_{c\text{-}myc}$ or VP64-NtcA-2x$NLS_{SV40}$. Cells were cultured under the indicated concentrations of dm-αKG for 48 hours prior to analysis. In (J-L), data are GFP mean fluorescence intensity (MFI) values normalized to baseline conditions without dm-αKG. ***$p < 0.001$ (ordinary one-way ANOVA). (**M-R**) Representative histograms (M-O) and quantification (P-R) of GFP expression in αKG-RE-expressing HEK293 cells transduced with EV or two different NtcA chimeras driven by the EF1α promoter: Zta-NtcA-$NLS_{c\text{-}myc}$ or VP64-NtcA-2x$NLS_{SV40}$. Cells were cultured under the indicated concentrations of dm-αKG for 48 hours prior to analysis. In (P-R), data are GFP mean fluorescence intensity (MFI) values normalized to baseline conditions without dm-αKG. ***$p < 0.001$ (ordinary one-way ANOVA). In (D-F, J-L, and P-R), data are means ± SEM.

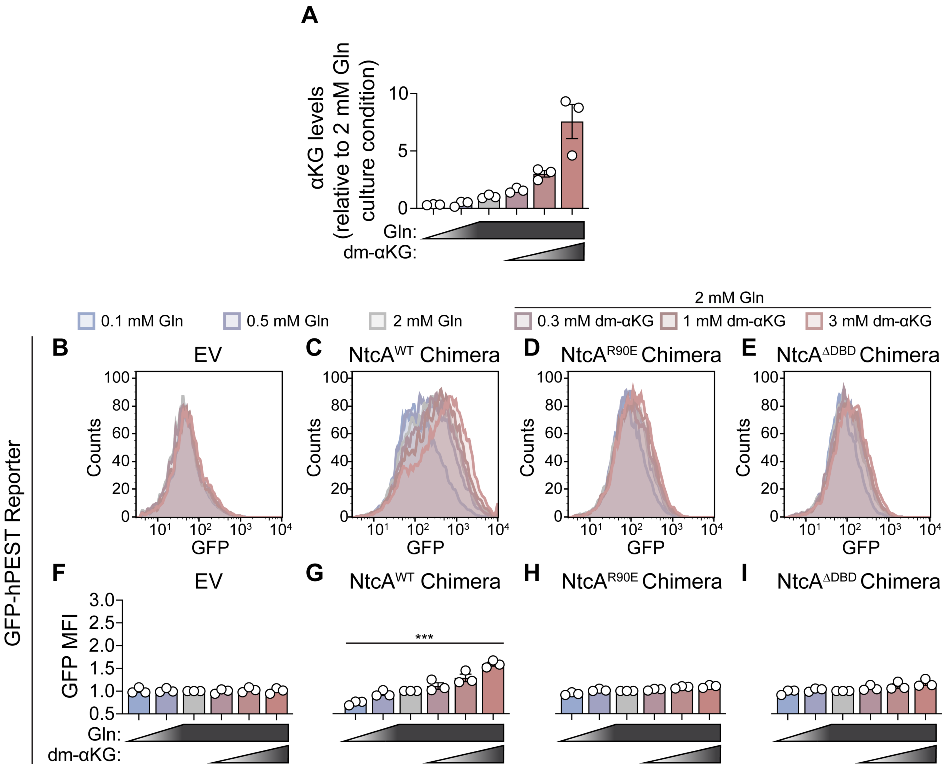
A
αKG levels (relative to 2 mM Gln culture condition)
Gln:
dm-αKG:
2 mM Gln
0.1 mM Gln
0.5 mM Gln
2 mM Gln
0.3 mM dm-αKG
1 mM dm-αKG
3 mM dm-αKG
GFP-hPEST Reporter
B
EV
C
NtcA^WT Chimera
D
NtcA^R90E Chimera
E
NtcA^ΔDBD Chimera
Counts
GFP
F
EV
G
NtcA^WT Chimera
H
NtcA^R90E Chimera
I
NtcA^ΔDBD Chimera
GFP MFI
***
Gln:
dm-αKG:

**Fig. S3. Perturbing cellular αKG pools regulates output of an αKG-ON biosensor system featuring an hPEST degron-fused GFP reporter.** (**A**) αKG levels in HEK293 cells. Cells were cultured under the indicated concentrations of glutamine and/or dm-αKG for 72 hours prior to analysis. (**B-I**) Representative histograms (B-E) and quantification (F-I) of GFP in HEK293 cells transduced with an αKG-RE/GFP-hPEST expression cassette and with EV or one of three VP64-NtcA-2xNLS$_{SV40}$ chimeras: wildtype (WT) NtcA, R90E NtcA mutant, DNA binding domain (ΔDBD)-deleted NtcA mutant. Cells were cultured under the indicated concentrations of glutamine and/or dm-αKG for 72 hours prior to analysis. In (F-I), data are GFP mean fluorescence intensity (MFI) values normalized to 2 mM Gln condition. ***$p < 0.001$ (ordinary one-way ANOVA). In (A and F-I), data are means ± SEM.

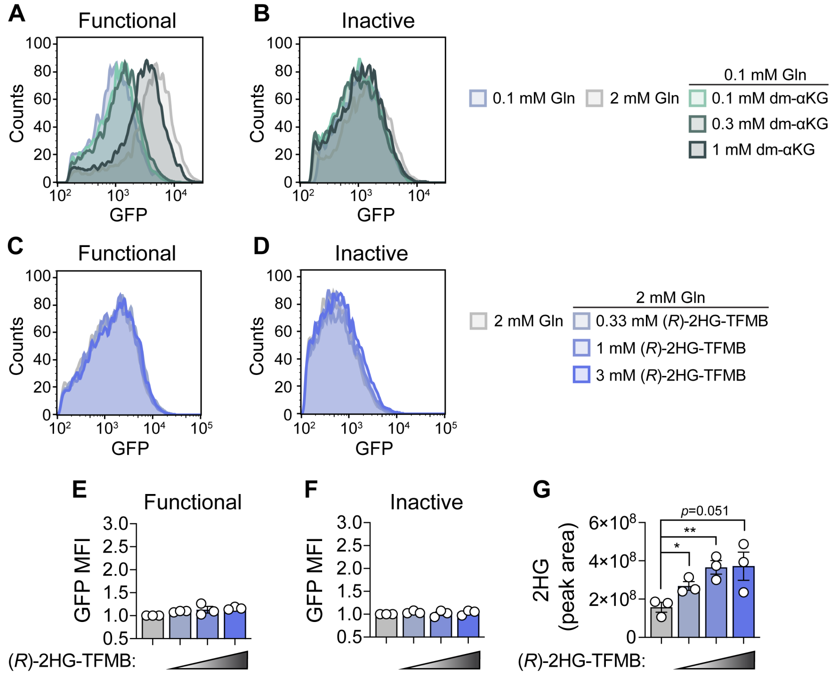

A
Functional
B
Inactive
Counts
GFP
0.1 mM Gln
2 mM Gln
0.1 mM Gln
0.1 mM dm-αKG
0.3 mM dm-αKG
1 mM dm-αKG
C
Functional
D
Inactive
2 mM Gln
2 mM Gln
0.33 mM (R)-2HG-TFMB
1 mM (R)-2HG-TFMB
3 mM (R)-2HG-TFMB
E
Functional
F
Inactive
GFP MFI
(R)-2HG-TFMB:
G
2HG (peak area)
p=0.051
**
*

**Fig. S4. The αKG-ON biosensor system specifically responds to altered nuclear αKG levels.** (**A-B**) Representative histograms of GFP expression in HEK293 cells transduced with an αKG-RE/GFP expression cassette and with one of two forms of VP64-NtcA-2xNLS$_{SV40}$ chimeras: wildtype (WT) NtcA or R90E NtcA mutant. Cells were cultured under the indicated concentrations of glutamine and/or dm-αKG for 72 hours prior to analysis. (**C-F**) Representative histograms (C-D) and quantification (E-F) of GFP expression in HEK293 cells engineered as in (A-B). Cells were cultured under the indicated concentrations of glutamine and/or (*R*)-2-HG-TFMB for 72 hours prior to analysis. In (E-F), data are GFP mean fluorescence intensity (MFI) values normalized to 2 mM Gln condition. (**G**) 2HG quantification in HEK293 cells transduced with an αKG-RE/GFP expression cassette and with the VP64-NtcA-2xNLS$_{SV40}$ chimera. Cells were cultured with DMSO or 0.3, 1, or 3 mM (*R*)-2HG-TFMB for 48 hours prior to analysis. $^*p < 0.05$, $^{**}p < 0.01$ (unpaired t-tests). In (E-G), data are means ± SEM.

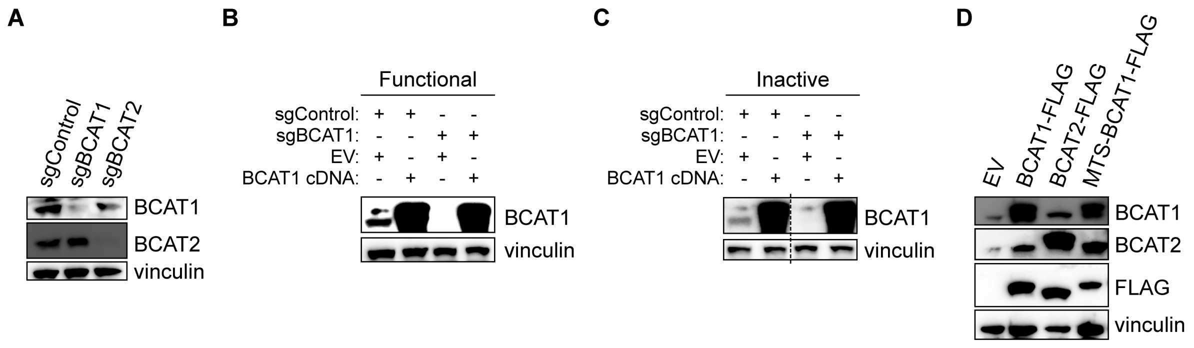

A
sgControl
sgBCAT1
sgBCAT2
BCAT1
BCAT2
vinculin
B
Functional
sgControl: + + - -
sgBCAT1: - - + +
EV: + - + -
BCAT1 cDNA: - + - +
BCAT1
vinculin
C
Inactive
sgControl: + + - -
sgBCAT1: - - + +
EV: + - + -
BCAT1 cDNA: - + - +
BCAT1
vinculin
D
EV
BCAT1-FLAG
BCAT2-FLAG
MTS-BCAT1-FLAG
BCAT1
BCAT2
FLAG
vinculin

**Fig. S5. Validating genetic manipulation of BCAT1 and BCAT2 expression.** (**A**) Immunoblots of BCAT1 and BCAT2 in HEK293 cells engineered to express Cas9 and control, BCAT1, or BCAT2 sgRNAs. (**B** and **C**) Immunoblots of BCAT1 expression in HEK293 cells engineered to express Cas9 and control or BCAT1 sgRNA, as well as an empty vector or sgRNA-resistant BCAT1 cDNA. HEK293 cells also expressed functional (B) or inactive (C) versions of the αKG-ON biosensor system. (**D**) Immunoblots of BCAT1, BCAT2, and FLAG in HEK293 cells engineered to express FLAG-tagged $BCAT1^{WT}$ enzyme, $BCAT2^{WT}$ enzyme, or a BCAT1 mutant enzyme with an N-terminal mitochondrial targeting signal (MTS) peptide.

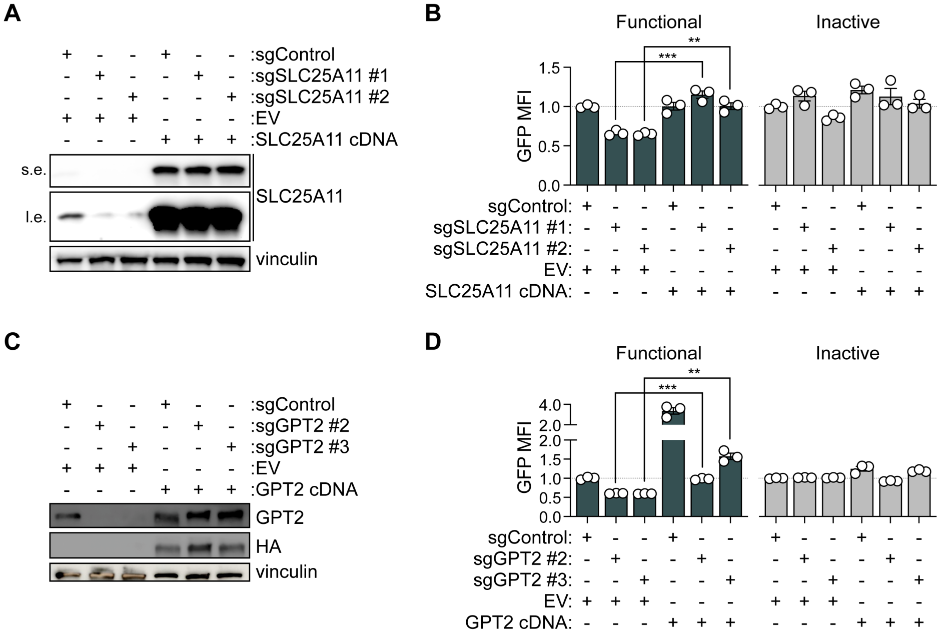
A
:sgControl
:sgSLC25A11 #1
:sgSLC25A11 #2
:EV
:SLC25A11 cDNA
s.e.
l.e.
SLC25A11
vinculin
B
Functional
Inactive
GFP MFI
sgControl:
sgSLC25A11 #1:
sgSLC25A11 #2:
EV:
SLC25A11 cDNA:
C
:sgControl
:sgGPT2 #2
:sgGPT2 #3
:EV
:GPT2 cDNA
GPT2
HA
vinculin
D
Functional
Inactive
GFP MFI
sgControl:
sgGPT2 #2:
sgGPT2 #3:
EV:
GPT2 cDNA:

**Fig. S6. Effects of SLC25A11- and GPT2-targeting sgRNAs on biosensor output are on-target.** (**A**) Immunoblots of SLC25A11 in U251 cells engineered to express Cas9 and control or SLC25A11 sgRNAs, as well as an empty vector or sgRNA-resistant SLC25A11 cDNA. s.e. = short exposure; l.e. = long exposure. (**B**) Quantification of GFP expression (GFP MFI normalized to sgControl, EV-expressing lines) in cells from (A) engineered with functional or inactive versions of the αKG-ON biosensor system. $^{**}p < 0.01$, $^{***}p < 0.001$ (unpaired t-tests). (**C**) Immunoblots of GPT2 and HA in U251 cells engineered to express Cas9 and control or GPT2 sgRNAs, as well as an empty vector or sgRNA-resistant GPT2 cDNA with HA tag. (**D**) Quantification of GFP expression (GFP MFI normalized to sgControl, EV-expressing lines) in cells from (C) engineered with functional or inactive versions of the αKG-ON biosensor system. $^{**}p < 0.01$, $^{***}p < 0.001$ (unpaired t-tests). In (B and D), data are means ± SEM.

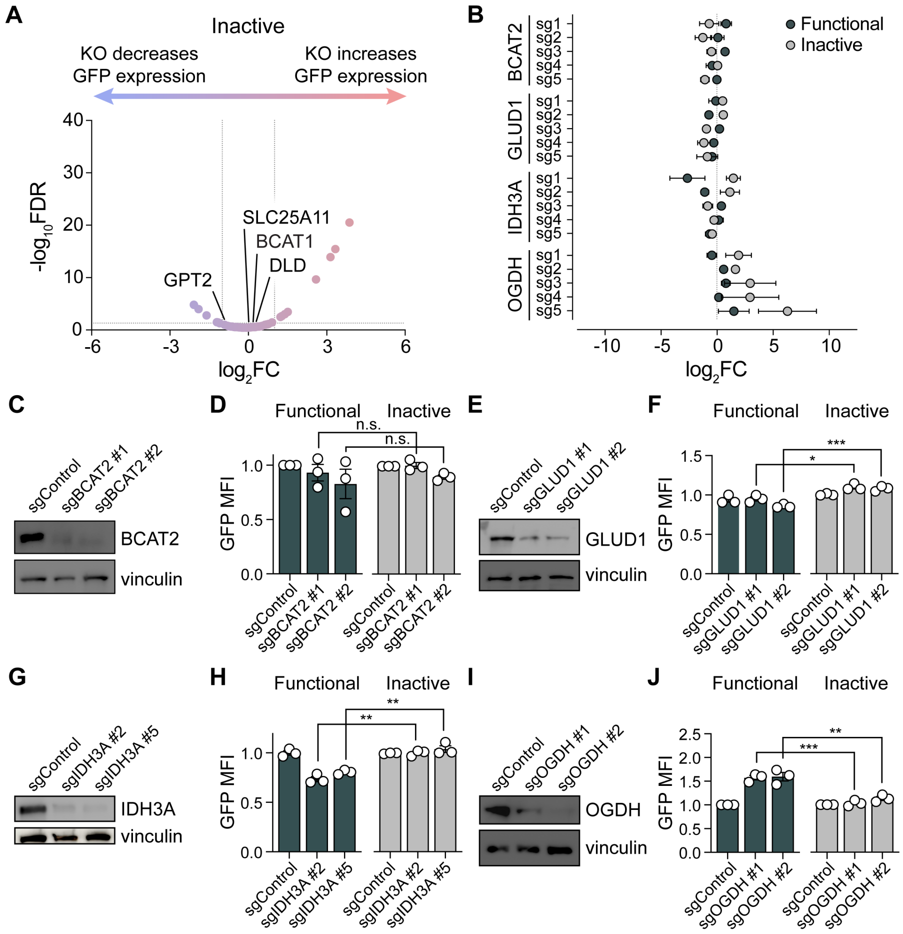

A
Inactive
KO decreases GFP expression
KO increases GFP expression
-log10FDR
log2FC
SLC25A11
BCAT1
DLD
GPT2
B
Functional
Inactive
BCAT2
GLUD1
IDH3A
OGDH
sg1
sg2
sg3
sg4
sg5
log2FC
C
sgControl
sgBCAT2 #1
sgBCAT2 #2
BCAT2
vinculin
D
Functional
Inactive
n.s.
n.s.
GFP MFI
E
sgControl
sgGLUD1 #1
sgGLUD1 #2
GLUD1
vinculin
F
Functional
Inactive
***
*
GFP MFI
G
sgControl
sgIDH3A #2
sgIDH3A #5
IDH3A
vinculin
H
Functional
Inactive
**
**
GFP MFI
I
sgControl
sgOGDH #1
sgOGDH #2
OGDH
vinculin
J
Functional
Inactive
**
***
GFP MFI

**Fig. S7. Evaluating genes with putative control over the nuclear αKG pool.** (**A**) Volcano plot of gene-level statistics from the CRISPR-Cas9 screen in U251 cells engineered with the inactive αKG-ON biosensor system shown in Fig. 3A. FC = fold change; FDR = false discovery rate. Gene-level scores were derived from the ratio of sgRNA read counts in the top 10% versus bottom 10% of GFP-expressing cells. (**B**) Enrichment or depletion of individual sgRNAs targeting *BCAT2*, *GLUD1*, *IDH3A* or *OGDH* genes from CRISPR-Cas9 screens in U251 cells engineered with functional or inactive versions of the αKG-ON biosensor system. (**C-K**) Assessment of nuclear αKG pool regulation by sgRNAs targeting *BCAT2*, *GLUD1*, *IDH3A* or *OGDH* genes. (C, E, G, and I) Immunoblots of BCAT2, GLUD1, IDH3A and OGDH expression in U251 cells engineered to express Cas9 and control or indicated sgRNAs. (D, F, H, and J) Quantification of GFP expression (GFP MFI normalized to sgControl lines) in cells from (C, E, G, and I) engineered with functional or inactive versions of the αKG-ON biosensor system. *$p < 0.05$, **$p < 0.01$, ***$p < 0.001$ (unpaired t-tests). In (B, D, F, H, and J), data are means ± SEM.

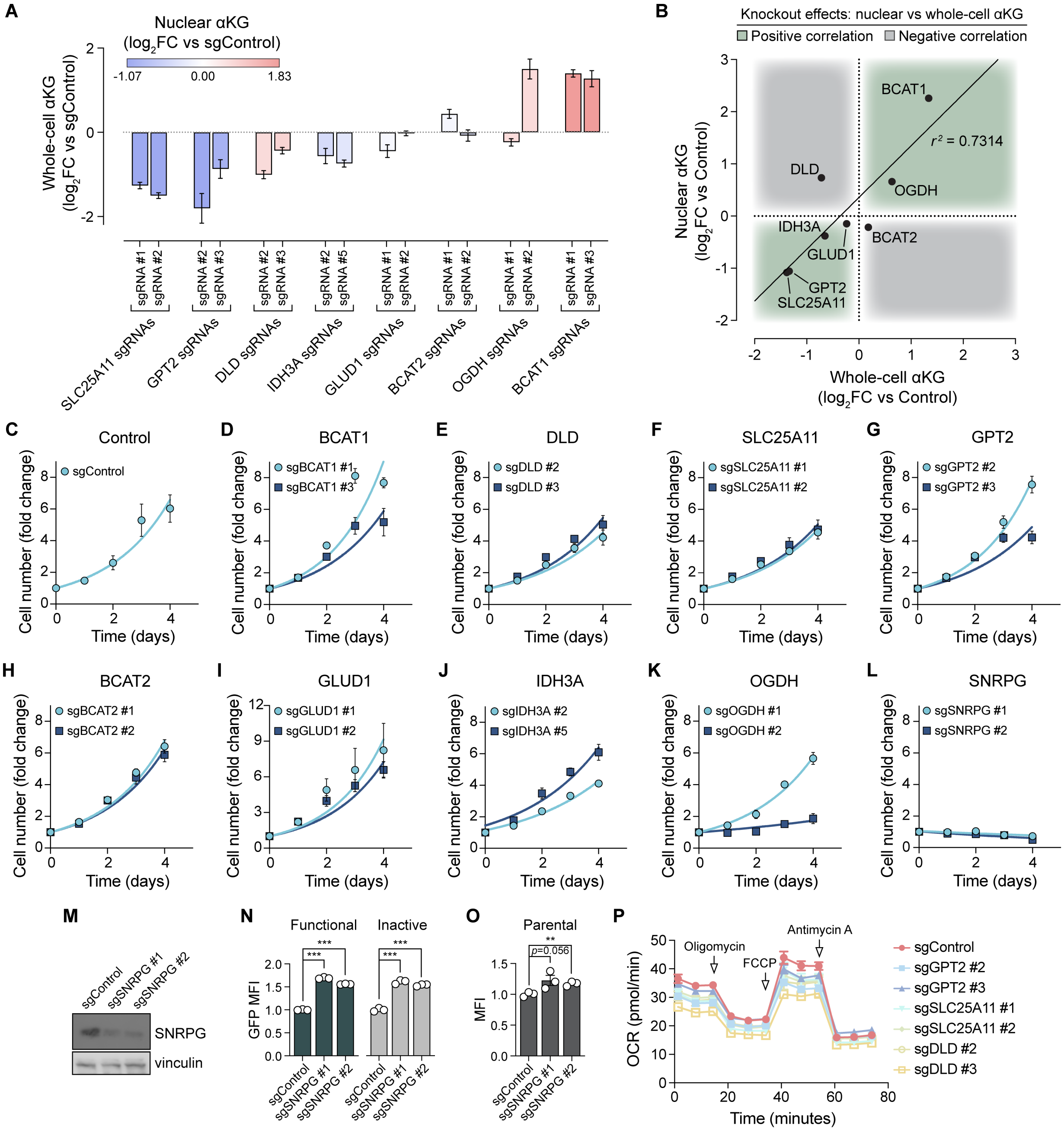

A
Nuclear αKG
(log2FC vs sgControl)
-1.07
0.00
1.83
Whole-cell αKG
(log2FC vs sgControl)
SLC25A11 sgRNAs
GPT2 sgRNAs
DLD sgRNAs
IDH3A sgRNAs
GLUD1 sgRNAs
BCAT2 sgRNAs
OGDH sgRNAs
BCAT1 sgRNAs
B
Knockout effects: nuclear vs whole-cell αKG
Positive correlation
Negative correlation
BCAT1
DLD
OGDH
IDH3A
GLUD1
BCAT2
GPT2
SLC25A11
r2 = 0.7314
Nuclear αKG
(log2FC vs Control)
Whole-cell αKG
(log2FC vs Control)
C
Control
sgControl
D
BCAT1
sgBCAT1 #1
sgBCAT1 #3
E
DLD
sgDLD #2
sgDLD #3
F
SLC25A11
sgSLC25A11 #1
sgSLC25A11 #2
G
GPT2
sgGPT2 #2
sgGPT2 #3
H
BCAT2
sgBCAT2 #1
sgBCAT2 #2
I
GLUD1
sgGLUD1 #1
sgGLUD1 #2
J
IDH3A
sgIDH3A #2
sgIDH3A #5
K
OGDH
sgOGDH #1
sgOGDH #2
L
SNRPG
sgSNRPG #1
sgSNRPG #2
Cell number (fold change)
Time (days)
M
sgControl
sgSNRPG #1
sgSNRPG #2
SNRPG
vinculin
N
Functional
Inactive
GFP MFI
O
Parental
p=0.056
MFI
P
Oligomycin
FCCP
Antimycin A
OCR (pmol/min)
Time (minutes)
sgControl
sgGPT2 #2
sgGPT2 #3
sgSLC25A11 #1
sgSLC25A11 #2
sgDLD #2
sgDLD #3

**Fig. S8. Metabolic and cell fitness effects of knockout of empirically determined or putative gene regulators of nuclear αKG. (A-B)** Quantification (A) and correlation (B) of changes in nuclear and whole-cell αKG levels upon knockout of indicated genes in U251 cells. **(C-L)** Quantification of U251 cell growth and viability (cell number normalized to day 0) in U251 cells engineered to express Cas9 and control, BCAT1, DLD, SLC25A11, GPT2, BCAT2, GLUD1, IDH3A, OGDH or SNRPG sgRNAs. (**M**) Immunoblots of SNRPG expression in U251 cells engineered to express Cas9 and control or SNRPG sgRNAs. (**N**) Quantification of GFP expression (GFP MFI normalized to sgControl lines) in cells from (M) engineered with functional or inactive versions of the αKG-ON biosensor system. ***$p < 0.001$ (unpaired t-tests). (**O**) Quantification of autofluorescence in the GFP channel (MFI normalized to sgControl line) displayed by U251 cells engineered to express Cas9 and control or SNRPG sgRNAs. U251 cells lacked an αKG-RE promoter/GFP reporter construct. **$p < 0.01$ (unpaired t-tests). (**P**) Oxygen consumption rates in U251 cells engineered to express Cas9 and control, GPT2, SLC25A11, or DLD sgRNAs. Mitochondrial inhibitors were injected at the indicated timepoints. In (A, C-L, and N-P), data are means ± SEM.

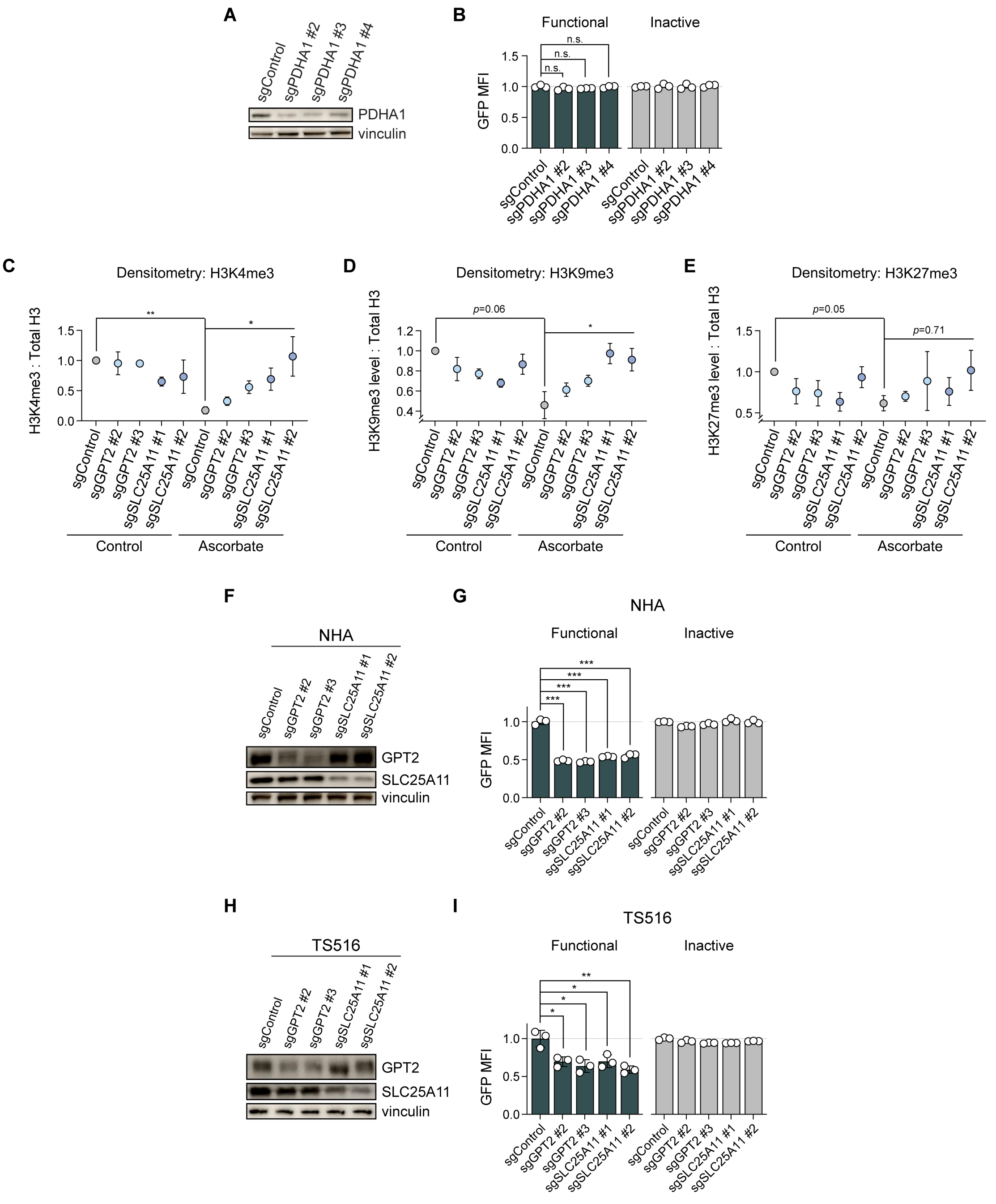
A
sgControl
sgPDHA1 #2
sgPDHA1 #3
sgPDHA1 #4
PDHA1
vinculin
B
Functional
Inactive
n.s.
GFP MFI
C
Densitometry: H3K4me3
H3K4me3 : Total H3
Control
Ascorbate
sgGPT2 #2
sgGPT2 #3
sgSLC25A11 #1
sgSLC25A11 #2
D
Densitometry: H3K9me3
H3K9me3 level : Total H3
p=0.06
E
Densitometry: H3K27me3
H3K27me3 level : Total H3
p=0.05
p=0.71
F
NHA
GPT2
SLC25A11
G
NHA
H
TS516
I
TS516

**Fig. S9. GPT2 or SLC25A11 knockout depletes nuclear αKG and impairs histone demethylation.** (**A**) Immunoblots of PDHA1 expression in U251 cells engineered to express Cas9 and control or PDHA1 sgRNAs. (**B**) Quantification of GFP expression (GFP MFI normalized to sgControl lines) in cells in cells from (A) engineered with functional or inactive versions of the αKG-ON biosensor system. n.s. = not significant (unpaired t-tests). (**C-E**) Quantification of histone methylation marks from immunoblots, including H3K4me3 (C), H3K9me3 (D), and H3K27me3 (E). Representative blots are shown in Fig. 4B. Lysates were prepared from U251 cells engineered to express Cas9 and control, GPT2, or SLC25A11 sgRNAs. Cells were treated with or without ascorbate-2-phosphate prior to harvest. Total histone H3 is a loading control. Densitometry was used to calculate the ratio of each methylation mark to total histone H3. Ratios were normalized to sgControl-expressing, untreated cells. *$p < 0.05$, **$p < 0.01$ (ordinary one-way ANOVA and unpaired t-tests). (**F**) Immunoblots of GPT2 and SLC25A11 expression in NHA cells engineered to express Cas9 and control, GPT2, or SLC25A11 sgRNAs. (**G**) Quantification of GFP expression (GFP MFI normalized to sgControl lines) in cells from (F) engineered with functional or inactive versions of the αKG-ON biosensor system. ***$p < 0.001$ (unpaired t-tests). (**H**) Immunoblots of GPT2 and SLC25A11 expression in TS516 cells engineered to express Cas9 and control, GPT2, or SLC25A11 sgRNAs. (**I**) Quantification of GFP expression (GFP MFI normalized to sgControl lines) in cells from (H) engineered with functional or inactive versions of the αKG-ON biosensor system. *$p < 0.05$, **$p < 0.01$ (unpaired t-tests). In (B-E, G and I), data are means ± SEM.

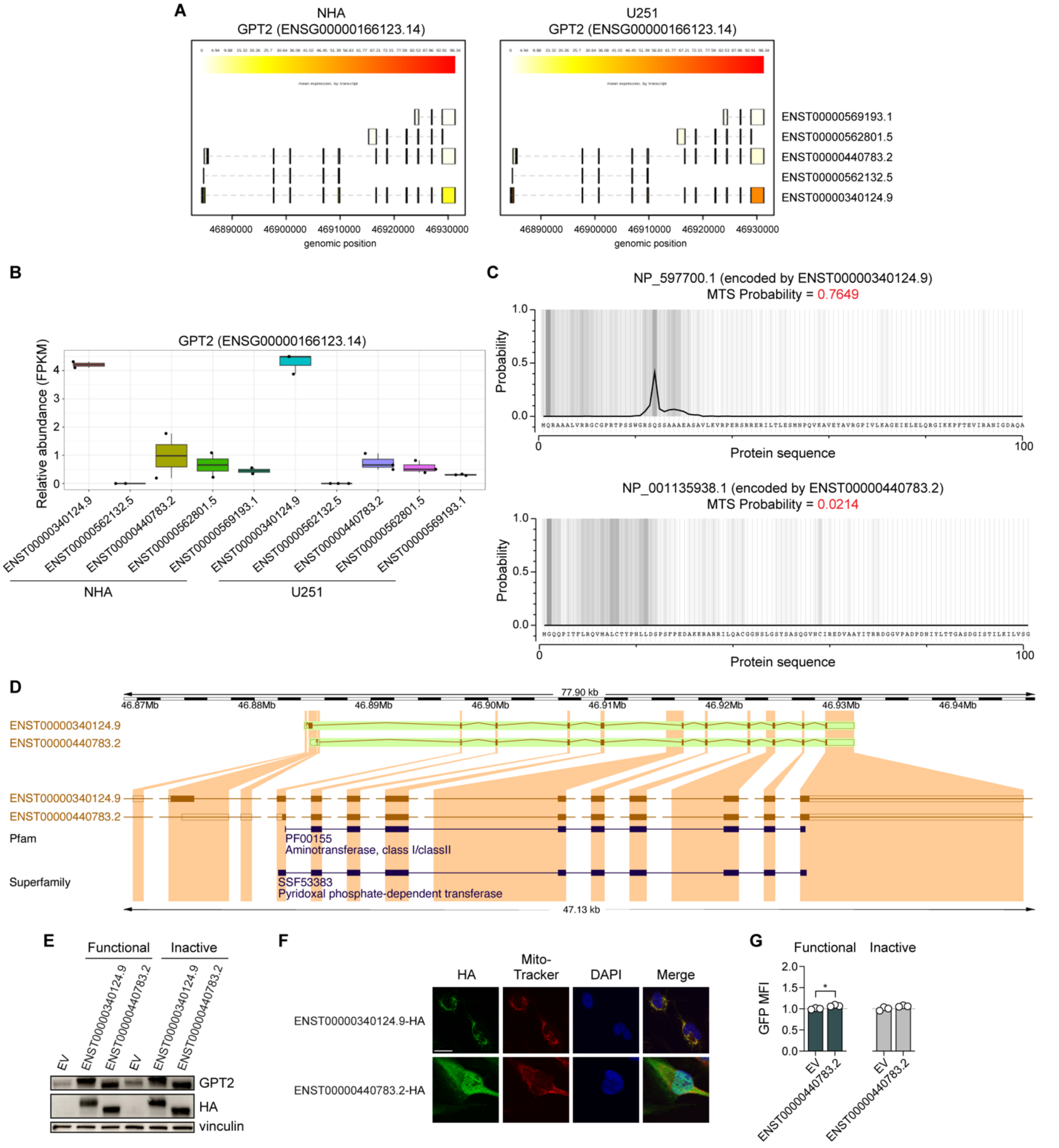

A
NHA
GPT2 (ENSG00000166123.14)
U251
GPT2 (ENSG00000166123.14)
ENST00000569193.1
ENST00000562801.5
ENST00000440783.2
ENST00000562132.5
ENST00000340124.9
46890000
46900000
46910000
46920000
46930000
genomic position
B
GPT2 (ENSG00000166123.14)
Relative abundance (FPKM)
NHA
U251
C
NP_597700.1 (encoded by ENST00000340124.9)
MTS Probability = 0.7649
NP_001135938.1 (encoded by ENST00000440783.2)
MTS Probability = 0.0214
Probability
Protein sequence
D
77.90 kb
46.87Mb
46.88Mb
46.89Mb
46.90Mb
46.91Mb
46.92Mb
46.93Mb
46.94Mb
ENST00000340124.9
ENST00000440783.2
Pfam
PF00155
Aminotransferase, class I/classII
Superfamily
SSF53383
Pyridoxal phosphate-dependent transferase
47.13 kb
E
Functional
Inactive
EV
ENST00000340124.9
ENST00000440783.2
GPT2
HA
vinculin
F
HA
Mito-Tracker
DAPI
Merge
ENST00000340124.9-HA
ENST00000440783.2-HA
G
Functional
Inactive
GFP MFI
EV
ENST00000440783.2

**Fig. S10. A truncated, extramitochondrial GPT2 isoform has negligible effects on the nuclear αKG pool.** (**A-B**) GPT2 transcript levels depicted as heatmaps (A) or quantified (B) in U251 and NHA cell lines. (**C**) Mitochondrial targeting sequence (MTS) prediction for GPT2 isoforms. Predicted probability of MTS signals in two GPT2 protein isoforms, NP_597700.1 (encoded by ENST00000340124.9) and NP_001135938.1 (encoded by ENST00000440783.2). (**D**) Genomic and protein domain organization of ENST00000340124.9 and ENST00000440783.2 GPT2 transcripts. (**E**) Immunoblots of GPT2 and HA expression in U251 cells engineered to express empty vector or HA-tagged ENST00000340124.9 or ENST00000440783.2 GPT2 transcripts. Cells also expressed wither functional or inactive versions of the αKG-ON biosensor system. (**F**) Representative immunofluorescence microscopy images of HA tag expression and Mito-Tracker or DAPI stains in U251 cells engineered to express HA-tagged ENST00000340124.9 or ENST00000440783.2 GPT2 transcripts. Merge shows overlay of all signals. Scale bar = 10 μm. (**G**) Relative nuclear αKG levels (GFP MFI normalized to EV line) in cells from (E). *$p < 0.05$ (unpaired t-test). In (G), data are means ± SEM.

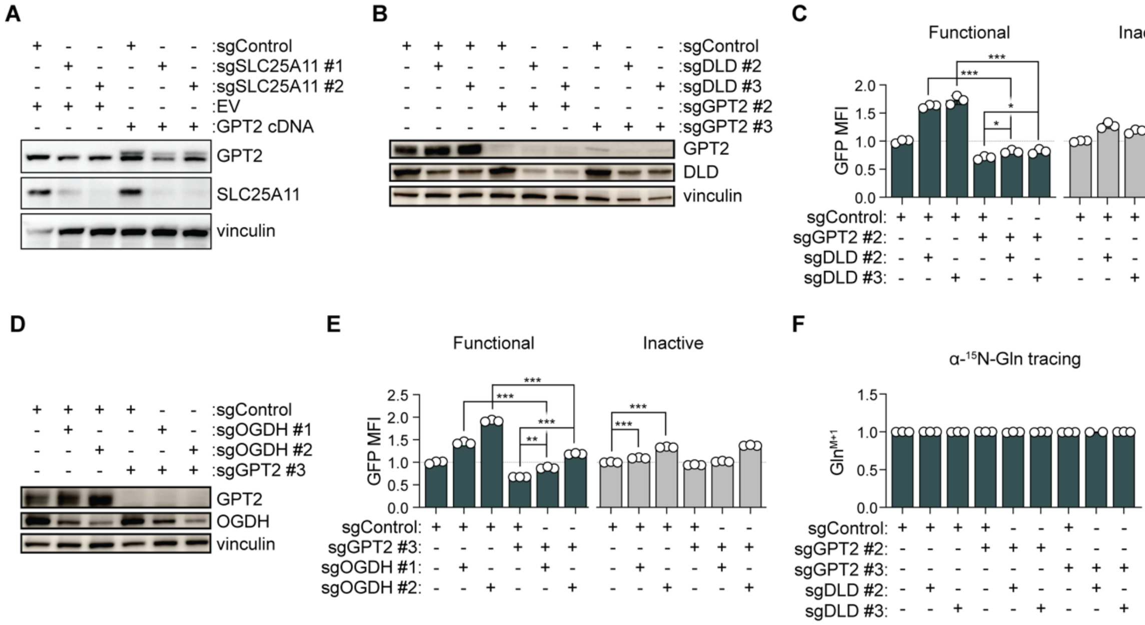
A
:sgControl
:sgSLC25A11 #1
:sgSLC25A11 #2
:EV
:GPT2 cDNA
GPT2
SLC25A11
vinculin
B
:sgControl
:sgDLD #2
:sgDLD #3
:sgGPT2 #2
:sgGPT2 #3
GPT2
DLD
vinculin
C
Functional
Inactive
GFP MFI
sgControl:
sgGPT2 #2:
sgDLD #2:
sgDLD #3:
D
:sgControl
:sgOGDH #1
:sgOGDH #2
:sgGPT2 #3
GPT2
OGDH
vinculin
E
Functional
Inactive
GFP MFI
sgControl:
sgGPT2 #3:
sgOGDH #1:
sgOGDH #2:
F
α-15N-Gln tracing
Gln M+1
sgControl:
sgGPT2 #2:
sgGPT2 #3:
sgDLD #2:
sgDLD #3:

**Fig. S11. Mechanistic dissection of the nuclear αKG regulatory network.** (**A**) Immunoblots of GPT2 and SLC25A11 expression in U251 cells engineered to express Cas9 and control or SLC25A11 sgRNAs, as well as empty vector or sgRNA-resistant GPT2 cDNA with an HA tag. (**B**) Immunoblots of GPT2 and DLD expression in U251 cells engineered to express Cas9 and control, GPT2, or DLD sgRNAs. (**C**) Quantification of GFP expression (GFP MFI normalized to sgControl lines) in cells from (B) engineered with functional or inactive versions of the αKG-ON biosensor system. $^{*}p < 0.05$, $^{***}p < 0.001$ (unpaired t-tests). (**D**) Immunoblots of GPT2 and OGDH expression in U251 cells engineered to express Cas9 and control, GPT2, or OGDH sgRNAs. (**E**) Quantification of GFP expression (GFP MFI normalized to sgControl lines) in cells from (D) engineered with functional or inactive versions of the αKG-ON biosensor system. $^{**}p < 0.01$, $^{***}p < 0.001$ (unpaired t-tests). (**F**) α-$^{15}$N-Glutamine stable isotope tracing for 2 hours in U251 cells engineered to express Cas9 and control, GPT2, or DLD sgRNAs. Fractional enrichment in the Gln (M+1) isotopologue is shown for each line. In (C, E, and F), data are means ± SEM.

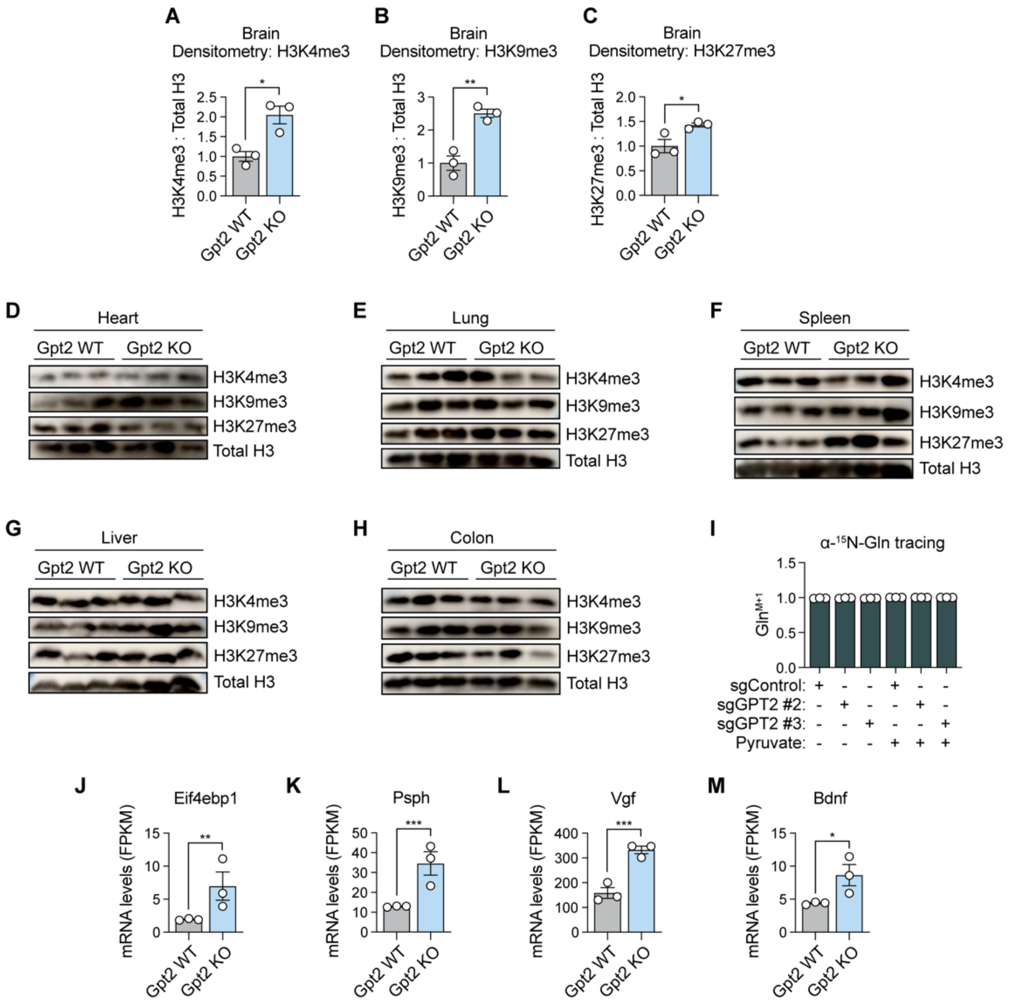

A
Brain
Densitometry: H3K4me3
H3K4me3 : Total H3
Gpt2 WT
Gpt2 KO
B
Brain
Densitometry: H3K9me3
H3K9me3 : Total H3
C
Brain
Densitometry: H3K27me3
H3K27me3 : Total H3
D
Heart
E
Lung
F
Spleen
G
Liver
H
Colon
H3K4me3
H3K9me3
H3K27me3
Total H3
I
α-15N-Gln tracing
sgControl:
sgGPT2 #2:
sgGPT2 #3:
Pyruvate:
J
Eif4ebp1
K
Psph
L
Vgf
M
Bdnf
mRNA levels (FPKM)

**Fig. S12. Organ-specific effects of Gpt2 knockout on histone methylation and gene expression.** (**A–C**) Quantification of histone methylation marks including H3K4me3 (A), H3K9me3 (B), and H3K27me3 (C) from blots shown in Fig. 5A. Blots were performed on brain tissue lysates from Gpt2 WT or Gpt2 KO mice, with total H3 as a loading control. Densitometry was used to calculate the ratio of each methylation mark to total histone H3. Ratios were normalized to Gpt2 WT samples. *$p < 0.05$, **$p < 0.01$ (unpaired t-tests). (**D-H**) Immunoblots of histone H3 total protein levels or trimethyllysine post-translational modifications in heart (D), lung (E), spleen (F), liver (G), or colon (H) tissues from Gpt2 WT or Gpt2 KO mice at P19-21. (**I**) $\alpha$-$^{15}$N-Glutamine stable isotope tracing for 2 hours in U251 cells engineered to express Cas9 and control or GPT2 sgRNAs and cultured with or without pyruvate. Fractional enrichment in the Gln (M+1) isotopologue is shown for each condition. (**J-M**) mRNA expression levels of *Eif4ebp1* (J), *Psph* (K), *Vgf* (L) and *Bdnf* (M) genes from brain tissues of Gpt2 WT or Gpt2 KO mice. *$p < 0.05$, **$p < 0.01$, ***$p < 0.01$ (unpaired t-tests). In (A-C and I-M), data are means ± SEM.

**Table S1. List of Antibodies**

| Antibody | Company | Cat. # | RRID | Application |
|---|---|---|---|---|
| NtcA | Gift from Javier Florencio/M. Isabel Muro Pastor | | | Immunoblotting |
| Lamin B1 | Proteintech | 66095-1 | AB_2721256 | Immunoblotting |
| GAPDH | Proteintech | 60004-1 | AB_2107436 | Immunoblotting |
| FLAG Tag | Cell Signaling | 8146S | AB_10950495 | Immunofluorescence |
| FLAG Tag | Millipore Sigma | F3165 | AB_259529 | Immunoblotting |
| HA Tag | BioLegend | 901501 | AB_2565006 | Immunoblotting, Immunofluorescence |
| BCAT1 | BD Biosciences | 611271 | AB_398799 | Immunoblotting |
| DLD | Proteintech | 16431-1-AP | AB_2091888 | Immunoblotting |
| GPT2 | Sigma Aldrich | HPA051514 | AB_2681516 | Immunoblotting |
| GPT2 | Thermo Fisher | PA5-62426 | AB_2642189 | Immunoblotting |
| SLC25A11 | Proteintech | 12253-1-AP | AB_2877840 | Immunoblotting |
| BCAT2 | Abcam | ab197917 | AB_2721840 | Immunoblotting |
| GLUD1 | Proteintech | 14299-1-AP | AB_2110515 | Immunoblotting |
| IDH3A | Proteintech | 15909-1-AP | AB_2123282 | Immunoblotting |
| OGDH | Proteintech | 15212-1-AP | AB_2156759 | Immunoblotting |
| SNRPG | Proteintech | 15084-1-AP | AB_2191789 | Immunoblotting |
| Vinculin | Sigma-Aldrich | V9131 | AB_477629 | Immunoblotting |
| PDHA1 | Abcam | ab110330 | AB_10858459 | Immunoblotting |
| H3K4me3 | Cell Signaling | 9751 | AB_2616028 | Immunoblotting |
| H3K9me3 | Cell Signaling | 13969 | AB_2798355 | Immunoblotting |
| H3K27me3 | Cell Signaling | 9733 | AB_2616029 | Immunoblotting |
| Histone H3 | Cell Signaling | 3638 | AB_1642229 | Immunoblotting |
| H3K4me3 (ChIP) | Active Motif | 39159 | AB_2615077 | ChIP |
| H3K9me3 (ChIP) | Abcam | ab8898 | AB_306848 | ChIP |
| Spike-in Antibody | Active Motif | 61686 | AB_2737370 | Spike-in for ChIP |
| Goat anti-mouse IgG, HRP-linked secondary antibody | Cell Signaling | 7076 | AB_330924 | Immunoblotting |
| Goat anti-rabbit IgG, HRP-linked secondary antibody | Cell Signaling | 7074 | AB_2099233 | Immunoblotting |

### Table S2. List of Synthetic DNA Sequences

| Synthetic DNA | Function | GenBank # |
|---|---|---|
| NtcA-NLS$_{c\text{-}myc}$-VP64 cDNA | NtcA chimera expression | Pending |
| NtcA-VP64-NLS$_{c\text{-}myc}$ cDNA | NtcA chimera expression | Pending |
| NLS$_{c\text{-}myc}$-NtcA-VP64 cDNA | NtcA chimera expression | Pending |
| NLS$_{c\text{-}myc}$-VP64-NtcA cDNA | NtcA chimera expression | Pending |
| VP64-NtcA-NLS$_{c\text{-}myc}$ cDNA | NtcA chimera expression | Pending |
| VP64-NLS$_{c\text{-}myc}$-NtcA cDNA | NtcA chimera expression | Pending |
| *glnA* PCC 6803 1x site with native sequence | αKG response element | Pending |
| *glnA* PCC 6803 5x sites with native spacers & sequence | αKG response element | Pending |
| *glnA* PCC 6803 5x sites with TRE spacers & sequence | αKG response element | Pending |
| *glnN* PCC 6803 1x site with native sequence | αKG response element | Pending |
| *glnN* PCC 6803 5x sites with native spacers & sequence | αKG response element | Pending |
| *glnN* PCC 6803 5x sites with TRE spacers & sequence | αKG response element | Pending |
| *glnA* PCC 6903 1x site with native sequence | αKG response element | Pending |
| *glnA* PCC 6903 5x sites with native spacers & sequence | αKG response element | Pending |
| *glnA* PCC 6903 5x sites with TRE spacers & sequence | αKG response element | Pending |
| VP16-NtcA-NLS$_{c\text{-}myc}$ cDNA | NtcA chimera expression | Pending |
| VP32-NtcA-NLS$_{c\text{-}myc}$ cDNA | NtcA chimera expression | Pending |
| VP48-NtcA-NLS$_{c\text{-}myc}$ cDNA | NtcA chimera expression | Pending |
| p65-NtcA-NLS$_{c\text{-}myc}$ cDNA | NtcA chimera expression | Pending |
| Rta-NtcA-NLS$_{c\text{-}myc}$ cDNA | NtcA chimera expression | Pending |
| VPR-NtcA-NLS$_{c\text{-}myc}$ cDNA | NtcA chimera expression | Pending |
| E2F4-NtcA-NLS$_{c\text{-}myc}$ cDNA | NtcA chimera expression | Pending |
| Zta-NtcA-NLS$_{c\text{-}myc}$ cDNA | NtcA chimera expression | Pending |
| VP64-NtcA-2xNLS$_{SV40}$ | NtcA chimera expression | Pending |
| VP64-NtcA$^{R90E}$-2xNLS$_{SV40}$ | NtcA chimera expression | Pending |
| VP64-NtcA$^{\Delta DBD}$-2xNLS$_{SV40}$ | NtcA chimera expression | Pending |
| mCherry-T2A-VP64-NtcA-2xNLS$_{SV40}$ | NtcA chimera expression | Pending |
| mCherry-T2A-VP64-NtcA$^{R90E}$-2xNLS$_{SV40}$ | NtcA chimera expression | Pending |
| MTS-BCAT1-FLAG | BCAT1 mutant expression | Pending |

### Table S3. List of DNA Plasmids

| ID | Insert | Vector | Expression |
|---|---|---|---|
| 1 | *glnA* PCC 6803 1x site with native sequence | pLenti Neo DEST $P_{CMVmin}$-blast$^{R}$-P2A-GFP | Lentiviral |
| 2 | *glnA* PCC 6803 5x sites with native spacers & sequence | pLenti Neo DEST $P_{CMVmin}$-blast$^{R}$-P2A-GFP | Lentiviral |
| 3 | *glnA* PCC 6803 5x sites with TRE spacers & sequence | pLenti Neo DEST $P_{CMVmin}$-blast$^{R}$-P2A-GFP | Lentiviral |
| 4 | *glnN* PCC 6803 1x site with native sequence | pLenti Neo DEST $P_{CMVmin}$-blast$^{R}$-P2A-GFP | Lentiviral |
| 5 | *glnN* PCC 6803 5x sites with native spacers & sequence | pLenti Neo DEST $P_{CMVmin}$-blast$^{R}$-P2A-GFP | Lentiviral |
| 6 | *glnN* PCC 6803 5x sites with TRE spacers & sequence | pLenti Neo DEST $P_{CMVmin}$-blast$^{R}$-P2A-GFP | Lentiviral |
| 7 | *glnN* PCC 6903 1x site with native sequence | pLenti Neo DEST $P_{CMVmin}$-blast$^{R}$-P2A-GFP | Lentiviral |
| 8 | *glnN* PCC 6903 5x sites with native spacers & sequence | pLenti Neo DEST $P_{CMVmin}$-blast$^{R}$-P2A-GFP | Lentiviral |
| 9 | *glnN* PCC 6903 5x sites with TRE spacers & sequence | pLenti Neo DEST $P_{CMVmin}$-blast$^{R}$-P2A-GFP | Lentiviral |
| 10 | EV | pEZY3 | Transient |
| 11 | NtcA-NLS$_{c\text{-}myc}$-VP64 | pEZY3 | Transient |
| 12 | NtcA-VP64-NLS$_{c\text{-}myc}$ | pEZY3 | Transient |
| 13 | NLS$_{c\text{-}myc}$-NtcA-VP64 | pEZY3 | Transient |
| 14 | NLS$_{c\text{-}myc}$-VP64-NtcA | pEZY3 | Transient |
| 15 | VP64-NtcA-NLS$_{c\text{-}myc}$ | pEZY3 | Transient |
| 16 | VP64-NLS$_{c\text{-}myc}$-NtcA | pEZY3 | Transient |
| 17 | VP64-NtcA-2xNLS$_{SV40}$ | pEZY3 | Transient |
| 18 | EV | pLenti-EF1α-Gateway-PGK-Hygro | Lentiviral |
| 19 | VP64-NtcA-2xNLS$_{SV40}$ | pLenti-EF1α-Gateway-PGK-Hygro | Lentiviral |
| 20 | pEZY3 | NtcA | Transient |
| 21 | VP16-NtcA-NLS$_{c\text{-}myc}$ | pEZY3 | Transient |
| 22 | VP32-NtcA-NLS$_{c\text{-}myc}$ | pEZY3 | Transient |
| 23 | VP48-NtcA-NLS$_{c\text{-}myc}$ | pEZY3 | Transient |
| 24 | p65-NtcA-NLS$_{c\text{-}myc}$ | pEZY3 | Transient |
| 25 | Rta-NtcA-NLS$_{c\text{-}myc}$ | pEZY3 | Transient |
| 26 | VPR-NtcA-NLS$_{c\text{-}myc}$ | pEZY3 | Transient |
| 27 | E2F4-NtcA-NLS$_{c\text{-}myc}$ | pEZY3 | Transient |
| 28 | Zta-NtcA-NLS$_{c\text{-}myc}$ | pEZY3 | Transient |
| 29 | *glnN* PCC 6903 5x sites with TRE spacers & sequence + VP64-NtcA-2xNLS$_{SV40}$ | pLenti Neo DEST $P_{CMVmin}$-blast$^{R}$-P2A-GFP | Lentiviral |
| 30 | *glnN* PCC 6903 5x sites with TRE spacers & sequence + VP64-NtcA$^{R90E}$-2xNLS$_{SV40}$ | pLenti Neo DEST $P_{CMVmin}$-blast$^{R}$-P2A-GFP | Lentiviral |
| 31 | *glnN* PCC 6903 5x sites with TRE spacers & sequence + VP64-NtcA$^{\Delta DBD}$-2xNLS$_{SV40}$ | pLenti Neo DEST $P_{CMVmin}$-blast$^{R}$-P2A-GFP | Lentiviral |
| 32 | *glnN* PCC 6903 5x sites with TRE spacers & sequence + mCherry-T2A-VP64-NtcA-2xNLS$_{SV40}$ | pLenti Neo DEST $P_{CMVmin}$-blast$^{R}$-P2A-GFP | Lentiviral |
| 33 | *glnN* PCC 6903 5x sites with TRE spacers & sequence + mCherry-T2A-VP64-NtcA$^{R90E}$-2xNLS$_{SV40}$ | pLenti Neo DEST $P_{CMVmin}$-blast$^{R}$-P2A-GFP | Lentiviral |
| 34 | EV | pLenti-Ubc-HA-Gate-PGK-HYG | Lentiviral |
| 35 | Zta-NtcA-NLS$_{c\text{-}myc}$ | pLenti-Ubc-HA-Gate-PGK-HYG | Lentiviral |

| | | | |
|---|---|---|---|
| 36 | VP64-NtcA-2xNLS$_{SV40}$ | pLenti-Ubc-HA-Gate-PGK-HYG | Lentiviral |
| 37 | EV | pLenti PGK Hygro DEST (w530-1) | Lentiviral |
| 38 | Zta-NtcA-NLS$_{c\text{-}myc}$ | pLenti PGK Hygro DEST (w530-1) | Lentiviral |
| 39 | VP64-NtcA-2xNLS$_{SV40}$ | pLenti PGK Hygro DEST (w530-1) | Lentiviral |
| 40 | EV | pLenti-EF1α-Gateway-PGK-Hygro | Lentiviral |
| 41 | Zta-NtcA-NLS$_{c\text{-}myc}$ | pLenti-EF1α-Gateway-PGK-Hygro | Lentiviral |
| 42 | VP64-NtcA-2xNLS$_{SV40}$ | pLenti-EF1α-Gateway-PGK-Hygro | Lentiviral |
| 43 | *glnN* PCC 6903 5x sites with TRE spacers & sequence | pLenti Neo DEST $P_{CMVmin}$-blast$^R$-P2A-GFP-hPEST | Lentiviral |
| 44 | *glnN* PCC 6903 5x sites with TRE spacers & sequence + VP64-NtcA-2xNLS$_{SV40}$ | pLenti Neo DEST $P_{CMVmin}$-blast$^R$-P2A-GFP-hPEST | Lentiviral |
| 45 | *glnN* PCC 6903 5x sites with TRE spacers & sequence + VP64-NtcA$^{R90E}$-2xNLS$_{SV40}$ | pLenti Neo DEST $P_{CMVmin}$-blast$^R$-P2A-GFP-hPEST | Lentiviral |
| 46 | *glnN* PCC 6903 5x sites with TRE spacers & sequence + VP64-NtcA$^{\Delta DBD}$-2xNLS$_{SV40}$ | pLenti Neo DEST $P_{CMVmin}$-blast$^R$-P2A-GFP-hPEST | Lentiviral |
| 47 | Cas9 | pLenti-EF1α-IRES-Zeo | Lentiviral |
| 48 | sgControl | lentiGuide-puro | Lentiviral |
| 49 | sgBCAT1 | lentiGuide-puro | Lentiviral |
| 50 | sgBCAT2 | lentiGuide-puro | Lentiviral |
| 51 | EV | pLenti-EF1α-IRES-Hygro | Lentiviral |
| 52 | BCAT1-FLAG | pLenti-EF1α-IRES-Hygro | Lentiviral |
| 53 | EV | pLenti-EF1α-Gateway-PGK-Hygro | Lentiviral |
| 54 | BCAT1-FLAG | pLenti-EF1α-Gateway-PGK-Hygro | Lentiviral |
| 55 | BCAT2-FLAG | pLenti-EF1α-Gateway-PGK-Hygro | Lentiviral |
| 56 | MTS-BCAT1-FLAG | pLenti-EF1α-Gateway-PGK-Hygro | Lentiviral |
| 57 | mCherry-T2A-VP64-NtcA-2xNLS$_{SV40}$ | pLenti-EF1α-Gateway-PGK-Hygro | Lentiviral |
| 58 | mCherry-T2A-VP64-NtcA$^{R90E}$-2xNLS$_{SV40}$ | pLenti-EF1α-Gateway-PGK-Hygro | Lentiviral |
| 59 | sgControl | lentiCRISPR_v2-puro | Lentiviral |
| 60 | sgBCAT1 | lentiCRISPR_v2-puro | Lentiviral |
| 61 | sgBCAT1 #1 | lentiCRISPR_v2-puro | Lentiviral |
| 62 | sgBCAT1 #3 | lentiCRISPR_v2-puro | Lentiviral |
| 63 | sgBCAT2 #1 | lentiCRISPR_v2-puro | Lentiviral |
| 64 | sgBCAT2 #2 | lentiCRISPR_v2-puro | Lentiviral |
| 65 | sgOGDH #1 | lentiCRISPR_v2-puro | Lentiviral |
| 66 | sgOGDH #2 | lentiCRISPR_v2-puro | Lentiviral |
| 67 | sgDLD #2 | lentiCRISPR_v2-puro | Lentiviral |
| 68 | sgDLD #3 | lentiCRISPR_v2-puro | Lentiviral |
| 69 | sgGPT2 #2 | lentiCRISPR_v2-puro<br>lentiCRISPR_v2-blast<br>lentiCRISPR_v2-mCherry | Lentiviral |
| 70 | sgGPT2 #3 | lentiCRISPR_v2-puro<br>lentiCRISPR_v2-blast<br>lentiCRISPR_v2-mCherry | Lentiviral |
| 71 | sgSLC25A11 #1 | lentiCRISPR_v2-puro<br>lentiCRISPR_v2-mCherry | Lentiviral |
| 72 | sgSLC25A11 #2 | lentiCRISPR_v2-puro<br>lentiCRISPR_v2-mCherry | Lentiviral |
| 73 | sgSNRPG #1 | lentiCRISPR_v2-puro | Lentiviral |
| 74 | sgSNRPG #2 | lentiCRISPR_v2-puro | Lentiviral |
| 75 | sgGLUD1 #1 | lentiCRISPR_v2-puro | Lentiviral |

| | | | |
|---|---|---|---|
| 76 | sgGLUD1 #2 | lentiCRISPR_v2-puro | Lentiviral |
| 77 | sgIDH3A #2 | lentiCRISPR_v2-puro | Lentiviral |
| 78 | sgIDH3A #5 | lentiCRISPR_v2-puro | Lentiviral |
| 79 | sgPDHA1 #2 | lentiCRISPR_v2-puro | Lentiviral |
| 80 | sgPDHA1 #3 | lentiCRISPR_v2-puro | Lentiviral |
| 81 | sgPDHA1 #4 | lentiCRISPR_v2-puro | Lentiviral |
| 82 | EV | pLenti PGK Blast DEST (w524-1) | Lentiviral |
| 83 | GPT2-HA (ENST00000340124.9) | pLenti PGK Blast DEST (w524-1) | Lentiviral |
| 84 | GPT2-HA (ENST00000440783.2) | pLenti PGK Blast DEST (w524-1) | Lentiviral |
| 85 | SLC25A11 | pLenti PGK Blast DEST (w524-1) | Lentiviral |
| 86 | *glnA* PCC 6803 5x sites with native spacers & sequence | pLenti Neo DEST $P_{CMVmin}$-GFP | Lentiviral |

**Table S4. List of CRISPR sgRNAs**

| sgRNA ID | Sequence | Source |
|---|---|---|
| sgControl | GGAGGCTAAGCGTCGCAA | PMID: 30220459 |
| sgBCAT1 | TATTAGGTCTTTAGCCTG | PMID: 30220459 |
| sgBCAT2 | TCCACTACTCCCTGCAGG | PMID: 30220459 |
| sgBCAT1 #1 | CAGAACCTGTCATTGCACCC | This Paper |
| sgBCAT1 #3 | GTTTCAGCCAAACCTCAACA | This Paper |
| sgBCAT2 #1 | ACGAACAGGAGCGCGCGCGT | This Paper |
| sgBCAT2 #2 | CAACAGGAGGCACTCAAGCG | This Paper |
| sgOGDH #1 | TTGGCCACTCATAGATACGA | This Paper |
| sgOGDH #2 | GACTAGTTCGAACTATGTGG | This Paper |
| sgDLD #2 | GGTGGAACATGCTTGAATGT | This Paper |
| sgDLD #3 | AATTCTTAGTAAAGGGTCGT | This Paper |
| sgGPT2 #2 | ATGCTAAGAAACGTGCCCGG | This Paper |
| sgGPT2 #3 | GCGGTGGAGTACGCCGTGCG | This Paper |
| sgSLC25A11 #1 | CGCCCTGATTCGAATCACCC | This Paper |
| sgSLC25A11 #2 | GCTGTTTGAGCGCCTGACTG | This Paper |
| sgSNRPG #1 | TTAATAGTGAAATTAAATGG | This Paper |
| sgSNRPG #2 | CATGTCCAAGGAATATTGCG | This Paper |
| sgGLUD1 #1 | CGCCCGGCGCCACTACAGCG | This Paper |
| sgGLUD1 #2 | TCAGTGCTGTAACGGATACC | This Paper |
| sgIDH3A #2 | CAACCACCGGAGCAACGTCA | This Paper |
| sgIDH3A #5 | GAGTATCAAGCTCATCACCG | This Paper |
| sgPDHA1 #2 | AGTAAAGCCGTGAGCCCGGT | This Paper |
| sgPDHA1 #3 | TATGCCAAGAACTTCTACGG | This Paper |
| sgPDHA1 #4 | AGCACTGATTACTACAAGAG | This Paper |

**Table S5. DNA plasmids used to express the αKG-ON biosensor system in each figure panel**

| Figure | Panel(s) | Plasmid IDs (from Table S3) |
|---|---|---|
| 1 | C | 1-16 |
| | D | 10, 17 |
| | E | 9, 18, 19 |
| 2 | A, E | 9 |
| | B, F | 29 |
| | C, G | 30 |
| | D, H | 31 |
| | I | 32-33 |
| | K, L | 32, 47-50 |
| | M | 32-33, 51-52, 59-60 |
| | N | 54-56 |
| | O | 32, 53-56 |
| 3 | A-C | 2, 47, 57-58 |
| | D, E | 2, 57-59, 61-62 |
| | F, G | 2, 57-59, 67-68 |
| | H, I | 2, 57-59, 71-72 |
| | J, K | 2, 57-59, 69-70 |
| 4 | B, C | 59, 69-72 |
| | D | 57-59, 71-72, 82-83, 86 |
| | E | 57-59, 67-68, 70, 86 |
| | F | 59, 67-70 |
| 5 | G | 59, 69-70 |
| | H | 2, 57-59, 69-70 |
| S1 | A, D | 10-16, 20 |
| | F | 10, 15. 21-23 |
| | G | 9-10, 15, 21 |
| | H | 10, 15, 24-28 |
| | I | 10, 15, 17, 28 |
| | J | 9-10, 15, 17, 28 |
| S2 | A, D | 9, 34 |
| | B, E | 9, 35 |
| | C, F | 9, 36 |
| | G, J | 9, 37 |
| | H, K | 9, 38 |
| | I, L | 9, 39 |
| | M, P | 9, 40 |
| | N, Q | 9, 41 |
| | O, R | 9, 42 |
| S3 | B, F | 43 |
| | C, G | 44 |
| | D, H | 45 |
| | E, I | 46 |

| | | |
|---|---|---|
| S4 | A, C, E, G | 29 |
| | B, D, F | 30 |
| S5 | A | 32, 47-50 |
| | B | 32, 51-52, 59-60 |
| | C | 33, 51-52, 59-60 |
| | D | 53-56 |
| S6 | A | 57, 59, 71-72, 82, 85-86 |
| | B | 57-59, 71-72, 82, 85-86 |
| | C | 57, 59, 69-70, 82-83, 86 |
| | D | 57-59, 69-70, 82-83, 86 |
| S7 | A | 2, 47, 58 |
| | B | 2, 47, 57-58 |
| | C | 2, 57, 59, 63-64 |
| | D | 2, 57-59, 63-64 |
| | E | 2, 57, 59, 75-76 |
| | F | 2, 57-59, 75-76 |
| | G | 2, 57, 59, 77-78 |
| | H | 2, 57-59, 77-78 |
| | I | 2, 57, 59, 65-66 |
| | J | 2, 57-59, 65-66 |
| S8 | A | 59, 61-72, 75-78 |
| | B | 2, 57-59, 61-72, 75-78 |
| | C | 59 |
| | D | 61-62 |
| | E | 67-68 |
| | F | 71-72 |
| | G | 69-70 |
| | H | 63-64 |
| | I | 75-76 |
| | J | 77-78 |
| | K | 65-66 |
| | L | 73-74 |
| | M | 2, 57, 59, 73-74 |
| | N | 2, 57-59, 73-74 |
| | O | 59, 73-74 |
| | P | 59, 67-72 |
| S9 | A | 2, 57, 59, 79-81 |
| | B | 2, 57-59, 79-81 |
| | C-E | 59, 69-72 |
| | F | 57, 59, 69-72, 86 |
| | G | 57-59, 69-72, 86 |

| | | |
|---|---|---|
| | H | 57, 59, 69-72, 86 |
| | I | 57-59, 69-72, 86 |
| S10 | E | 57, 82-84, 86 |
| | F | 83-84 |
| | G | 57-58, 82-84, 86 |
| S11 | A | 57, 59, 71-72, 82-83, 86 |
| | B | 57, 59, 67-70, 86 |
| | C | 57-59, 67-69, 86 |
| | D | 57, 59, 65-66, 70, 86 |
| | E | 57-59, 65-66, 70, 86 |
| | F | 59, 65-66, 69-70 |
| S12 | I | 59, 69-70 |

**References**


1. Q. Ouyang, T. Nakayama, O. Baytas, S. M. Davidson, C. Yang, M. Schmidt, S. B. Lizarraga, S. Mishra, M. Ei-Quessny, S. Niaz, M. Gul Butt, S. Imran Murtaza, A. Javed, H. R. Chaudhry, D. J. Vaughan, R. S. Hill, J. N. Partlow, S.-Y. Yoo, A.-T. N. Lam, R. Nasir, M. Al-Saffar, A. J. Barkovich, M. Schwede, S. Nagpal, A. Rajab, R. J. DeBerardinis, D. E. Housman, G. H. Mochida, E. M. Morrow, Mutations in mitochondrial enzyme GPT2 cause metabolic dysfunction and neurological disease with developmental and progressive features. *Proc Natl Acad Sci U S A* **113**, E5598-5607 (2016).

2. Y. Sonoda, T. Ozawa, Y. Hirose, K. D. Aldape, M. McMahon, M. S. Berger, R. O. Pieper, Formation of intracranial tumors by genetically modified human astrocytes defines four pathways critical in the development of human anaplastic astrocytoma. *Cancer Res* **61**, 4956–60 (2001).

3. D. Rohle, J. Popovici-Muller, N. Palaskas, S. Turcan, C. Grommes, C. Campos, J. Tsoi, O. Clark, B. Oldrini, E. Komisopoulou, K. Kunii, A. Pedraza, S. Schalm, L. Silverman, A. Miller, F. Wang, H. Yang, Y. Chen, A. Kernytsky, M. K. Rosenblum, W. Liu, S. A. Biller, S. M. Su, C. W. Brennan, T. A. Chan, T. G. Graeber, K. E. Yen, I. K. Mellinghoff, An Inhibitor of Mutant IDH1 Delays Growth and Promotes Differentiation of Glioma Cells. *Science* **340**, 626–30 (2013).

4. T. la Cour, L. Kiemer, A. Mølgaard, R. Gupta, K. Skriver, S. Brunak, Analysis and prediction of leucine-rich nuclear export signals. *Protein Eng Des Sel* **17**, 527–536 (2004).

5. W. Li, H. Xu, T. Xiao, L. Cong, M. I. Love, F. Zhang, R. A. Irizarry, J. S. Liu, M. Brown, X. S. Liu, MAGeCK enables robust identification of essential genes from genome-scale CRISPR/Cas9 knockout screens. *Genome Biol* **15**, 554 (2014).

6. “Geigy scientific tables. 1: Units of measurement, body fluids, composition of the body, nutrition” (Ciba-Geigy, Basel, 8., rev.enl. ed., 1981).

7. X. Su, W. Lu, J. D. Rabinowitz, Metabolite Spectral Accuracy on Orbitraps. *Anal. Chem.* **89**, 5940–5948 (2017).

8. S. K. McBrayer, J. R. Mayers, G. J. DiNatale, D. D. Shi, J. Khanal, A. A. Chakraborty, K. A. Sarosiek, K. J. Briggs, A. K. Robbins, T. Sewastianik, S. J. Shareef, B. A. Olenchock, S. J. Parker, K. Tateishi, J. B. Spinelli, M. Islam, M. C. Haigis, R. E. Looper, K. L. Ligon, B. E. Bernstein, R. D. Carrasco, D. P. Cahill, J. M. Asara, C. M. Metallo, N. H. Yennawar, M. G. Vander Heiden, W. G. Kaelin, Transaminase Inhibition by 2-Hydroxyglutarate Impairs Glutamate Biosynthesis and Redox Homeostasis in Glioma. *Cell* **175**, 101-116.e25 (2018).

9. H. Yoo, M. R. Antoniewicz, G. Stephanopoulos, J. K. Kelleher, Quantifying reductive carboxylation flux of glutamine to lipid in a brown adipocyte cell line. *J Biol Chem* **283**, 20621–7 (2008).

10. M. Pertea, D. Kim, G. M. Pertea, J. T. Leek, S. L. Salzberg, Transcript-level expression analysis of RNA-seq experiments with HISAT, StringTie and Ballgown. *Nat Protoc* **11**, 1650–1667 (2016).

11. P. Akan, A. Alexeyenko, P. I. Costea, L. Hedberg, B. W. Solnestam, S. Lundin, J. Hällman, E. Lundberg, M. Uhlén, J. Lundeberg, Comprehensive analysis of the genome transcriptome and proteome landscapes of three tumor cell lines. *Genome Med* **4**, 86 (2012).

12. J. F. de Sousa, P. da Silva, R. B. Serafim, R. P. Nociti, C. G. Moreira, W. A. Silva, V. Valente, RNA sequencing data of different grade astrocytoma cell lines. *Data Brief* **34**, 106643 (2021).
